# Multiscale regulation of nutrient stress responses in *Escherichia coli* from chromatin structure to small regulatory RNAs

**DOI:** 10.1101/2024.06.20.599902

**Authors:** Alyssa M. Ekdahl, Tatiana Julien, Sahana Suraj, Judith Kribelbauer, Saeed Tavazoie, P. Lydia Freddolino, Lydia M. Contreras

## Abstract

Recent research has indicated the presence of heterochromatin-like regions of extended protein occupancy and transcriptional silencing of bacterial genomes. We utilized an integrative approach to track chromatin structure and transcription in *E. coli* K-12 across a wide range of nutrient conditions. In the process, we identified multiple loci which act similarly to facultative heterochromatin in eukaryotes, normally silenced but permitting expression of genes under specific conditions. We also found a strong enrichment of small regulatory RNAs (sRNAs) among the set of differentially expressed transcripts during nutrient stress. Using a newly developed bioinformatic pipeline, the transcription factors regulating sRNA expression were bioinformatically predicted, with experimental follow-up revealing novel relationships for 36 sRNA-transcription factors candidates. Direct regulation of sRNA expression was confirmed by mutational analysis for five sRNAs of metabolic interest: IsrB, CsrB and CsrC, GcvB, and GadY. Our integrative analysis thus reveals additional layers of complexity in the nutrient stress response in *E. coli* and provides a framework for revealing similar poorly understood regulatory logic in other organisms.

## INTRODUCTION

Bacteria tailor their physiology to changing nutrient availability in their environments throughout their lifecycle. For example, enteric *Escherichia coli* must survive in the mammalian intestinal tract, which depending on location can vary substantially in nutrient composition (carbon, amino acids, oxygen availability) (1). As nutrients become scarce, bacteria cells use the general stress response to switch to a more stagnant growth referred to as stationary phase (2, 3). The central regulator of the general stress response, including the response to carbon source starvation, is the alternative sigma factor, RpoS. RpoS binds to RNA polymerase (RNAP) to facilitate transcription of hundreds of genes (3). Alternatively, when starved of amino acids or nitrogen, the stringent response is activated to direct RNAP to essential processes via ppGpp signaling (4–6). Among these global metabolic responses, there are other regulatory networks that manage stresses coinciding with nutrient deprivation, like oxidation, anaerobiosis, pH shifts, among others (2, 7).

At the DNA level, transcriptional regulation of genes is tuned by DNA binding proteins, including transcription factors, sigma factors, and nucleoid-associated proteins. Locally, transcription factors can bind near promoters to alter transcription of neighboring genes by facilitating or blocking sigma factors and RNAP from initiating transcription; sigma factors themselves play a major role in guiding RNA polymerase to specific subsets of promoters. While transcription factors are often associated with a singular stress, many regulate expression during the bacteria’s lifecycle due to coinciding stresses that occur with fluctuating nutrients (2). Globally, nucleoid-associated proteins can bind long segments of the genome to bend and compact the DNA in response to changing environments (8). Like eukaryotic heterochromatin, these elongated protein occupancy domains, referred to as EPODs (9), can condense long sections of the DNA and silence transcription by blocking the accessibility of RNAP to internal promoters (10–13) or aiding in RNAP stalling and backtracking (14, 15). Nucleoid-associated proteins often respond to growth and nutrient variability, typified in *E. coli* by proteins such as the amino acid regulator Lrp (6), and the growth phase dependent HU and Dps (16). Taken together, protein-directed transcriptional regulation relies on dynamic coordination at a global and local level to respond to changing environments.

At the RNA level, post-transcriptional regulation of RNAs is tuned by RNA binding proteins (17, 18) and small regulatory RNAs (sRNAs) (19–22). These global regulators quickly control translation by binding mRNAs to toggle the ribosomal binding site accessibility and transcript stability (23, 24). Transcriptional and post-transcriptional regulation are highly interdependent (25–28). For instance, the mRNA encoding the general stress response regulator, *rpoS*, is regulated by five sRNAs [ArcZ (29), DsrA (30), OxyS (31), CyaR (32), RprA (33)] that, upon base-pairing, repress or activate *rpoS* translation. Additionally, a multitude of DNA binding proteins regulate sRNA expression to complement their dynamic functionalities; as an example, at least four transcription factors (OmpR (34), MarA, SoxS, and Rob (35)) and two nucleoid-associated proteins (Lrp (36) and IHF (37)) directly regulate expression of the sRNA MicF (38). Yet, much remains unknown about the interplay between transcriptional and post-transcriptional regulators, including in response to nutrient stress. Transcriptional regulation of sRNAs have largely been understudied, with the majority of sRNAs in *E. coli* lacking even one known transcription factor regulating them (24). Additionally, the regulation of sRNAs by DNA condensation through nucleoid-associated proteins has not been globally investigated, although intragenic regions (which often encode sRNAs) have previously been found to be enriched in nucleoid-associated protein binding (9, 11, 13). We thus sought to uncover the protein-directed regulation of sRNAs during nutrient stress, both during glucose and amino acid starvation.

The complexity of the regulatory responses noted above – which span transcriptional regulation at the levels of local regulators, global regulators and chromatin structure, to additional interwoven mechanisms of post-transcriptional and even post-translational regulation – necessitates the integration of data at many regulatory layers to unravel the logic driving cellular regulatory responses. In this study, we explore the global and local protein-directed transcriptional regulation of the nutrient stress response over growth of *E. coli* when starved of glucose or amino acids. Samples were analyzed by RNA-seq, RNAP chromatin immunoprecipitation (ChIP), and *in vivo* protein occupancy display – high resolution (IPOD-HR) (10) to reveal the dynamic gene regulatory networks at the DNA and RNA level. IPOD-HR is particularly well suited for interrogating complex regulatory responses, because it permits both unbiased identification of transcription factor binding sites (without the need for *a priori* knowledge of which factors should be targeted), and measurement of heterochromatin-like EPODs. Our results revealed the presence of many EPODs which were static across growth conditions (thus effectively behaving like constitutive heterochromatin), and others which were dynamic and opened to allow gene expression in some cases, thus behaving like facultative heterochromatin in eukaryotes. Our transcriptomic data also highlight sRNAs as one of the most substantially changing classes across nutrient stress conditions. Notably, the DNA regions encoding sRNAs were predominantly outside of EPODs, suggesting the nutrient stress dynamics of sRNA expression are largely due to complex transcription factor regulation. We go further to bioinformatically predict and experimentally validate novel transcription factors of sRNAs. Mutational analyses confirmed novel direct regulation for five sRNAs important for metabolic regulation (IsrB, CsrB, CsrC, GcvB, and GadY), while supporting another 36 sRNA-transcription factor relationships.

## MATERIAL AND METHODS

### Bacterial strains and media

*Escherichia coli* K-12 MG1655 and its transcription factor knockout derivatives, as listed in Table S1, were either previously created by the Keio Collection (39) or, in the cases where the particular Keio strain was unable to be cured or confirmed (Δ*crp*, Δ*oxyR,* and Δ*ulaR*), were gifted by the Wade Lab (University of Albany, Albany, NY) in the K-12 MG1655 background. In both cases, the transcription factor genes were knocked out by P1 transduction of the FRT-flanked *kanR* cassette. Curing of the marker was performed by FLP recombinase using the pCP20 plasmid (40). Curing was checked by selective plates to ensure loss of pCP20 plasmid and kanamycin resistance. All gene deletions were confirmed by PCR fragment size surrounding the scar site before and after curing. Genomic *lacZ* reporters are described in *Transcriptional Reporter Construction*.

Strains were grown from cryogenically preserved glycerol stocks and streaked onto Luria-Bertani, LB (10 g/l tryptone, 5 g/l yeast extract, 5 g/l NaCl, 15g/l agar) solid media plates and grown overnight at 37°C. Single colonies were picked from plates and grown overnight at 37°C in liquid media to saturation. Solid media plates and liquid media contained 100 μg/ml Carbenicillin when applicable for plasmid retention. LB (10 g/l tryptone, 5 g/l yeast extract, 5 g/l NaCl) or SOB (5 g/l yeast extract, 20 g/l tryptone, 0.5 g/l NaCl, 0.186 g/l KCl, 2.4 g/l MgSO_4_ adjusted to pH 7.5) media was used for all cloning and general growth conditions.

For physiological experiments, the chemically-defined media used (10) are supplemented versions of M9 base (6 g/L Na_2_HPO_4_, 3 g/L KH_2_PO_4_, 1 g/L NH_4_Cl, 0.5 g/L NaCl, 1 mM MgSO_4_) (41). The glucose minimal medium, Glu, contains 0.2% (w/v) glucose, MOPS micronutrients (42), 2 mM MgSO_4_, 0.1 mM CaCl_2_, and 2 μM ferric citrate. The casamino acid minimal medium, Cas, contains 1% (w/v) casamino acids, MOPS micronutrients (42), 0.1 mM CaCl2, and 2 μM ferric citrate. For the Miller assays, the Glu media was slightly modified to 1mM MgSO_4_, 0.4 mM CaCl_2_ and 40 μM ferric citrate to mimic previous IPOD-HR studies (10). The M9 rich-defined medium, RDM, as previously used (1) is supplemented with 0.4% (w/v) glucose, MOPS micronutrients (42), 1mM MgSO_4_, 4 μM CaCl_2_, 40 μM ferric citrate, and 1X supplement ACGU and EZ used in MOPS EZ media (Teknova).

### Sample collection for IPOD-HR

Sample collection for IPOD-HR was performed largely as previously described (10). *E. coli* K-12 MG1655 cells were grown overnight in the respective media (either Glu or Cas). For the Glu medium samples, 1 mL of overnight sample was seeded into 200 mL fresh Glu medium and grown shaking at 37°C to the different time points: 6 hours for mid-logarithmic (OD_600_ ∼0.25), 10 hours for stationary (OD_600_ ∼1.7) and 18 hours for late stationary. For the Cas medium samples, 134 μL of overnight sample was seeded into 200 mL fresh Cas medium and grown shaking at 37°C to the different time points: 2.5 hours for early stationary (OD_600_ ∼0.1), 4.5 hours for mid-logarithmic (OD_600_ ∼0.5), 6.33 hours for late logarithmic (OD_600_ ∼1.4), 10 hours for stationary (OD_600_ ∼4), and 18 hours for late stationary (OD_600_ ∼5.5).

Samples were treated with 150 μg/mL rifampicin for 10 minutes at 37°C with continuous shaking to stall RNA polymerase at active promoters and allow transcriptional runoff for already-elongating polymerase. Formaldehyde to 1% final concentration was added for crosslinking and incubated at room temperature for 5 minutes shaking. Excess glycine was then added to quench the crosslinking reaction (final concentration 0.333M) for 5 minutes at room temperature, with shaking. Cultures were then chilled on ice and washed twice using phosphate-buffered saline. Final cross-linked cell pellets were stored at -80°C until ready for extraction.

### Cell lysis and DNA/RNA preparation

Detailed instructions for IPOD-HR interface extraction, RNA polymerase chromatin immunoprecipitation, and crosslinking reversal and recovery of DNA have been previously published (10), and were followed precisely here. Illumina sequencing libraries were prepared using a NebNext Ultra DNA kit, following the manufacturer’s instructions except that a Qiaquick Nucleotide Removal Kit was used for pre-ligation cleanup to avoid loss of low molecular weight fragments. RNA-seq samples were prepared using an Epicentre ScriptSeq v2 kit following the manufacturer’s instructions.

### DNA sequencing and analysis

The IPOD and RNAP ChIP DNA sequencing and analysis were performed as previously described and validated (10). All reads were aligned to the *E. coli* K-12 MG1655 reference genome (U00096.3).

### Extended Protein Occupancy Domains (EPODs) calling

EPOD calling was performed as previously described (10, 11). Briefly, we calculated 512 bp and 256 bp rolling means across the genome using the IPOD-HR rz-scores. Strict EPODs threshold score was set at the 90^th^ percentile, while loose threshold score was set to 75^th^ percentile from the 256 bp rolling mean signal. Regions at least 1, 024 bp long with median occupancy score above the threshold score were considered a loose or strict EPOD, depending on the maximum threshold passed by their occupancy. These regions were then expanded in both directions on the genome until the median occupancy score dropped below the threshold or the occupancy rz-score dropped below zero. EPODs were further classified as static by comparing boundaries of called loose EPODs using bedtools (43) intersect function, and further filtered to overlapping regions that achieve the strict threshold in at least one of the compared conditions.

### RNA sequencing and differential expression analysis

RNA-seq datasets were quality checked by FastQC, then aligned to the *E. coli* K-12 MG1655 genome (GenBank U00096.3) using the Burrows-Wheeler Alligner tool (BWA-MEM) in Samtools (44). Gene counts were then called using HTSeq (45) with a modified gene annotation file containing all sRNAs of interest that are not listed in U00096.3 (coordinates provided in Table S1). Differential expression was performed by DESeq2 (46) using the LRT test with log_2_FC shrink method “ashr”(47) between each neighboring growth phase condition (*i.e.*, early logarithmic versus mid logarithmic, mid logarithmic versus late logarithmic, etc.) and between similar growth condition between media (*i.e.,* mid logarithmic in Glu versus Cas, etc.). All other growth phase combinations were also compared for differential expression to compare to the maximum to minimum protein occupancy growth phases for sRNA peak calling (described in *Prediction of Transcription Factors for sRNA Promoters*).

### Gene Ontology (GO) analysis

All genes overlapping a strict EPOD were input into the David webtool (48, 49) for functional annotation analysis. GO analysis was also performed on static EPODs in Cas and Glu media. Alternatively, for the RNA-seq analyses, all genes differentially expressed (|log_2_FC|>1, p-adj < 0.05) for each condition were input into the DAVID webtool for functional annotation analysis. Enrichment of Gene Ontology (GO) biological processes terms was defined as terms with EASE scores (50), a modified Fisher Exact p-value, of <0.05. The enriched GO terms were summarized using select terms for comparison between conditions.

### Prediction of transcription factors for sRNA promoters

Adjusted z-scores, using the *p*-value for enrichment at each site (log_10_ *p*-values are calculated under the null hypothesis that the distribution of the robust z-scores is standard normal) was used for peak calling across the genome. Peak calling was performed using the continuous wavelet transform (CWT) peak calling in the scripy.signal package, as previously done (10, 51). Simply, the peak width of 60 bp was used over a range of signal-to-noise thresholds, with the maximum threshold at which the peak call is found defines the protein occupancy for downstream analyses. Peaks were then filtered for occurring near sRNAs ([-200, +50] bp surrounding the sRNA encoded region). This adjustment was selected based on prior success with sRNA promoters (51).

Peaks near sRNAs were merged between the 8 conditions, in which overlaps of 50 bp or greater was considered the same binding site. The maximum threshold in each condition was collected for each merged peak, with a minimum threshold of 1 in at least one condition required to be considered a peak. The thresholds were tracked over growth for each media. The growth phase(s) of maximum to minimum protein occupancy was identified and compared to the RNA-seq differential expression conditions. If significant differential expression of the respective sRNA occurred over this growth timeframe, that region is identified as a likely transcription-altering protein binding region.

All identified regions were put through the motif searching algorithm, FIMO (52), using three separate motif databases as previously curated (51): dpInteract (53), SwissRegulon (54), and Prodoric2 (55). Found motifs within transcription-altering protein binding regions were compared to external RNA-seq database results, as analyzed and compiled previously (51). Motifs were filtered by the external differential expression results (*i.e.*, sRNA was found to be differentially expressed in a condition known to activate or repress the activity of the predicted transcription factor or nucleoid-associated proteins). These motifs that were predicted to bind within the [-100, 10] bp surrounding the sRNA transcription start site were collected for follow-up transcriptional reporter testing.

### Transcriptional reporter construction

Transcriptional reporters were created in high-copy plasmids using pNM-12, a pBAD24 derivative, as a foundation (30). pNM-12 was modified by first introducing a strong bidirectional terminator (5’-actaTACCACCGTCAAAAAAAACGGCGCTTTTTAGCGCCGTTTTTATTTTTCAACCTTaagctt-3’) just downstream of the pBAD promoter to prevent leaky expression. Each reporter then contains, downstream of the bidirectional terminator, the respective sRNA promoter (200nt upstream to through the RegulonDB annotated transcription start site by 10nt) followed by a strong synthetic ribosomal binding site [R148K (56)] and ß-galactosidase encoding *lacZ* gene. An AmpR cassette was used for antibiotic selection. Plasmids were cloned in house using Gibson Assembly with the NEB Hifi Builder Master Mix (New England Biolabs, Ipswich, MA) and/or contracted (GenScript, Piscataway, NJ, USA), and confirmed via Sanger Sequencing (Eton Biosciences, San Diego, CA) and/or full plasmid sequencing (Plasmidsaurus, Eugene, OR). Of note, the Spf (or Spot42) promoter was unable to be cloned without deleterious point deletions in the *lacZ* gene. A list of sRNA promoter sequences and primers used are included in Table S1.

For genomic Miller assays, the transcriptional reporters were amplified by PCR from the plasmid, keeping ∼1, 000bp homology on either side of the amplicon with the ara operon. The Keio collection (BW25113 strain) has part of the ara operon deleted (starting after the pBAD promoter to 8 amino acids into the *araD* gene). The homologous arms therefore contain part of the *araC* gene through pBAD promoter, as well as the last 15 amino acids of the *araD* gene and downstream. As noted in *Bacterial Strains and Media* section, some transcription factor knockout strains from the Keio collection were unable to be confirmed. Therefore, the Δ*oxyR* strain in K-12 MG1655 background was used and compared to the K-12 MG1655 wildtype strain for genomically testing regulation of GcvB. The K-12 MG1655 strain does not have a modified *ara* operon, so the genomic alterations in the wildtype and associated Δ*oxyR* strain included also removing the *araA* and *araB* genes (making the resulting insertion equivalent between strains).

Amplicons were inserted into the Keio BW25113 or K-12 MG1655 wildtype strains (as noted above) with respective transcription factor knockout strains using a CRISPR-recombineering lambda-red system, graciously shared by the Pfleger Lab (University of Wisconsin, Madison, WI) (57). Correct substitution of the transcriptional reporter for the truncated *araD* gene was confirmed via size by PCR and sequence confirmed by Sanger Sequencing (Eton Biosciences, San Diego, CA).

### Plasmid-based screening by Miller assays

For each proposed transcription factor for a given sRNA, the sRNA transcriptional reporter was transformed in both the transcription factor knockout strain and associated wildtype strain using the Keio collection (BW25113) with few aforementioned exceptions using K-12 MG1655 strain deletions (Δ*crp*, Δ*oxyR,* Δ*ulaR*). To account for strain differences in RNA polymerase availability, a negative control plasmid containing the Anderson promoter J23110, in place of the sRNA promoter, was used.

Plasmid copy number cannot be controlled and will deviate when the cell burden is high, often resulting in stunted growth. To account for this, cell growth was monitored by OD_600_ and no particular test cases showed stunted growth outside of strain issues (*i.e.*, slow growing strains such as Δ*crp* in Glu and RDM media, Δ*fur* and Δ*oxyR* in Glu medium, Δ*cra* and Δ*ompR* in Cas medium). Of note, Δ*ompR* failed to grow in the Glu medium and Δ*crp* failed to grow in the Cas medium, and thus sRNA promoters were unable to be tested in these conditions.

### Genomic Miller assays of mutated sRNA promoters

The cases found to be statistically significant in the plasmid-based Miller assays were followed up via genomic insertion of the wildtype transcriptional reporter as well as a rationally mutated reporter into the wildtype BW25113 strain at the *ara* operon (except for testing GcvB regulation by OxyR which was constructed in the K-12 MG1655 wildtype and respective *ΔoxyR* strains as explained in the *Transcriptional Reporter Construction* section). The mutated reporters consisted of mutating select nucleotides of high frequency to low frequency for the found transcription factor consensus motif. Using FIMO default settings (52), wildtype and mutant sRNA promoters were analyzed with the 3 motif databases as previously described, minimizing differences between predicted motifs other than the desired transcription factor consensus motif.

### Miller assays

Kinetic Miller assays were performed as previously documented in detail (10). Briefly, for the plasmid-based Miller assays, cell colonies containing the transcriptional reporters were plucked from LB plates containing 100μg/ml carbenicillin in biological triplicate and grown to saturation in the respective media. For the genomic mutant reporters, cell colonies containing the transcriptional reporters were plucked from LB plates in biological quadruplicates and grown to saturation in their respective liquid media. Cultures were then seeded 1:100 into fresh media, and samples were taken at mid logarithmic (RDM: 2hrs, Cas: 3hr, Glu: 4hr, OD 1cm path length equivalent ∼0.5-0.7) and stationary phase (RDM: 5hrs, Cas and Glu: 9hrs, OD 1cm path length equivalent ∼2-3). A few noteworthy strains were particularly slow growing and thus required longer growth times to achieve the stationary growth phase: Δ*crp* in RDM medium were sampled at 8hrs, Δ*cra* and Δ*ompR* in Cas medium and Δ*crp,* Δ*fur*, and Δ*oxyR* in Glu medium were sampled at 12hrs.

A 96-well plate was loaded with 200 μl of each sample for OD_600_ measurement. In separate wells, cell culture (80 μl for mid logarithmic, 10 μl with 70 μl blank media for stationary) was mixed with 120 μl of Miller assay reagent as previously described (10) (8 mL Z-buffer [60 mM Na_2_HPO_4_, 40 mM NaH_2_PO_4_, 10 mM KCl, 1 mM MgSO_4_], 21.6 μl β-mercaptoethanol, 3 mL ONPG solution [4 mg/ml ONPG in Z-buffer], 800 μl PopCulture Reagent (Sigma-Aldrich), 200 μl Lysozyme solution from chicken egg white (Sigma-Aldrich)).

Measurements of the β-galactosidase reaction were taken every 2 minutes for 1 hour on a BioTek Cytation 3 Plate Reader held at 30°C with moderate shaking between measurements. Specifically, the absorbances for β -galactosidase activity (via ONPG to ONP conversion, A_420_) and cell debris correction (1.7*A_550_) were measured over time to enable the kinetic Miller assay calculation as previously documented (10) and illustrated below, using a LOESS fit to calculate the maximum slope:

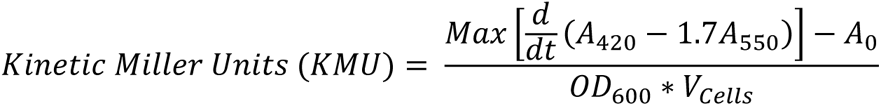

where V_cells_ is the volume of cells used in the reaction, and A_0_ is the average slope of the enzymatic activity (A_420_-1.7*A_550_) of the blank wells at the time.

For the plasmid-based Miller assays, Kinetic Miller Units (KMUs) between the wildtype and transcription factor knock outs for each sRNA promoter condition were statistically compared by two-tail independent t-test. Statistical significance was determined to be p-value < 0.05. Ratios of wildtype to transcription factor knock outs were baselined to the wildtype to transcription factor knockout ratios of the negative control reporter, containing the Anderson J23110 promoter in place of the sRNA promoter, to compensate for RNAP activity and prevalence between strains. This modified equation is shown below:

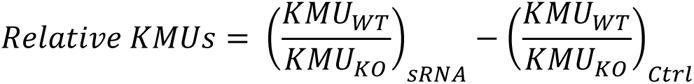

For the genomic mutant reporters, KMUs between the wildtype and mutant sRNA promoters inserted in the wildtype strain (either BW25113 or MG1655), as well as comparisons between wildtype and transcription factor knockout strains, were statistically compared by two-tail independent t-test. Statistical significance was determined to be p-value < 0.05.

## RESULTS

### Transcriptome and DNA protein occupancy indicate nutrient stress response coincides with other activated stress networks across growth

We investigated the global regulatory strategies used by *E. coli* to cope with a variety of nutrient stresses by growing K-12 wildtype cells in liquid cultures of two differing minimal media and sampling them throughout growth, as illustrated in **Figure 1A** (strain details provided in **Table S1**). Two M9 minimal media were used, either supplemented with 0.2% glucose (referred to as Glu) or 0.4% casamino acids (referred to as Cas), to observe nutrient stress differences between carbon sources (*i.e.*, amino acid starvation versus glucose starvation). Samples were taken at different growth stages to observe the coinciding general stress and nutrient stress responses that typically escalate through growth as carbon sources diminish: in Cas medium early, mid, and late logarithmic (due to the presence of multiple apparent auxic shifts likely due to exhaustion of different amino acids from the media), as well as stationary and late stationary growth stages; in Glu medium, mid logarithmic, stationary, and late stationary growth stages, with approximate OD_600_ values provided in **Figure 1A**. We analyzed each sample using RNA-seq, RNAP ChIP, and IPOD-HR to investigate the dynamics of protein-directed transcriptional regulation. Specifically, these three methods together provide genomic locations of DNA-bound proteins, RNA polymerase bound promoters (as samples were treated with Rifampicin to stall RNAP elongation) and resulting transcriptome at each sample condition.

**Figure 1.**
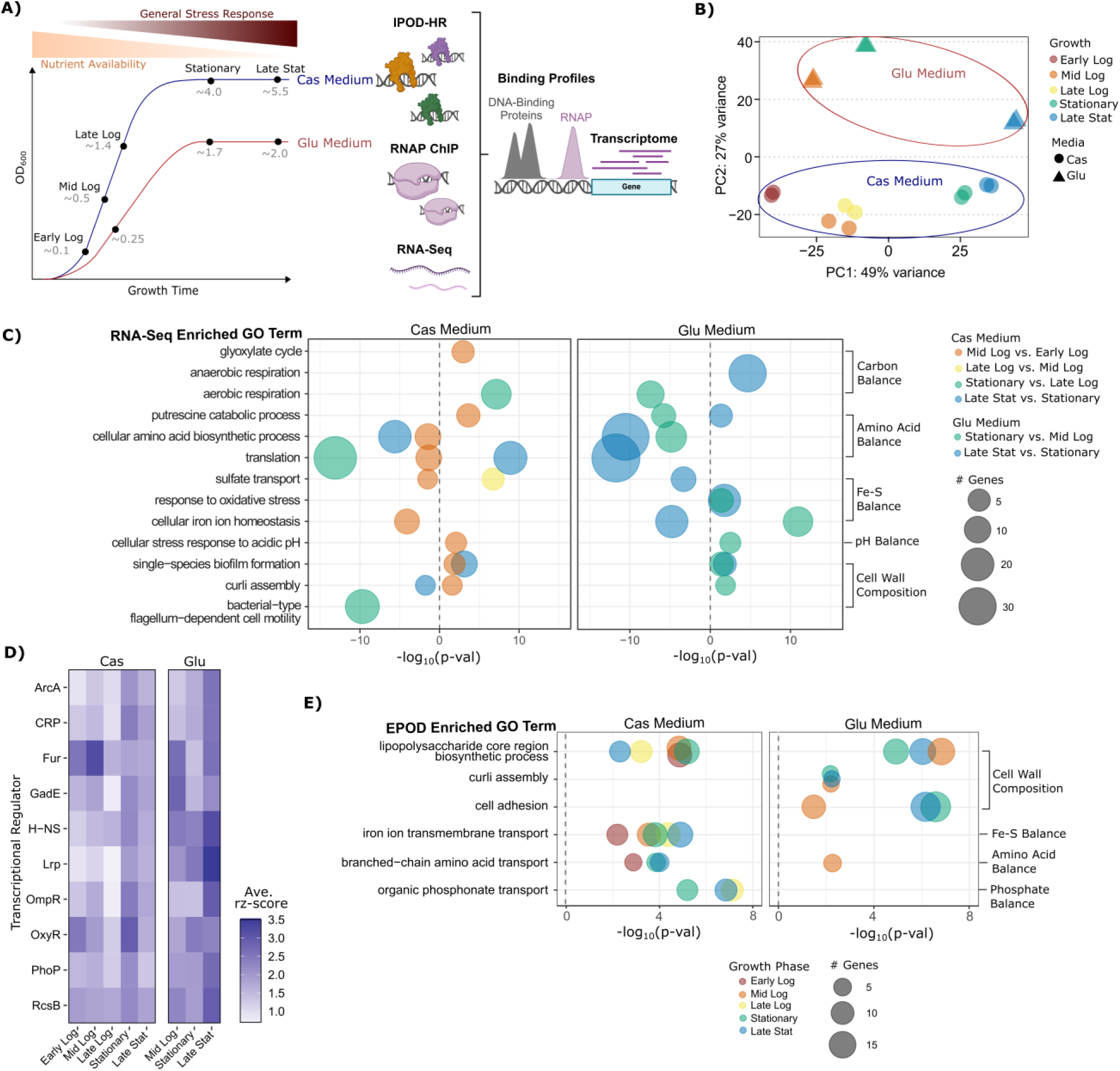
DNA protein occupancy and transcriptome dynamics capture nutrient stress response along with coinciding stress networks across growth. **A)** Illustration of the sample time points in both media conditions in this study (black dots) as defined by the growth curve (approximate 1cm pathlength OD_600_ for each sample provided in grey). Above the growth curves is a depiction of the expected reduction of nutrients across growth, alongside increased general stress response upon entry to stationary phase. The three types of analyses performed on the samples are illustrated to the right: *in vivo* protein occupancy display – high resolution (IPOD-HR), RNA polymerase chromatin immunoprecipitation (RNAP ChIP), and RNA-Seq. Coupled results enable the simultaneous measurements of DNA protein and RNAP binding profiles with resulting transcriptome. Created in part by BioRender.com. **B)** Principal component analysis (PCA) plot of the RNA-seq results for the 8 conditions tested in duplicate. Principal component (PC) 1, accounting for 49% of the variance, is associated with growth phase while PC2, accounting for 27% of the variance, is associated with media. **C)** Gene ontology analysis (using GO biological processes terms) of differentially expressed genes (p-adj <0.05, |log_2_FC| > 1) between each neighboring growth point in each media, represented by a bubble plot of select enriched GO terms for each media shown (Cas Media: left, Glu Media: right). The significance of GO term enrichment (by -log_10_ of the EASE score, a modified Fisher Exact p-value) is plotted on the x-axis (positive values indicate GO term enriched by genes with log_2_FC >1, while negative values indicate enrichment from genes with log_2_FC <1). Point color represents the condition in which the GO term was enriched, while point size represents the number of differentially expressed genes associated with the GO term. All significant GO terms for each compared condition are provided in Table S2. **D)** A heatmap displaying the average of the maximum IPOD-HR raw z-scores (rz-scores) within known DNA binding sites of each transcription factor (from RegulonDB) for each condition (all factors provided in Figure S2). Dynamic regulons show both growth dependence, such as ArcA and CRP that increase with growth in both media, and media dependence, such as GadE and OxyR that show different patterns with growth between media. **E)** Gene Ontology analysis of the genes within strict extended protein occupancy domains (EPODs) in all tested conditions, with select enriched GO terms presented. Point color represents the condition in which the GO term was enriched, while point size represents the number of genes associated with the GO term within the strict EPODs. Significance of enrichment provided as the -log_10_(EASE Score, a modified Fisher Exact p-value) on the x axis. All significant GO terms from the EPOD analysis are provided in Table S3.

As shown in **Figure 1B**, we found using principal component analysis (PCA) on the gene expression profiles that growth phase and media account for most transcriptome differences between our conditions (PC1 with 49% variance, PC2 with 27% variance). To compare transcriptomes between conditions, differential expression was determined between neighboring growth stages (*e.g.*, mid logarithmic versus early logarithmic) for each medium, and between media at similar growth stages (*e.g.*, mid logarithmic in Cas medium versus mid logarithmic in Glu medium) (volcano plots provided in **Figure S1,** values in **Table S2**). We observed distinct metabolic shifts present between each of our chosen growth points (**Figure 1C**) using the DAVID functional annotation tool to assess enriched gene ontology (GO) terms (48, 49) for all differentially expressed genes (p-adj <0.05, |log_2_FC| > 1). As expected, given the differing carbon sources (glucose or casamino acids), the enriched GO terms associated with carbon metabolism networks differed between media. For example, the Cas medium induced genes associated with general aerobic metabolism, while Glu medium relied on genes associated with anaerobic respiration at late growth stages. Both media types cause repression of genes associated with the amino acid biosynthesis and translation GO terms during stationary phase, aligned with prior studies of nutrient deprivation (58, 59).

To understand the global regulatory network accounting for the transcriptional responses that we identified in our growth conditions, without bias toward only well-characterized regulators, we measured global protein occupancy across the genome using IPOD-HR as previously described (10). Briefly, protein-DNA complexes were crosslinked and isolated, and then DNA bound to protein were sequenced. Quantification of aligned sequences by genomic position were normalized by input material and RNAP ChIP signal to generate a raw z-score (rz-score), a quantification for amount of protein excluding RNAP bound at a genomic position.

Transcription factor regulon activities were investigated by averaging the maximum rz-scores within known DNA binding sites as annotated by RegulonDB (60). Results for some representative regulators are provided in **Figure 1D** (all regulators in **Figure S2**). Dynamic occupancies highlight the activated regulatory networks with growth as well as between media conditions. For example, occupancy at binding sites for the secondary carbon source assimilation regulator, CRP, and microaerobic regulator, ArcA, both increase DNA occupancy (as indicated by increasing average rz-scores) through growth regardless of media condition. As carbon sources decrease through growth, CRP and ArcA are expected to bind DNA to shift carbon metabolism to less preferred pathways alongside the general stress response.

Less intuitive transcriptome results also align with the transcription factor activities. For example, the iron regulator, Fur, and oxidative stress regulator, OxyR, show high protein occupancy early in growth and are likely regulating the genes involved in iron acquisition and oxidative stress response as found by the Gene Ontology analysis (**Figure 1C**). Additionally, the acid regulator, GadE, with higher occupancy early in growth, is likely contributing to the enrichment of the GO term cellular stress response to acidic pH. However, the dynamic outer membrane regulator OmpR is quite contrasting to the stagnant behavior of the biofilm regulator, RcsB. These different binding dynamics of envelope stress regulators are likely contributing to the varying transcriptome behavior as represented by the enriched GO terms related to cell wall composition (e.g., biofilm, curli, and flagellum regulation). Altogether, the transcriptome and protein occupancy analyses capture the glucose and amino acid starvation responses throughout growth, while also emphasizing other stress networks activated alongside nutrient stress, such as low pH, envelope, iron, and oxidative stresses.

### Extended Protein Occupancy Domains (EPODs) indicate higher DNA condensation during amino acid limitation

From our protein regulon analysis (**Figure 1D**), we noted that H-NS and Lrp, along with other nucleoid-associated proteins, have generally higher occupancy in the Glu medium compared to the Cas medium. This observation is extended to the IPOD-HR data globally, where a general trend of higher protein binding (rz-scores) is found in dense regions of the genome in Glu medium compared to the same locations in Cas medium. The higher nucleoid-associated protein occupancy in Glu over Cas medium suggests potential differences in global DNA condensation between nutrient conditions. Indeed, we have previously observed heterochromatin-like regions of high protein occupancy, which we have termed extended protein occupancy domains (EPODs). EPODs can be formed by large-scale regions of nucleoid-associated protein occupancy and can participate in condition-dependent regulation of metabolic genes (10, 11). We found that in the present set of media conditions, the number of EPODs varies between conditions, ranging from 146 in Cas mid logarithmic phase, up to 198 in Glu late stationary phase, consistent with increased heterochromatinization under poorer media conditions (**Table S3**; unless otherwise noted, we use the “strict” definition of EPODs following (10): DNA regions of at least 1, 024 bp long with median protein occupancy greater than the 90^th^ percentile threshold of the 256 bp rolling mean).

As depicted in **Figure 1E**, a few enriched GO terms are shared in both media including lipopolysaccharide synthesis and branched chain amino acid transport. However, other enriched GO terms for EPODs differ between media; for example, iron and phosphonate transport that are enriched only in the Cas medium, while cell adhesion and curli formation are enriched only in the Glu medium. Note the EPOD enriched GO terms partially align with the prior discussed transcriptome enriched GO terms (**Figure 1C**), involving iron homeostasis, cell wall composition, and amino acid balance, suggesting contributions of chromatin structure and nucleoid-associated protein occupancy to transcriptional regulation during nutrient stress, alongside the more classically considered roles of local regulators.

### Static and dynamic EPODs rely on nucleoid-associated proteins and transcription factors to regulate expression

To map the contributions of chromatin structure to gene regulation across conditions in our study, we further classified EPODs as either static or dynamic. Static EPODs denote extended protein occupancy domains that are present at all growth stages in a particular medium, while dynamic EPODs refer to those extended protein occupancy domains that are present in only some growth stages in a particular medium. When comparing between media compositions, the number of static EPODs are strikingly different, with Glu medium maintaining 233 EPOD regions throughout growth while Cas only maintains 160 (**Figure S3**), again supporting the prior observation of higher overall DNA condensation during amino acid starvation and suggesting more substantial metabolic shifts across growth in the presence of the more complex set of carbon sources in Cas medium (*i.e.,* casamino acids supplied as the carbon source versus glucose in the Glu media). Gene ontology analysis of the genes contained within static EPODs indicate the enrichment of lipopolysaccharide biosynthesis genes in silenced regions in both media types, phosphonate and iron homeostasis in Cas medium (such as the iron regulatory *fecIRABCDE* operon in **Figure S4**), and cell wall composition pathways for Glu medium (such as the cryptic flagellum *yehEDCBA* operon in **Figure S5** and curli assembly *csgDEFG* and *csgBAC* operons in **Figure S6**). As illustrated in the following examples, static EPODs appear to predominantly reduce transcription of internal promoters as previously proposed (11, 61), while dynamic EPODs follow a more complex model requiring coordination between nucleoid-associated proteins and transcription factors to regulate transcription.

A clear example of a static EPOD, accounting for the GO term enrichment for lipopolysaccharide core region biosynthesis revealed by our analysis, is the *rfaDwaaFCL*, *rirA, waaQGPSBOJ* and *waaYZU* operons shown in **Figure 2A**. Protein occupancy is plotted over genomic coordinates at mid logarithmic (top), stationary (middle) and late stationary (bottom) phases in both Glu medium (grey) and Cas medium (green). Above each protein occupancy chart are bars representing the regions of EPODs in each media (grey: Glu, green: Cas) as well as known regulator binding sites (orange: activator, purple: repressor). As illustrated, the EPODs in both media remain relatively static over growth, however with the slight reduction of protein occupancy in the Cas medium at late stationary phase. Differential RNA expression of these genes, depicted as a heatmap in **Figure 2B**, compare expression in Cas versus Glu media at the same three growth stages as the plotted protein occupancies. When the EPOD behavior is similar between media (*i.e.*, mid logarithmic and stationary growth phases), RNA expression is similar as indicated by non-significant differential expression. Aligned with prior documented EPOD functionality of silencing transcription (11, 61), as the EPOD diminishes in late stationary phase in Cas medium, the RNA expression for the *rirA, waaQGPSBOJ* and *waaYZU* operons increases in Cas compared to Glu media.

**Figure 2.**
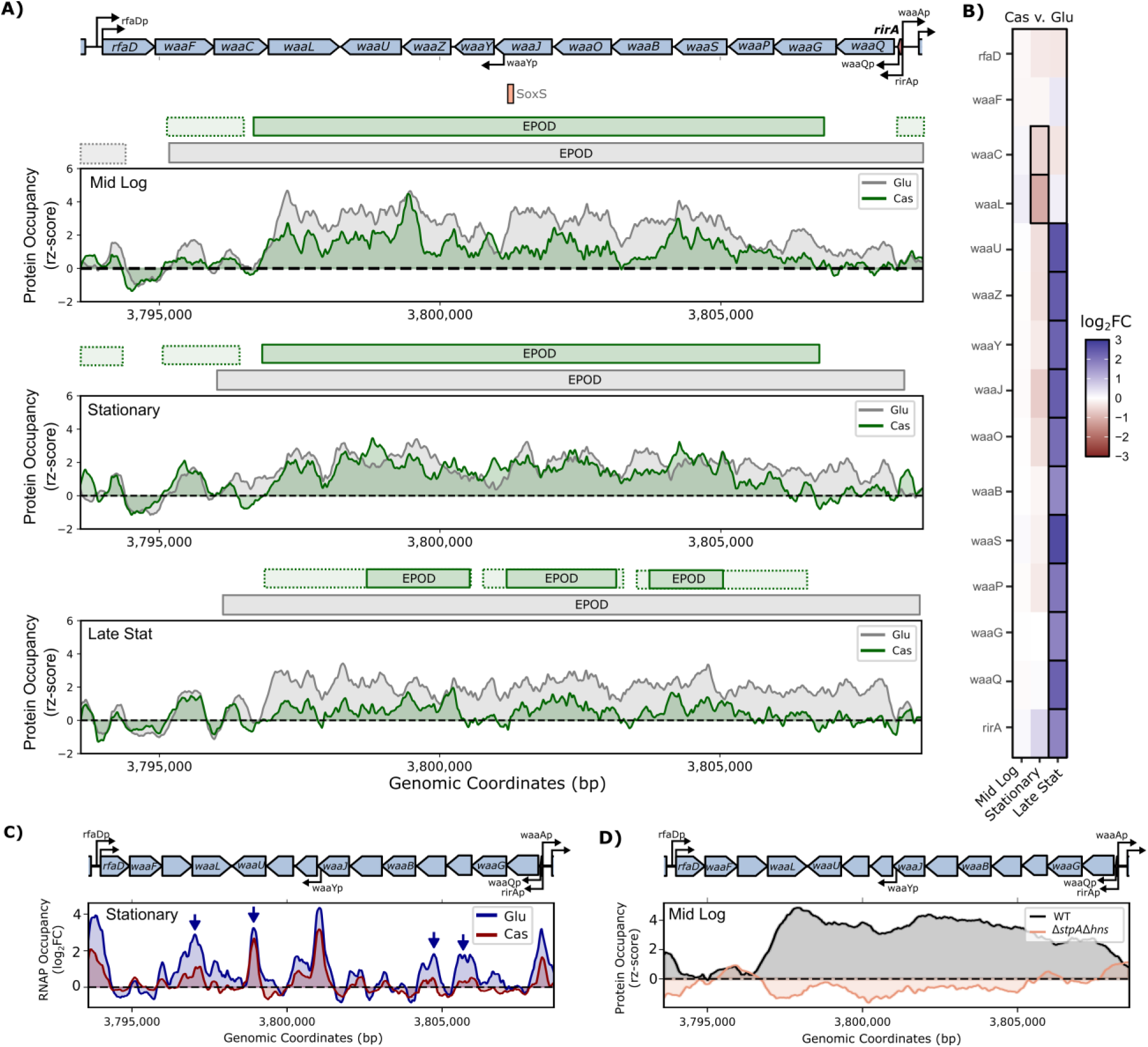
Static EPODs repress internal cryptic promoters regulating cell wall composition in both media, example at lipopolysaccharide biosynthesis operon. **A)** Protein occupancy (via IPOD-HR rz-scores, moving average window of 250 bp) plotted over genomic coordinates surrounding a static EPOD containing the lipopolysaccharide core biosynthesis operons: *rfaDwaaFCL*, *rirA*, *waaQGPSBOJ* and *waaYZU*. Gene locations and promoters are annotated with blue and black arrows respectively above plots, with sRNA encoded region (RirA) displayed in red in top right. Known regulator binding sites from RegulonDB are illustrated below the genes (orange: activator, purple: repressor). Three comparable growth stages (top: mid logarithmic, middle: stationary, bottom: late stationary) are plotted for both Glu (grey) and Cas media (green). Regions of EPODs are illustrated with horizontal bars above each plot, color according to media, strict EPODs with solid borders, and loose EPODs with dashed borders. **B)** Heatmap of the differential expression (log_2_FC) of each gene in Cas versus Glu media (*i.e.,* positive values indicate higher expression in Cas medium, while negative values indicate higher expression in Glu medium). Significant log_2_FC values (p-adj < 0.05) illustrated with black borders. **C)** An example of the RNAP occupancy (log_2_FC of RNAP ChIP signal to input, moving average window of 50 bp) at the discussed operons. Plotted values are at stationary phase in Glu (blue) and Cas (red) media. Blue vertical arrows indicate internal cryptic promoters within the operons. **D)** Protein occupancy (rz-scores) of wildtype (WT, grey) versus the genomic double deletion mutant (Δ*stpA*Δ*hns*, orange) at mid logarithmic growth stage surrounding discussed operons, taken from prior IPOD-HR study (11) (NCBI GEO GSE164796). Note the lack of protein occupancy in the deletion strain, supporting H-NS and StpA contributions to these EPODs.

Interestingly, from the RNAP ChIP results (example during stationary phase plotted in **Figure 2C**) internal cryptic promoters are still moderately bound by RNAP even when the EPOD is present in both media; the most prominent peaks are indicated with blue arrows. These internal promoters, while less strong than the promoters that occur outside of the EPODs (such as *rfaDp* and *waaQp*), highlight that while these EPODs may condense the DNA to help suppress transcription, RNAP is still able to access these promoters to a moderate level, in line with the RNAP stalling mechanism of EPODs (14, 15) – that is, it appears that much of the transcriptional repression in EPODs may not be from stopping RNAP binding per se, but rather, preventing promoter clearance and productive elongation. Indeed, from a prior IPOD-HR study comparing protein occupancy in wildtype and nucleoid-associated protein knockout strains (NCBI GEO GSE164796) (11), the EPODs noted here are likely made up of H-NS and StpA (both implicated in the RNAP stalling mechanism of transcriptional termination), as noted by the loss of protein occupancy and concomitant increased transcription in the Δ*stpA*Δ*hns* strain (**Figure 2D**; see also (11)).

In contrast with static EPODs, dynamic EPODs occur in at least one but not all growth stages in a given medium. An example of a dynamic EPOD region is illustrated in **Figure 3A** for the *hdeAByhiD*, *hdeD*, *arrS*, and *gadEF* operons. These genes are involved in the acid stress response (*gadEF, arrS*, *hdeABD*) and regulating cell wall composition (*yhiD*, *hdeABD*). As illustrated with the color-coded bars above the protein occupancy rz-scores, the entire region is constitutively heterochromatinized in Glu medium (grey); in contrast, an EPOD is present only in the divergent *arrS/gadF* promoter region during exponential and stationary growth in Cas, with the region overall more accessible. It is also notable that the EPOD over this region in Glu is fairly continuous in occupancy over the ∼5 kb locus, whereas especially at later timepoints the occupancy in this region in Cas media has defined protein occupancy peaks near documented promoters, likely indicating binding of local regulators rather than heterochromatinization. As indicated with green arrows, three prominent peaks in the Cas medium occur at the promoters of *hdeAp*, *hdeDp*, *gadEp*, and *arrSp*. These well-studied promoters are regulated by acid-responsive transcription factors (GadX, GadW, GadE, and YdeO) among others, illustrated with the known binding sites in orange (activators) and purple (repressors) bars. Importantly, these promoters control the transcription of two sRNAs, ArrS and GadF, as well as transcripts targeted by sRNAs to control mRNA processing and translation (*hdeD* by RprA and CyaR (62), and *gadE* by ArrS (63)).

**Figure 3.**
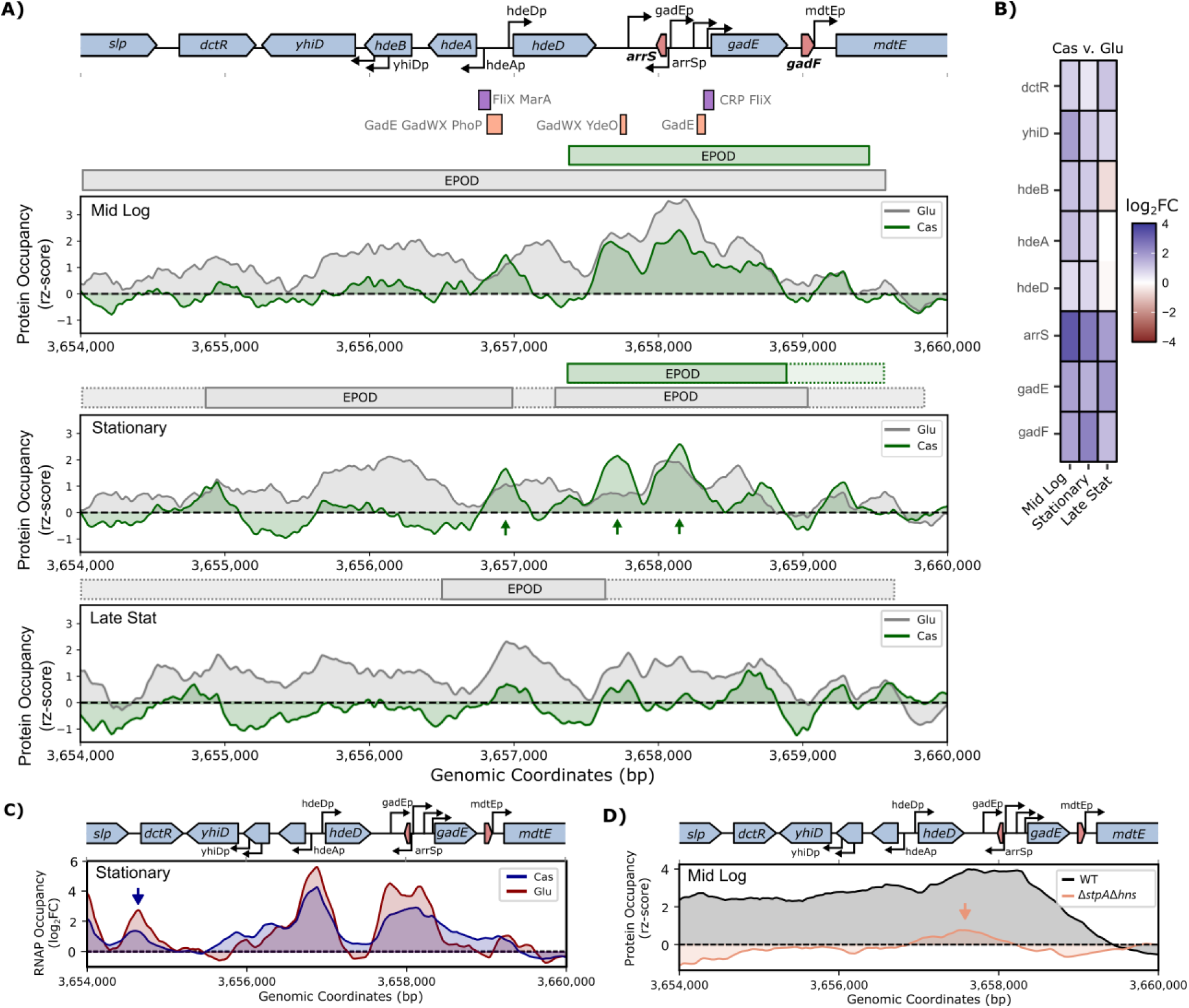
Dynamic EPODs coordinate nucleoid-associated protein with transcription factors to regulate gene expression, example at acid response operon. **A)** Protein occupancy (via IPOD-HR rz-scores, moving average window of 250 bp) plotted over genomic coordinates surrounding a dynamic EPOD region containing acid response and cell membrane genes: *hdeAByhiD*, *hdeD*, *arrS*, and *gadEF*. Gene locations and promoters are annotated with blue and black arrows respectively above plots, with sRNA encoded regions (ArrS and GadF) highlighted in red. Known regulator binding sites from RegulonDB are illustrated below the genes (orange: activator, purple: repressor). Three comparable growth stages (top: mid logarithmic, middle: stationary, bottom: late stationary) are plotted for both Glu (grey) and Cas (green) media. Regions of EPODs are illustrated with horizontal bars above each plot, color according to media condition, strict EPODs with solid borders, and loose EPODs with dashed borders Vertical green arrows point to three distinct protein occupancy peaks attributed to known transcription factor binding sites. **B)** Heatmap of the differential expression (log_2_FC) of each gene in Cas versus Glu media (i.e., positive values indicate higher expression in Cas media, while negative values indicate higher expression in Glu media). Significant log_2_FC values (p-adj < 0.05) illustrated with black borders. **C)** An example of the RNAP occupancy (log_2_FC of RNAP ChIP signal to input, moving average window of 50 bp) at the discussed operons. Plotted values are at stationary phase in Glu (blue) and Cas (red) media. Blue vertical arrow indicates a likely cryptic promoter for the transcriptional regulator gene *dctR*. **D)** Protein occupancy (rz-scores) of wildtype (WT, grey) versus the genomic double deletion mutant (Δ*stpA*Δ*hns*, orange) at mid logarithmic growth stage surrounding discussed operons, taken from prior IPOD-HR study (11) (NCBI GEO GSE164796). Note the lack of protein occupancy in the deletion strain, supporting H-NS and StpA contributions to these EPODs. Vertical orange arrow highlights the residual protein occupancy (although severely muted) that remains after deletion of H-NS and StpA, supporting contribution of other proteins like transcription factors to this region’s protein occupancy.

While different in protein binding behavior, it is notable that the entire genomic region shows similar expression patterns in which all genes are significantly more expressed in the Cas medium compared to Glu, with the slight exception of *hdeD* and *hdeAB* in late stationary phase (**Figure 3B**). Like as for the static EPOD discussed above, putative cryptic promoters are apparent in **Figure 3C**, such as the RNAP peak preceding *dctR* as indicated by the arrow. The prior IPOD-HR results using the Δ*stpA*Δ*hns* strain (NCBI GEO GSE164796) (11) again lacks most protein occupancy at the *hdeAByhiD*, *hdeD*, *arrS*, and *gadEF* operons (**Figure 3D**) compared to the wildtype strain indicating broad silencing of this region by H-NS and StpA. However, as indicated with the arrow, the double deletion fails to remove all protein occupancy at the promoters. It is likely that the remaining occupancy is binding of the known transcription factors regulating this region – thus, we observe a regulatory hand-off between constitutive silencing of the entire genomic region shown in **Figure 3A** in Glu media, to primarily local regulators activity in Cas media.

The differing protein binding behavior of the static and dynamic EPODs indicate two regulatory chromatin architectures occurring during nutrient stress. For the heterochromatin-like EPOD mode as previously proposed (11, 14, 15, 61), densely bound nucleoid-associated proteins suppress transcription of promoters within the EPODs through blocking or stalling of RNAP. As shown with the first example (**Figure 2**), static EPODs can follow this type of mechanism. Alternatively, as shown in the second example (**Figure 3**), a more complex mechanism with coordination between nucleoid-associated proteins and transcription factors may regulate transcription of other operons, as was observed for some of the dynamic EPODs. The integration of these regulatory modalities appears to be of critical importance, where for example we can observe heterochromatinization and silencing of a locus under one condition, giving way to control by local regulators in another – that is, loss of the EPOD leads to de-repression of a region, but this does not necessarily trigger transcriptional activation, but simply permits local control of the contained genes. We observe similar behavior at the curli synthesis operons *csgBAC* and *csgDEFG* (**Figure S6**) in which known transcription factor peaks appear mid-EPOD to activate transcription. Thus, transcription within static EPODs can also be coordinated between nucleoid-associated protein and transcription factors. It is also important to note that EPOD regions which appear static under conditions so far studied, may in fact still dissociate to allow de-repression of the genes that they contain under other physiological conditions that we have not considered.

### sRNAs disproportionately drive nutrient stress responses and are subject to complex transcriptional regulation

Upon global examination of the transcriptomic changes across the nutrient stress conditions studied here, we observed a consistent over-representation of sRNAs among transcripts showing condition-specific induction (**Figure 4A**); this effect was especially prominent during entry into stationary phase. The volcano plots provided in **Figure S1** of differentially expressed genes (p-adj value <0.05, |log_2_FC|>1) demonstrate the striking number of sRNAs regulated in both media during periods of transition. Namely, the entry into stationary phase resulted in differential expression of 40/91 sRNAs in the Cas medium and 42/91 in the Glu medium, with the majority (36 and 35 respectively) being induced. The transition from exponential growth to stationary phase requires a major overhaul in cellular metabolism, as nutrients become scarce cells must shift to less favorable metabolites and reduce replication rate. The large number of sRNAs induced during this growth transition implicates sRNAs as important regulators of adaptive cellular metabolism during the nutrient stresses of glucose and amino acid starvation.

**Figure 4.**
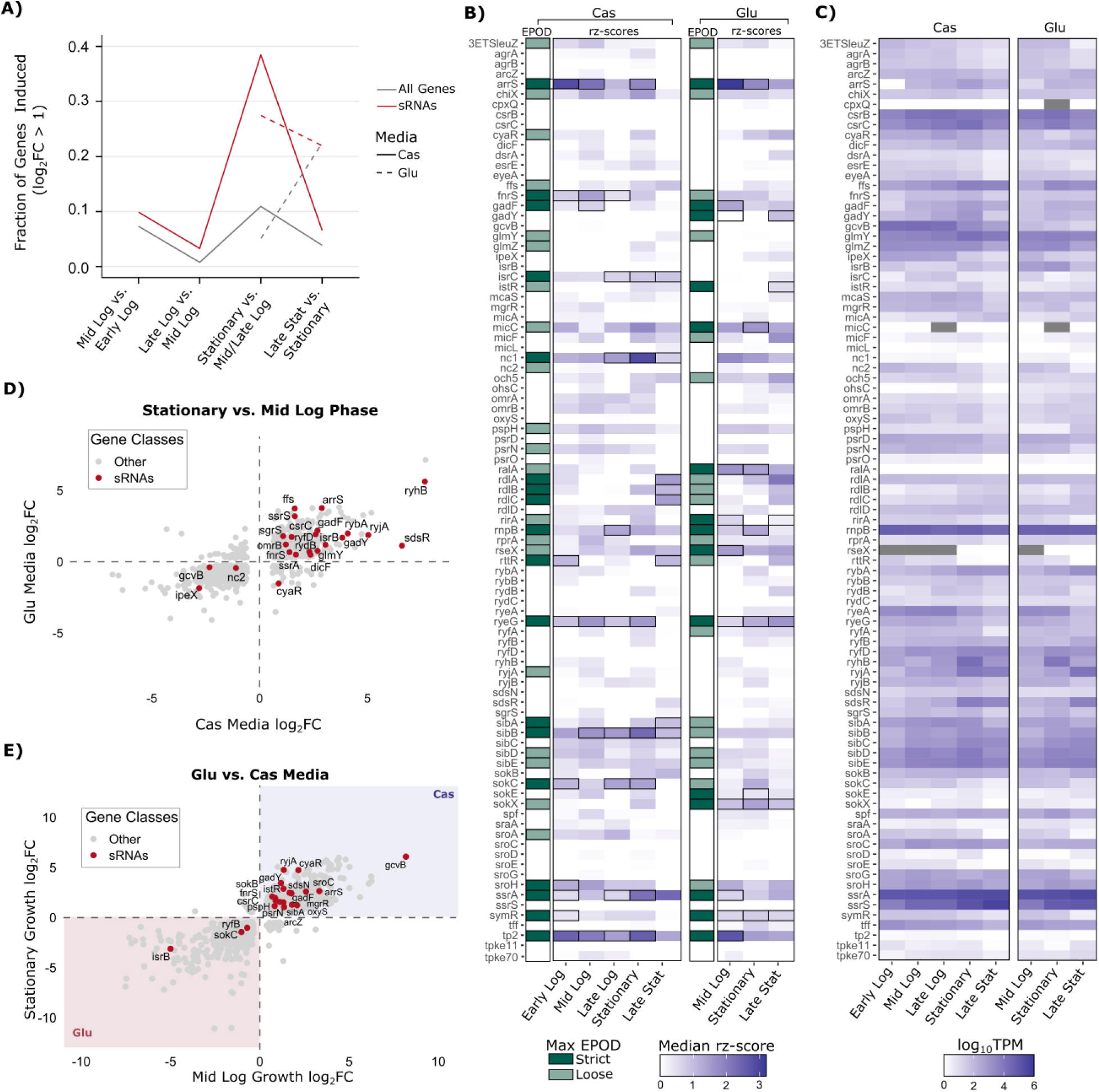
Dynamic protein occupancy and expression of sRNAs suggest diverse transcription factor regulators. **A)** Fraction of all genes (out of 4442, grey) and sRNAs (out of 91, red) significantly induced (p-adj < 0.05, log_2_FC >1) between each growth phase in each media (Cas: solid line, Glu: dashed line). **B)** Heatmap displaying the maximum EPOD classification (strict: dark green; loose: light green; never contained within an EPOD region: white) as well as the median rz-scores of the DNA regions [-200, +50] bp surrounding the encoded 91 well-annotated sRNAs investigated across the 8 growth and media conditions. Conditions in which the DNA region encoding the sRNA is contained within a strict EPOD are noted with black borders. **C)** Heatmap of the expression of the 91 sRNAs (log_10_TPM) in the 8 conditions tested. Grey indicates values in which the sRNA was not detected (TPM = 0). **D-E)** Significant differentially expressed genes (p-adj < 0.05) in **D)** both Cas and Glu media (log_2_FC values plotted on each axes) in stationary phase as compared to mid logarithmic phase, and **E)** both mid logarithmic and stationary phase in Cas vs. Glu media. Genes only differentially expressed in one of the plotted conditions are not included. sRNAs are plotted in red, all other genes in grey. Genes induced or repressed in both media (as depicted in panel D) suggest a growth-dependent regulation, while genes located in the shaded regions of panel E (blue: Cas, red: Glu media) suggest media-dependent regulation.

Upon cross-referencing with our protein occupancy data at locations surrounding well-annotated sRNAs (**Table S1**), a minority of sRNAs do occur in normally heterochromatinized regions. A heatmap is presented in **Figure 4B**, in which all sRNAs studied are listed with the median rz-scores (blue shading) of the [-200, +50] bp window surrounding the sRNA encoded region in all tested conditions. In addition, a categorical box (green) indicates the maximum EPOD level that fully contains the sRNA encoded region in each media (loose: light green; strict: dark green). Black borders surrounding the rz-scores indicate a strict EPOD call occurs over the sRNA encoded region in the given condition.

No sRNA-encoding DNA region is contained within a strict EPOD in all 8 tested conditions. However, the region encoding for sRNA RyeG is contained in a strict EPOD for 7/8 total conditions. RyeG encodes a toxic peptide, YodE, and therefore is expected to have repressed transcription during most conditions for optimal cell fitness. Other DNA regions encoding for sRNAs with toxic roles located within EPODs in at least one tested condition include many of the toxin-antitoxin sRNAs (SibA, SibB, RalA, IstR, RdlA, RdlB, RdlC, and SokE), the toxin encoded SokC, and the prophage-related IsrC. However, as mentioned, even with toxic roles, regions encoding these sRNAs are predominantly not located within EPODs in the tested conditions. Only 33% of the 91 sRNAs studied were contained within an EPOD in at least one condition, totaling only 65 of the possible 728 instances.

To compare EPOD occurrence to sRNA transcriptional regulation, the RNA-seq expression levels for all sRNAs (denoted as log_10_TPM) are provided in a heatmap in **Figure 4C** (differential expression results provided in **Figure S7**). Based on these data, and cross-referencing our protein occupancy, we observe several distinct patterns of regulation for sRNAs. Some sRNAs appear to follow the heterochromatin-like mode of silencing, such as Tp2 and SymR; both sRNAs are encoded within loose or strict EPODs for all conditions (and thus would be considered within static EPODs in our nomenclature above), resulting in low TPM counts and no differential expression across conditions. Alternatively, RyeG, located within an EPOD in 7/8 conditions, while lacking differential expression in most conditions, has moderate TPM counts suggesting more complex transcriptional regulation. As such, some of the more dynamically expressed sRNAs rarely occur within an EPOD, such as RyhB, IsrB, and GcvB. The lack of EPOD transcriptional silencing for most sRNAs suggest transcription factor regulation may predominantly control sRNA expression during starvation conditions; this regulatory paradigm appears appropriate for frequently-expressed sRNAs which may require rapid induction in the presence of commonly-encountered nutrient stresses.

### Nutrient and growth-dependent sRNA expression likely relies on complex transcription factor regulation

At the level of transcript abundance, a total of 88 of the annotated 91 sRNAs listed in **Figure 4C** were detected in all conditions, with RseX, not detected during early growth conditions and MicC and CpxQ lacking detection during late logarithmic or stationary growth (**Table S2**). Furthermore, 9 of the studied sRNAs that were detected in all conditions showed no statistical differences in expression between conditions tested (p-adj > 0.05 for all comparisons): ChiX, DsrA, Nc1, RalA, SokX, SraA, PsrO, SroE, Tp2).

To compare across growth, the log_2_FC values of genes differentially expressed in stationary versus mid logarithmic phase in both media are plotted in **Figure 4D** (genes differentially expressed in only one of the two conditions are not plotted). Notably, 18 sRNAs are induced and 3 (GcvB, Nc2, and IpeX) are repressed during starvation in both media conditions. The genes induced or repressed consistently between media conditions suggest growth-dependent regulation, such as by the stationary phase induced general stress response regulator RpoS. However, only half of these 22 sRNAs are known to be regulated by RpoS (ArrS, CyaR, GadF, GadY, RyjA, SdsR, SgrS, SsrS) or encoded in regions overlapping an EPOD in at least one of the compared conditions (ArrS, FnrS, GadF, GadY, GlmY, SsrA). The lack of RpoS and apparent EPOD regulation for the remaining 11 sRNAs suggest growth-dependent regulation at the level of transcription factors. A similar plot for the late stationary phase comparisons between media is provided in **Figure S8**.

To compare across media, the log_2_FC values of genes differentially expressed between Cas versus Glu media at mid logarithmic and stationary growth phases are plotted in **Figure 4E**. As color annotated, positive log_2_FC values mean higher expression in the Cas medium, while negative log_2_FC values mean higher expression in the Glu medium. Therefore, genes with positive or negative log_2_FC values in both mid logarithmic and stationary phase suggest media-dependent regulation. Notably, only 3 sRNAs (IsrB, SokC, and RyfB) are expressed more in Glu medium during mid logarithmic and stationary phase compared to Cas medium, while 18 are more expressed in Cas medium. Seven out of 21 of these sRNAs (ArrS, FnrS, GadF, GadY, IstR, SibA, SokC) are encoded in regions overlapping an EPOD region during at least one of the compared conditions (and thus might be controlled at the level of chromatin structure), pointing again to classical local regulation in the majority of cases. Similar plots for stationary versus late stationary phase, and mid logarithmic versus late stationary phase, are provided in **Figure S8**.

The diversity of sRNA expression during glucose (Cas) or amino acid (Glu) starvation, and the variety of clearly apparent regulatory strategies controlling them (with RpoS, EPODs, and local regulators all implicated in some cases), further support the complexity of nucleoid-associated proteins and transcription factors coordinating regulation of sRNAs, and raises the important question of precisely which signals (and which regulators) are responsible for sRNA activation under different nutrient stress conditions.

### Competitive binding by FNR, NarP, and PhoP repress IsrB for anaerobic metabolism

An emblematic example of the complex regulatory logic driving small RNA expression is observed at the promoter of the dual-function carbon regulating sRNA IsrB (**Figure 5A**). IsrB (also known as AzuR) (64–66), while already known to be regulated by the secondary carbon source regulator, CRP, is likely subject to additional transcriptional regulation. IsrB represses translation of two mRNAs, *cadA* and *galE* (64) that encode for enzymes involved in weak organic acid (67–69) and galactose metabolism (70), respectively, while also encoding the small peptide, AzuC, that supports aerobic glycolysis (66). Given IsrB’s aerobic metabolism function, an acidic and an anaerobic regulator that turns off IsrB expression to enable translation of *cadA* and *galE* have yet to be identified (66). In our RNA-seq data, we observed that IsrB shows a ‘pulsed’ expression pattern in both Glu and Cas media, with expression peaking in stationary phase and then dropping again in late stationary phase. The *isrB* promoter region contains two previously annotated CRP binding sites (64, 66), but the primary changes in protein occupancy visible using the IPOD-HR method occur not at those sites, but further upstream, prompting us to search for additional transcriptional regulators acting on this region. Indeed, we identified motif hits for FNR, PhoP, and NarP (**Figure 5A**), all occurring within 100 bp of the putative *isrB* transcription start site. A survey of protein occupancy across our experimental conditions indeed shows that substantial protein occupancy shifts are observed across conditions at those binding sites, with particularly high occupancy of the putative FNR and PhoP sites in Cas media during stationary phase, and the putative NarP-bound region in late stationary phase.

**Figure 5.**
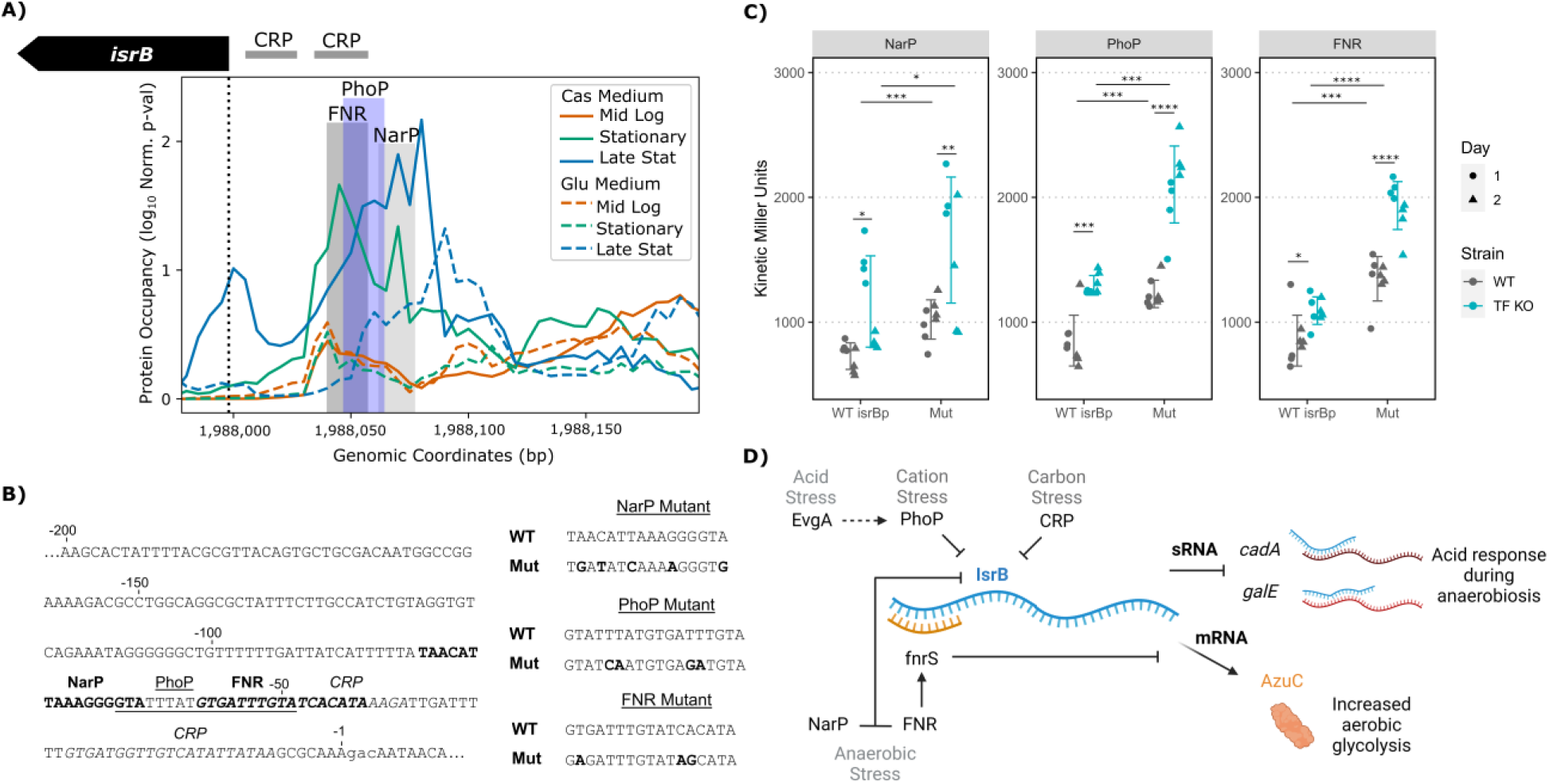
Overlapping transcriptional repression by FNR, NarP, and PhoP restricts IsrB function to aerobic conditions. **A)** IPOD-HR protein occupancy (log_10_ normalized p-values of rz-scores) at the DNA promoter of IsrB (also known as AzuCR) shows varied protein binding locations and intensities. As nutrients become scarce during later growth stages, like stationary and late stationary phases, protein occupancy increases particularly ∼30-90 bp before the transcription start site (annotated with the vertical dashed line). Some transcription factor motifs of particular interest found in these regions are shaded. The previously predicted binding sites of CRP are shown by horizontal bars above the plot. **B)** The promoter sequence [-200, 10] bp surrounding the transcription start site (lowercase) used in the Miller assays is listed. Transcription factor motif sequences are bolded (NarP and FNR as labeled), underlined (PhoP), and italicized (CRP). The rationally designed mutants (Mut) for each motif are listed compared to wildtype (WT). **C)** The genomic transcriptional reporters were tested via Miller assay for four biological replicates on two separate days (indicated by shape) for each of the tested transcription factors. Results shown are in Glu media at stationary phase to mimic conditions of significant plasmid screening results. Significance by unpaired two-tail t-test p-values as indicated: * <0.01, ** <0.005, *** <0.001, **** <0.0005. Mutant results mimicked the behavior of the transcription factor knockout strain (TF KO) with wildtype promoter (WT isrBp) (*i.e.*, increase in KMUs upon alleviation of repression when transcription factor is knocked out compared to when the binding site is mutated). **D)** Updated regulatory networks of IsrB with the additional transcriptional repressors noted along with prior study findings. Created in BioRender.com.

To directly assess the potential effects of FNR, PhoP, and NarP on *isrB* transcription, we tested rationally mutated IsrB promoters for each transcription factor via genomic Miller assays (**Figure 5B**). As shown in **Figure 5C** in grey, the kinetic Miller Units (KMUs) measured at stationary phase in Glu medium show all mutant promoters resulted in partial or full alleviation of repression in the wildtype strain (*i.e.*, increased KMUs compared to the wildtype promoter, WT IsrBp). The level of alleviation can also be compared to the wildtype promoter in the respective transcription factor knockout (KO) strains shown in blue. These three regulators explain the missing link for IsrB expression: how CadA and GalE translation are enabled during anaerobic (FNR and NarP) and low pH (PhoP) environments (**Figure 5D**).

### Chromatin structure-informed bioinformatic approach proposes a suite of transcription factors regulating sRNAs

The IsrB locus provides a concrete example of what we expect is a general trend, visible in our datasets, in which complex interactions of local regulators (and potentially also nucleoid-associated proteins) may regulate sRNAs, often through activity at binding sites that have not been previously characterized. We previously developed a bioinformatic approach to propose potential regulators of sRNAs using a limited IPOD-HR dataset (51). This proof-of-concept was experimentally validated for the elusive sRNA, RseX, which had no known transcriptional regulator until the nucleoid-associated protein, H-NS, was confirmed to silence its expression (51). In alignment, we find in our present dataset that RseX is indeed contained within loose or strict EPODs during 7/8 tested conditions (**Figure 4A**), likely comprised of H-NS. Given the prior success, we aimed to expand this bioinformatic approach using the nutrient stress IPOD-HR datasets with corresponding RNA-seq datasets, and experimentally following up on the predicted transcription factors of sRNAs.

We first identified protein binding regions at each growth phase for the two media conditions using peak calling on the IPOD-HR -log_10_ p-adjusted rz-scores (labeled as log_10_ pval). Regions surrounding the 91 identified sRNAs, overlapping or within the region 200 bp upstream of the transcription start site through 50 bp past the sRNA coding region, were compiled and merged across growth phases in each media. An example of merged protein peak regions for the sRNA, ArrS, in Cas medium is provided in **Figure 6A**. A total of 341 peaks were identified across all conditions and sRNAs studied (**Table S4**).

**Figure 6.**
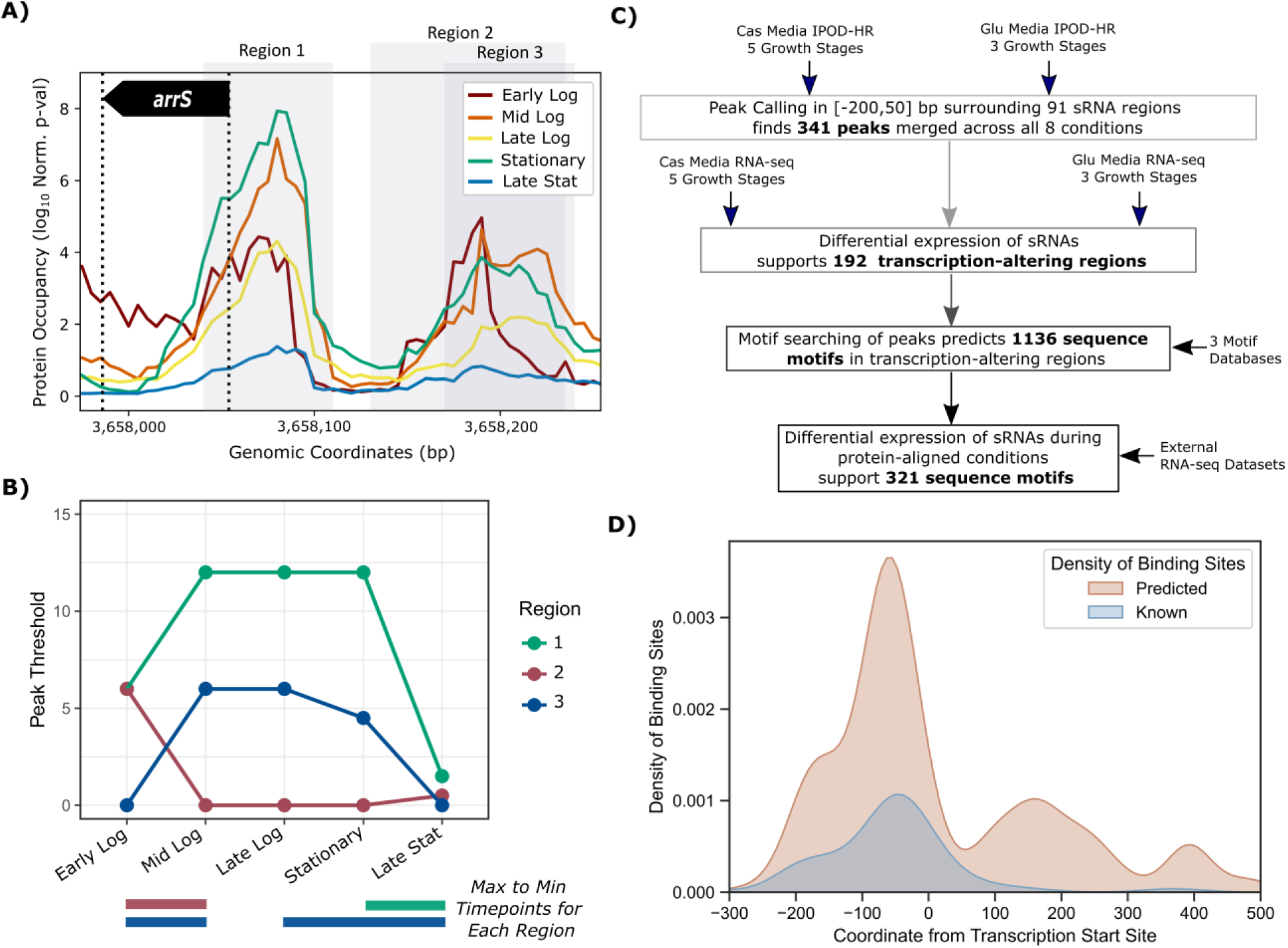
Bioinformatic approach informed by IPOD-HR and RNA-seq predicts transcriptional regulators of sRNAs, ArrS as an example. **A)** IPOD-HR protein occupancies (Log10 normalized p-value of rz-scores) at the 5 tested growth stages in Cas media surrounding the DNA region encoding the sRNA, ArrS (annotated with vertical dashed lines and black arrow). Protein occupancy regions identified via peak calling are shown by the grey bars, merged between all conditions. **B)** The protein occupancy maximum signal threshold for each called peak region is plotted over growth for Cas media. The range of growth stages from maximum to minimum protein occupancies for each region are highlighted with color-coordinated bars below each plot. These ranges are compared to the RNA-seq differential expression results (listed in Table S2). **C)** A workflow illustration emphasizing the internal datasets (vertical arrows) and external datasets (horizontal arrows) that were used in this study to filter down predicted transcriptional regulators of sRNAs. **D)** Density plot of the external RNA-seq supported binding motif locations (a total of 321), annotated as “Predicted”, compared to known binding motifs locations for sRNAs curated by RegulonDB. The coordinate axis is annotated from the transcription start site at 0, negative values indicate upstream.

The maximum peak threshold value for each region and time point was tracked, as illustrated in **Figure 6B**. The growth phase(s) of maximum protein occupancy to minimum protein occupancy in each media was identified. These growth timeframes of maximum to minimum protein occupancy for each region were compared to the RNA-seq differentially expressed conditions. If significant differential expression of the respective sRNA occurred over this growth timeframe, that region is identified as a likely transcription-altering region. For example, as shown in **Figure 6B**, maximum protein binding for Region 2 for the sRNA, ArrS, in Cas medium occurs in early logarithmic growth while minimum first occurs in mid logarithmic growth. ArrS was also found to be differentially expressed in mid logarithmic compared to early logarithmic growth. Therefore, Region 2 was identified as a likely region where protein(s) bind to regulate ArrS transcription. Of the 341 peak regions found surrounding the sRNAs, 192 were supported by the RNA-seq differential expression. In addition, 51 of the identified peak regions overlap with previously reported protein binding sites listed by RegulonDB (**Table S4**).

The 341 identified peak regions were also put through a sequence motif searching algorithm, FIMO (52), using three previously established motif databases: Swiss Regulon (54), dpInteract (53), and Prodoric2 (55). All motifs with E-values < 0.0001 were compiled (**Table S4**). A total of 1911 sequence motifs were identified to overlap or be contained within the [-200, +50] bp region surrounding the 91 sRNAs. These motifs were filtered to only those found in the 192 identified transcription-altering regions, resulting in 1136 sequence motifs (63 of which agree with known protein binding sites curated by RegulonDB (60)).

The 1136 motifs located in transcription-altering peaks were further supported using external RNA-seq differential expression previously performed (51). For example, Region 2 of ArrS (**Figure 6A**) had a predicted GadE sequence motif which was supported by an external Δ*gadE* RNA-seq dataset (71) which showed significantly reduced ArrS expression. GadE is a known activator of ArrS transcription, thus serving as a positive control captured by this approach. A total of 321 sequence motifs located in transcription-altering peaks were supported by external RNA-seq datasets, covering 47 unique sRNAs (**Table S4**). A summary of the bioinformatic workflow is provided in **Figure 6C**, noting the combination of new data presented in this study (vertical arrows) with the external mined datasets (horizontal arrows).

A density plot depicting the location of the predicted 321 sequence motifs compared to the location of known DNA protein binding sites of sRNAs is shown in **Figure 6D**. Observed large concentration of sequence motifs upstream of the transcription start site were in agreement with literature, with most sequence motifs near the canonical promoter region (-35 to -10 bp before the transcription start site).

### Miller assay screening supports dozens of new sRNA-transcription factor connections

In order to benchmark the set of new binding sites identified in our pipeline, we thus biochemically tested the putative transcription factor motifs nearest the promoter region, [-100, 10] bp surrounding the transcription start site, excluding nucleoid-associated proteins and sigma factors. The resulting test set consisted of motifs for 24 transcription factors upstream of 30 sRNAs, for a total of 62 unique putative sRNA - transcription factor pairs. A plasmid-based screening method was designed using transcriptional Miller assays to biochemically test regulation by the candidate 24 transcription factors predicted to bind upstream of the 30 sRNAs. Simply, as depicted in **Figure 7A**, the sRNA promoter of interest, [-200, 10] bp surrounding the sRNA transcription start site, was cloned upstream of a synthetic ribosomal binding site (RBS) followed by the *lacZ* gene reporter. Note a bidirectional terminator was placed upstream of the sRNA promoter to prevent leakiness from the upstream pBAD promoter. These plasmids were transformed into both a wildtype and the corresponding transcription factor knockout (TF KO) strain. Transcription rates were measured at both mid logarithmic and stationary phases in either Cas or Glu media. To control for strain differences in RNAP availability (72), and potential differences in plasmid copy number due to strain health and antibiotic use (73), a control plasmid using a moderate Anderson Promoter (J23110) was used as a baseline control. The resulting quantification of transcription rate, termed kinetic Miller Units (KMUs) are normalized between wildtype (WT) and knockout (KO) and subtracted from the corresponding sRNA ratios, defined as the Relative KMUs (**Figure 7A**, right). Significant results from the plasmid-based screening are plotted in **Figures 7B-C** for Cas medium, and **Figures 7D-E** for Glu medium. All sRNA-transcription factor pairs were also tested in a rich-defined media (RDM) that contains both glucose and amino acids, removing the nutrient stress factor. Significant results in RDM are plotted in **Figures 7F-G**. All significant plasmid-based screening results are summarized in **Table 1**, with all data provided in **Table S5.**

**Figure 7.**
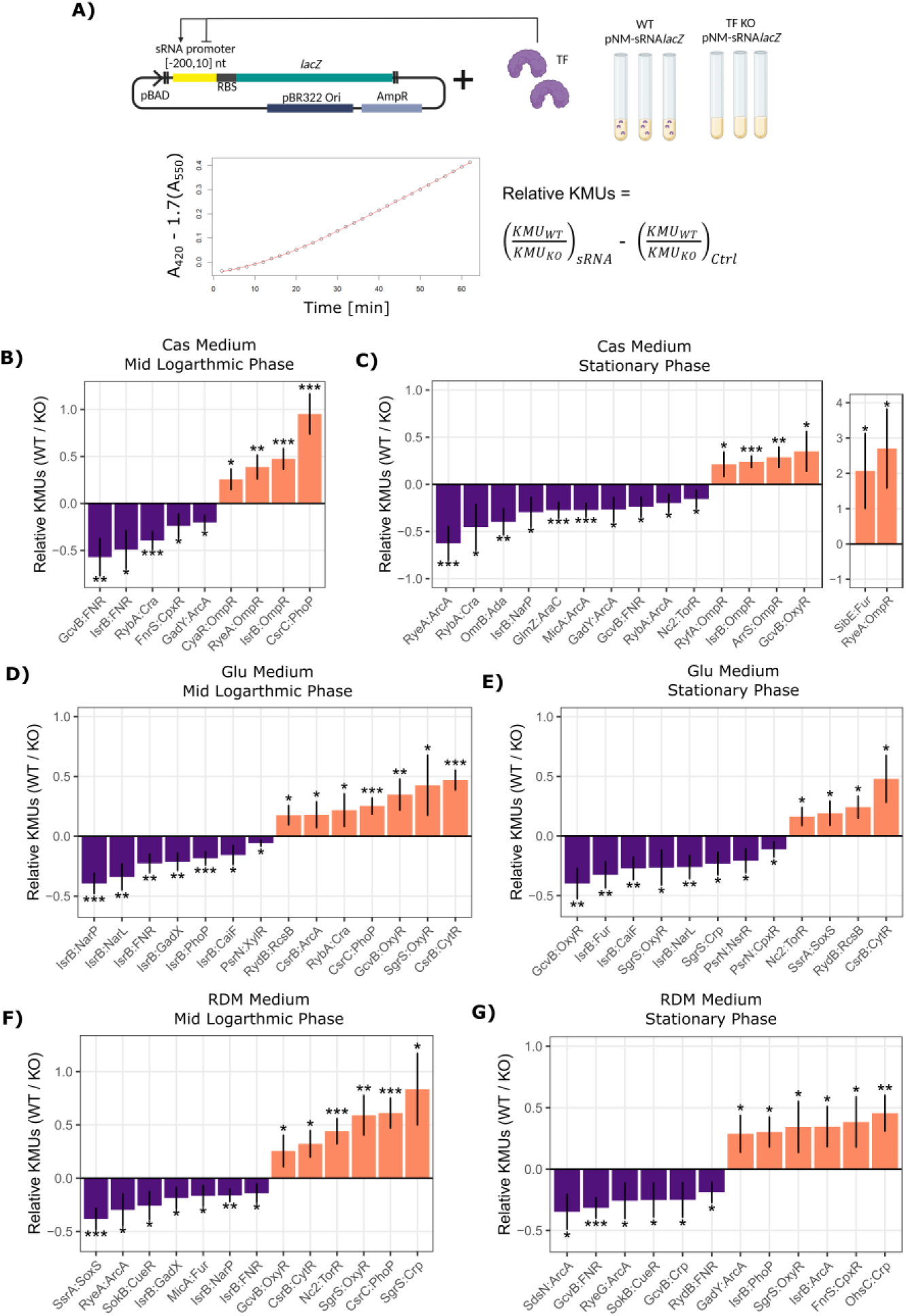
Plasmid Miller assays support the bioinformatically predicted transcription factors of sRNAs. **A)** Experimental set-up for the plasmid-based Miller assays to test the bioinformatically predicted transcription factors for sRNAs (predicted to bind within [-100, 0] bp upstream of the sRNA transcription start site) in both Cas and Glu media at mid logarithmic and stationary phases. Quantification of the Miller assay results are represented as normalized ratios for the sRNA-transcription factor (TF) pairs, where the kinetic Miller Units (KMUs) of the plasmid transcriptional reporters in wildtype (WT) is compared to the respective transcription factor knockout (KO), baselined to the Anderson promoter control (J23110) transcriptional reporter ratio (*i.e*., activators are > 0 while repressors are < 0). These normalized ratios are referred to as relative KMUs. Created in BioRender.com. **B-G)** Significant results of the plasmid Miller assays, represented as relative KMUs, at mid logarithmic and stationary growth phases in Cas **(B-C)**, Glu **(D-E)** and RDM **(F-G)** media, respectively. Transcriptional activators are shown in orange, repressors in purple. Significance is calculated by unpaired sample two-tailed t-tests each in biological triplicates (p-value: * < 0.05, ** <0.01, *** <0.005). Results of all tested sRNA-transcription factor pairs provided in Table S5.

**Table 1.**
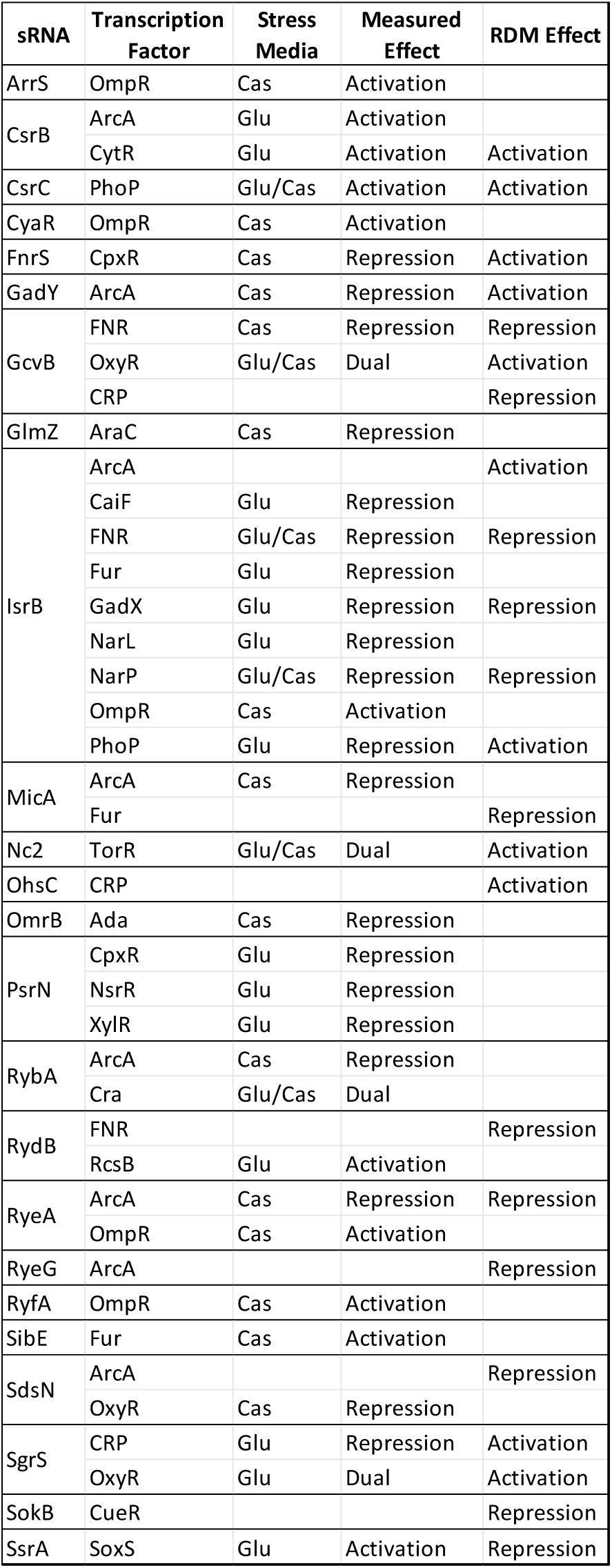
Novel transcription factors of sRNAs supported by plasmid Miller assays. List of transcription factor candidates for sRNAs that were predicted by the bioinformatic approach and supported by the plasmid-based Miller assays. Significant effects in rich-defined medium (RDM) are also provided (blank boxes in the “Stress Media” column indicate no significant effect was measured in the Glu or Cas media but measured in RDM). Results by plasmid support direct or indirect regulation of the sRNA expression by the candidate transcription factor.

Notably, 43 out of the predicted 62 sRNA-transcription factor pairs were found to significantly alter transcription in at least one of the media and growth conditions. The 43 significant sRNA-transcription factor pairs from the plasmid-based screening results suggest at least an indirect regulatory relationship, broadening potential sRNA function in the identified transcription factor’s associated stress networks. For example, RydB is a sRNA expressed in minimal media without any known targets or transcription factors (74). Given the transcriptional activation found for the “capsule synthesis” transcription factor, RcsB, in Glu media (**Table S5**), the sRNA RydB could regulate cell membrane composition.

To give a concrete example of the information provided in our pipeline, we considered the aforementioned case of *isrB* (**Figure 5**), where FIMO results show potential binding sites for several factors including FNR, PhoP, and NarP, as well several others (listed in **Table S4**). Based on external RNA-seq datasets that we previously analyzed (51), IsrB was repressed in low and high pH (75), anaerobic environments (76), differing carbon sources like fructose and acetate (77), and under iron starvation (78), among other conditions. Many of these factors are associated with anaerobic growth (ArcA, CaiF, FNR, NarL, and NarP), while others are associated with acidic environments, (GadX and PhoP). All were predicted to repress transcription in Glu medium, while FNR, NarL, NarP, OmpR, and PhoP were also predicted to activate transcription in Cas medium (**Table S4**). These predictions were screened via their respective transcriptional reporter plasmid in the wildtype and transcription factor knockout strains, and results are shown in **Figure 7B-G**, with all significantly altering the transcription of the IsrB promoter in the predicted direction. Thus, we observe that our integration of protein occupancy and transcriptional data enables the identification of previously cryptic regulatory connections controlling sRNA expression.

### Protein occupancy-informed predictions successfully identify direct transcriptional regulators of sRNAs during nutrient stress

To assess the impacts of new regulatory connections predicted by our pipeline in a more realistic biological context, we focused on a limited set of 21 sRNA-transcription factor candidates that had significant regulatory effects in the plasmid-based screening (**Table S5**). For this set, we sought to confirm direct functional regulation by the target transcription factors via mutating DNA binding sites in the sRNA promoters within the context of the *E. coli* chromosome. We tested sRNA promoter mutants that were rationally designed to reduce the ability of the transcription factor to bind the DNA sites. The mutant and wildtype promoters with downstream transcriptional reporter (**Figure 8A**) were genomically inserted at the *ara* operon. Genomic mutants were then tested alongside the unmutated promoters in wildtype strains for all pairs. The ratio of KMUs for the wildtype (WT) and mutant (Mut) promoters are plotted in **Figures 8B-C** for Cas medium, **Figures 8D-E** for Glu medium, and **Figures 8F-G** for RDM medium, at mid logarithmic and stationary phases, respectively. Twelve candidates exhibited reduced activity in the case of the mutant relative to wildtype promoter, in agreement with the plasmid screening results, supporting functional regulation due to direct protein binding (summarized in **Table 2**).

**Figure 8.**
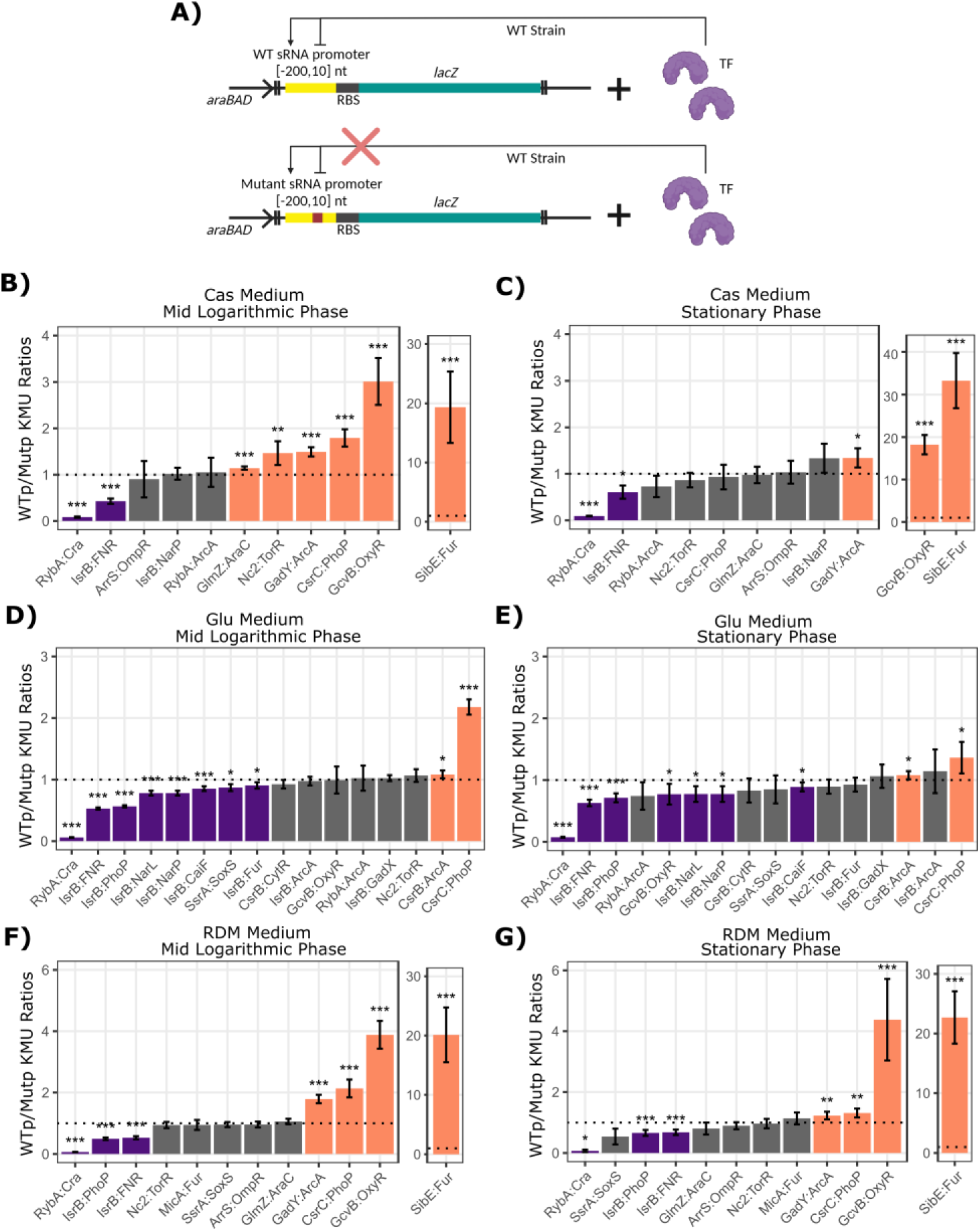
Genomic Miller assays confirm direct transcription factor regulation of sRNAs. **A)** 21 select sRNA-transcription factor candidates supported by the plasmid screening were further tested for direct functional regulation using genomic Miller assays. Rationally designed mutants (Mutp) of the transcription factor (TF) motif binding sites within the sRNA promoter were compared to the wildtype promoter (WTp) in the wildtype (WT) strain. Mutant promoters were designed to reduce the ability for the respective transcription factor to recognize and bind the DNA motif sequence. Created in BioRender.com. **B-G)** The ratios of wildtype to mutant promoter (WTp/Mutp) Kinetic Miller Units (KMUs) in the WT strain at mid logarithmic and stationary phases for Cas **(B-C)**, Glu **(D-E)** and RDM **(F-G)** media, respectively. Statistically significant ratios are colored in orange (activation, WTp/Mutp >1) and purple (repression, WTp/Mutp <1) and annotated with significance (from equal designated with dashed line, unpaired two-tailed t-test, each in biological quadruplicates, p-value: * < 0.05, ** <0.01, *** <0.005).

**Table 2.**
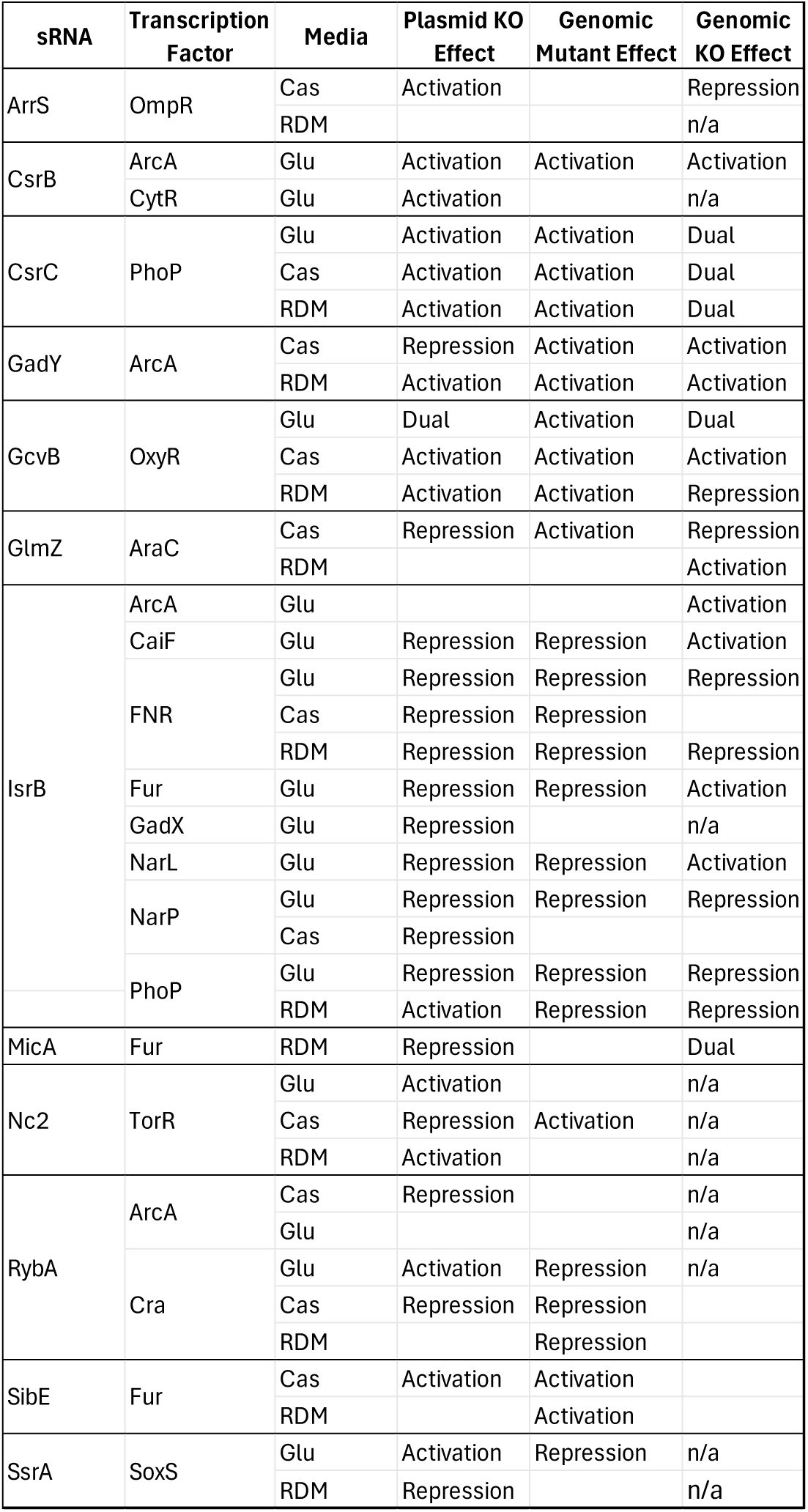
Summarized genomic Miller assay results for sRNA-transcription factor candidates. List of the 21 sRNA-transcription factor candidates tested by genomic Miller assays to support direct functional regulation by the transcription factors on sRNA expression. The measured transcriptional reporter plasmid effect in the wildtype to transcription factor knockout (KO) strains from Table 1 are provided for reference. Genomic mutant effect refers to significant effects between the wildtype and rationally designed mutant promoter in the wildtype strain. Genomic KO effect refers to significant effects genomically of the wildtype promoter measured between wildtype and the transcription factor KO strain. Boxes labeled n/a indicate not tested, while blank boxes indicate no significant effect measured.

The sRNA-transcription factor pairs with effective mutants were genomically tested in the transcription factor knockout (KO) strains using the wildtype and mutant promoters (**Figure S9**) to further verify the plasmid-based screening results of the identified transcription factor to the sRNA promoters. All data is provided in **Table S5** with summarized results in **Table 2**. Successful mutants that recapitulated the deletion strain results are defined as confirmed direct targets of regulation and summarized in **Table 3**. The novel regulation of these 5 sRNAs are highlighted with accompanying data in the following sections.

**Table 3.**
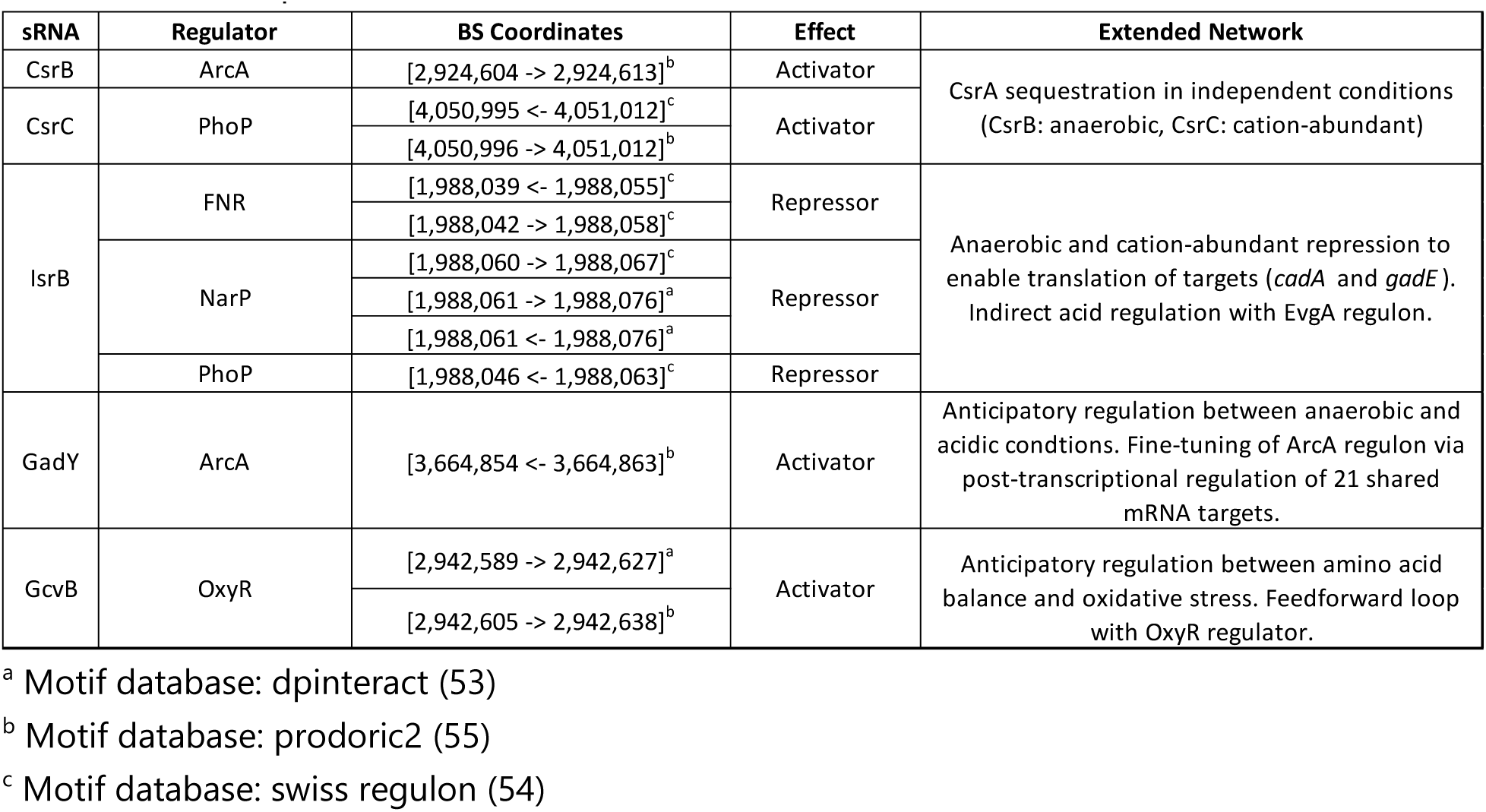
Confirmed transcription factor binding sites of sRNA promoters. Summary list of the sRNA-transcription factor pairs confirmed by the genomic Miller assays, with binding site (BS) coordinates provided of the found sequence motif that was rationally mutated for genomic transcriptional reporter testing. Coordinates based on reference genome U00096.3 with DNA strand indicated by arrow direction (-> positive, <-negative). A summary of the implications of these new transcription factors for the sRNAs is summarized.

### Divergent transcriptional regulation of CsrB and CsrC by ArcA and PhoP support independent roles for sequestering CsrA during glucose and amino acid starvation

A clear example of new *cis-*regulatory logic identified by our pipeline involves two critical post-transcriptional regulators of carbon metabolism: the sponge sRNAs CsrB and CsrC. CsrB and CsrC

(23, 79) are both sponge sRNAs that titrate availability of the carbon storage regulator protein CsrA. CsrA toggles translation of hundreds of mRNAs to tailor carbon metabolism (80, 81) and other stress responses (82); it has also been recently connected with sRNAs as a modulator of their post-transcriptional activities (83). CsrB and CsrC expression are controlled by a shared two-component system BarA/UvrY, that induces expression during an abundance of short-chain carboxylic acids (80, 81). Given their shared function, the sRNAs, CsrB and CsrC, are considered redundant without unique regulatory roles; however, their potential to have complementary, not fully overlapping, function remains. In our data, CsrB and CsrC had differing RNA-seq differential expression patterns without their promoters being contained within EPODs in any of the tested conditions (**Figure 4B-C**, **Figure S7**). Indeed, CsrC showed growth-dependent (**Figure 4D**) and media-dependent (**Figure 4E**) transcriptional regulation, while CsrB appears in neither comparison. These differences in expression patterns suggest divergent transcriptional regulation.

Upon inspecting IPOD-HR profiles for both CsrB (**Figure 9A**) and CsrC (**Figure 9B**) striking similarities emerge. Both show increasing protein occupancy ∼30 bp from the transcription start site throughout growth in Cas medium (solid lines). In Glu medium (dashed lines), peaks in mid logarithmic (orange) and stationary (green) phases are present at equivalent magnitudes, then quickly dissipate in late stationary phase. Note previously identified transcription factor binding sites for UvrY as well as the nucleoid-associated protein, IHF for CsrB (81), and carbon regulator, CRP for CsrC (84), are annotated with grey bars above the plots. While the protein occupancies are similar, the motif search resulted in different predicted transcription factors. CsrB had three potential activators in Glu medium, ArcA, CytR, and RcsB, while CsrC had two dual regulators in both media, Cra and PhoP.

**Figure 9.**
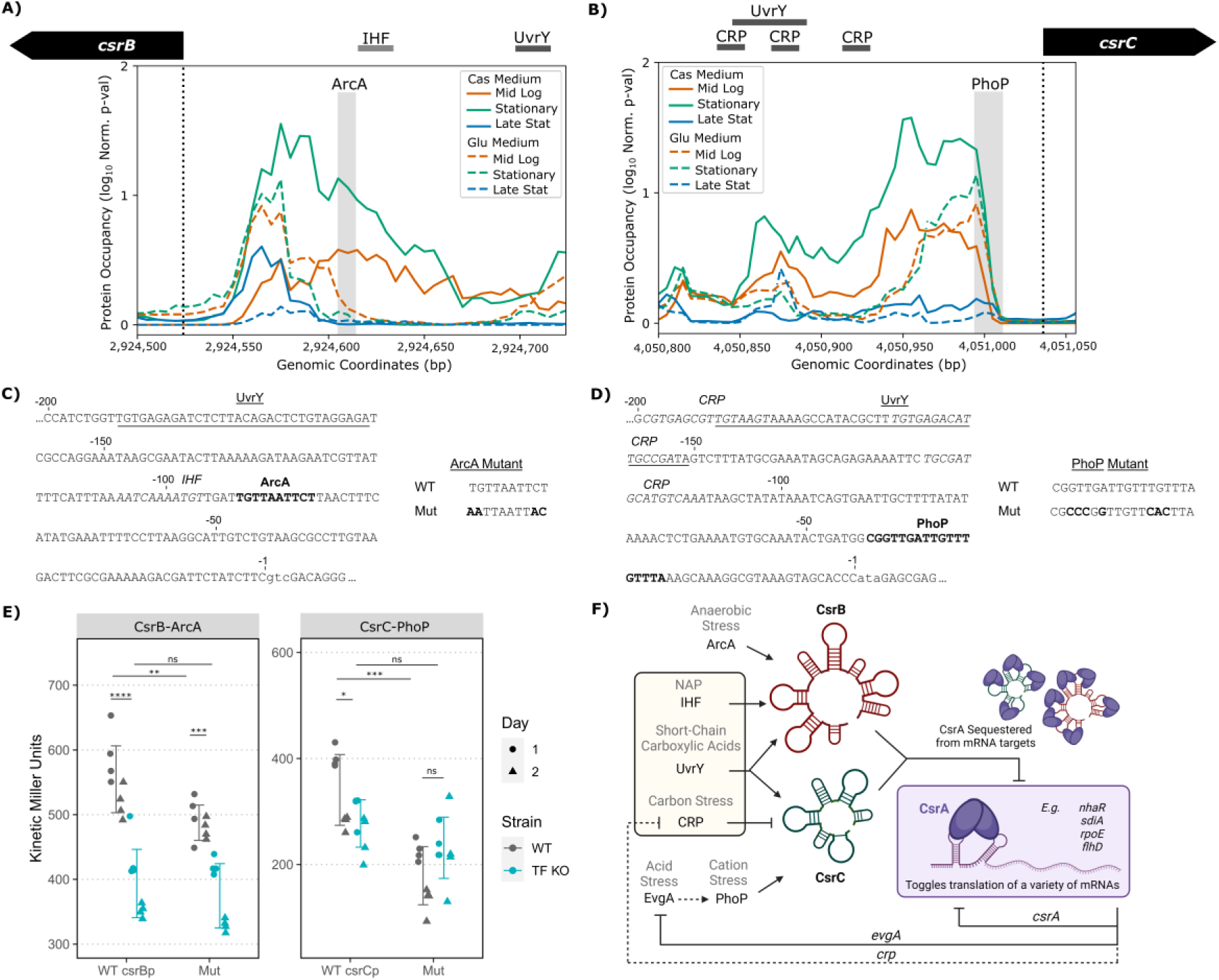
CsrB and CsrC independently regulated by ArcA and PhoP for fine-tuned sequestration of carbon regulator CsrA. IPOD-HR protein occupancy (log_10_ p-values of rz-scores) at promoters of sRNAs CsrB (**A**) and CsrC (**B**) show similar protein occupancy patterns across growth (transcription start site as vertical dashed line). Novel transcription factor motifs of particular interest, ArcA for CsrB and PhoP for CsrC, are shaded. The previously confirmed (dark grey) and predicted (light grey) transcription factor binding sites are shown above by horizontal bars. The promoter sequence [-200, 10] bp surrounding the transcription start site (lowercase) used for the Miller assays are listed for CsrB (**C**) and CsrC (**D**). The transcription factor motif sequences of interest are bolded (ArcA and PhoP for CsrB and CsrC, respectively, as labeled), with the previously identified protein binding sites italicized or underlined. Rationally designed mutants (Mut) for each motif are provided. **E)** The wildtype and mutant (Mut) promoters for CsrB (left) and CsrC (right) were tested, binned on the x-axis, in both the wildtype strain (grey) and respective transcription factor knockout strains (blue, TF KO) using kinetic Miller Assays. Plotted values are from mid-logarithmic phase in Glu medium for CsrB and Cas medium for CsrC. Significance by unpaired two-tail t-test for four biological replicates performed on two separate days (designated by shape), p-values as indicated: * <0.01, ** <0.005, *** <0.001, **** <0.0005. **F)** Updated sRNA networks for CsrB and CsrC, with novel transcription factors, ArcA and PhoP, added. Created in BioRender.com.

Upon screening via plasmid Miller assays (**Figure 7B-E**), both ArcA and CytR increased transcription of the CsrB promoter in Glu medium, while PhoP increased transcription of the CsrC promoter in both Glu and Cas media. Genomic Miller assays involving promoters in the context of wildtype and mutated DNA transcription factor binding sequences of CsrB-ArcA and CsrC-PhoP (**Figure 9C-D**) further support functional regulation by direct protein binding (**Figure 9E**). Both mutant promoters result in reduced KMUs, like their respective wildtype promoter KMUs in the transcription factor knockout strains. Notably, the same behavior was measured in Glu and RDM media for CsrC-PhoP (**Table 2**, **Table S5**). Collectively, these data confirm the novel transcription factors for the CsrA sponges, ArcA for CsrB and PhoP for CsrC, supporting divergent transcriptional regulation and thus functionality (**Figure 9F**). This suggests that the sequestering function of these sponges may vary with the environment.

It is also important to note that in each case there is an orphan peak, showing similar condition dependent dynamics, upstream of each transcription start site, indicating the presence of a protein of unknown identity which may bind and act as a transcriptional regulator of both *csrB* and csrC.

No sequence motifs in this study were found within the peaks in either the CsrB or CsrC promoters. A previous ChIP-exo study has measured UvrY binding overlapping this region (Figure 2 of (81)), but direct binding specifically attributable to the orphan occupancy peak, apart from the confirmed upstream UvrY binding sites, could not be established in gel shift assays (81). UvrY does not have a sequence motif reported in the motif databases used in this study (53–55). Given the binding behavior captured by IPOD-HR, which shows mimicked signal patterns between CsrB and CsrC as well as the upstream confirmed UvrY binding sites, it is possible the orphan peaks are dependent on UvrY binding perhaps in complex with another regulator, which would explain the discrepancy between *in vitro* and *in vivo* results.

### Anticipatory responses coordinated through transcriptional regulation of sRNAs: oxidative stress regulator OxyR for amino acid balancing sRNA GcvB, and anaerobic regulator ArcA for acid responsive sRNA GadY

Our integrative pipeline for identifying new transcriptional regulatory logic for small RNAs also made strong predictions regarding the regulation of two Hfq-dependent sRNAs with large mRNA target pools: the amino acid balancing sRNA, GcvB, and the acid responsive sRNA, GadY. GcvB, referred to as an amino acid regulating sRNA (85), controls dozens of direct and indirect targets (86) including numerous transcription factors (curli regulator CsgD (87), magnesium regulator PhoP (88), and oxidative stress regulator OxyR (89)). Given its diverse functionality in numerous stress networks, the transcriptional regulation of GcvB is also likely to be more diverse beyond its known glycine-dependent regulators, GcvA and GcvR (85). Similarly, the *cis*-acting sRNA GadY, which processes the *gadXW* mRNA by facilitating RNase III cleavage, is most known for its feedback loop regulating translation of its own transcription factors GadX and GadW (90). However, GadY expression is not induced at low pH, but instead upon entry to stationary phase by the sigma factor, RpoS (75, 90). Alongside its *cis*-regulatory role, GadY is also found in high abundance in complex with other mRNAs and the chaperone protein, Hfq (91–93). These studies suggest GadY is also a *trans*-acting sRNA, regulating a widespread global network of mRNAs beyond *gadXW* processing, and thus requiring complex transcriptional regulation.

From our RNA-seq results, both GcvB and GadY show growth-dependent (**Figure 4D**) and media-dependent regulation (**Figure 4E**), albeit for likely different reasons. While GcvB is not found in any EPODs, GadY is found within EPODs for all of the Glu medium growth stages as well as early stages in Cas medium. Heterochromatin-like transcriptional silencing could explain the higher expression of GadY in the Cas medium compared to Glu medium during stationary phase. However, the lack of EPODs in Cas medium at later growth stages permits transcriptional regulation by more classic local regulators, perhaps alongside the RpoS-dependent GadY promoter. We hypothesized that GcvB and GadY have additional transcription factor regulation given these observed protein dynamics and widespread regulatory interactions.

The protein occupancies for the promoters of GcvB (**Figure 10A**) and GadY (**Figure 10B**) further highlight the sRNAs’ transcriptional regulatory differences, and the likely presence of local regulatory logic acting at their promoters. The GcvB promoter contains strong discrete protein occupancy peaks ∼50-100 bp upstream of the transcription start site, notably of distinct shape (*i.e.,* double peak at mid logarithmic phase in Cas medium compared to the broad single peak present in stationary and late stationary growth phases in both media, likely indicating transcription factor binding rather than heterochromatinization). The GadY promoter, on the other hand, contains broad peaks during conditions of EPODs (*e.g.*, late stationary phase in Glu medium) and discrete peaks in Cas medium when EPODs are not present. The most prominent peak is likely from the known GadX and GadW binding sites (as annotated by horizontal bars above the plots), although a shelf off this dominant peak near the transcription start site suggests an additional regulator is also present. From our bioinformatic analysis, we predicted two OxyR binding site motifs for GcvB and an ArcA binding site motif for GadY (shaded).

**Figure 10.**
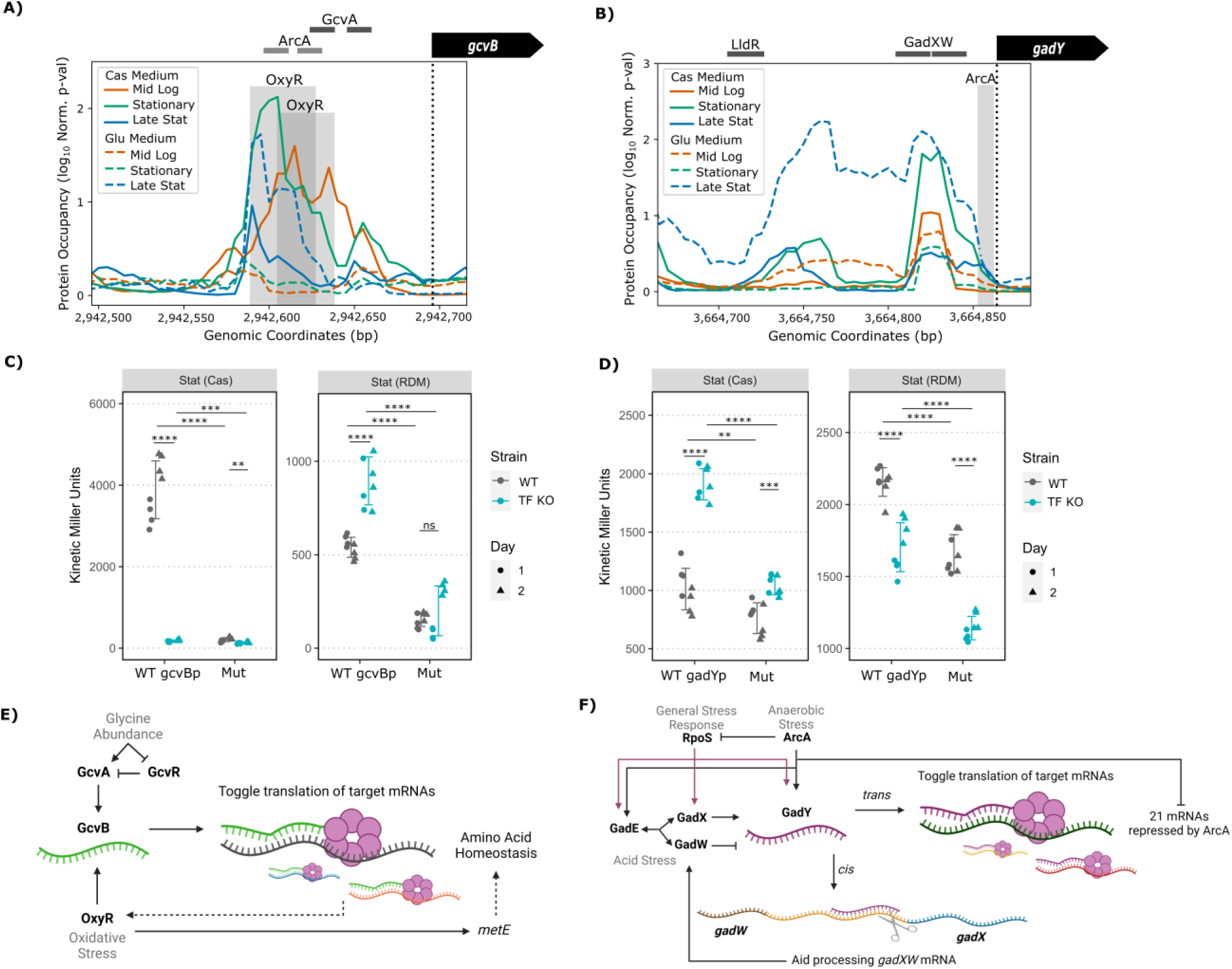
Novel anticipatory stress response connections through transcription factor regulation of sRNAs: oxidative stress and amino acid balance by GcvB, and anaerobic and acid stress by GadY. **A-B)** IPOD-HR protein occupancy (log_10_ p-values of rz-scores) at promoters of sRNAs GcvB **(A)** and GadY **(B)** show differing protein occupancy between media (transcription start site annotated as vertical dashed line). Novel transcription factor motifs of particular interest, OxyR for GcvB and ArcA for GadY, are shaded. The previously confirmed (dark grey) and predicted (light grey) transcription factor binding sites are shown above by horizontal bars. **C-D)** The wildtype and mutant (Mut) promoters for GcvB **(C)** and GadY **(D)** were tested, binned on the x-axis, in both the wildtype strain (grey) and respective transcription factor knockout strains (blue, TF KO) using genomic Miller Assays. Plotted values are from stationary phase in Cas (left) and RDM (right) media. Significance by unpaired two-tail t-test for four biological replicates performed on two separate days (designated by shape), p-values as indicated: * <0.01, ** <0.005, *** <0.001, **** <0.0005. **E-F)** Updated sRNA networks for GcvB **(E)** and GadY **(F)**, with novel transcription factors, OxyR and ArcA, respectively added. Created in BioRender.com.

Testing by genomic Miller assays revealed the media-dependent regulation of GcvB by OxyR (**Figure 10C**) and GadY by ArcA (**Figure 10D**). Tested in both Cas (left) and RDM (right) media, GcvB and GadY promoters show different effects when glucose is present (RDM) or absent (Cas). For GcvB, in the absence of glucose, transcription measured as KMUs is drastically induced in the wildtype strain (grey) that is strikingly diminished in Δ*oxyR* (blue) and when the promoter is rationally mutated (Mut). Alternatively, in the RDM medium, when glucose is present, the wildtype promoter shows moderate expression that is slightly increased in Δ*oxyR* as opposed to decreased by the mutant promoter. These results suggest interplay between OxyR and the known GcvA and GcvR regulators of GcvB in a glucose- and amino acid-dependent manner. A similar result is found for GadY, albeit in the opposite media: RDM medium shows activation in both *ΔarcA* and the mutated promoter, while the Cas medium lacking glucose shows decreased expression in *ΔarcA* as opposed to slight activation effect by the mutant promoter. Again, this media-dependent effect supports interplay between ArcA, RpoS, and the acid regulators GadX and GadW in transcriptionally regulating GadY.

For GcvB, the interplay between amino acid balance and oxidative stress networks (**Figure 10E**) could serve an anticipatory role (94) where oxidative stress unfolds proteins that are then degraded, causing an imbalance in free amino acids needed to replace the unfolded proteins. Similarly, for GadY, the interplay between acid and anaerobic stress networks (**Figure 10F**) along with the general stress responder RpoS, could serve an anticipatory role where anaerobic respiration produces carboxylic acids and thus decreases pH, particularly when nutrients are scarce later in growth.

## DISCUSSION

Proper cellular responses to nutrient stress conditions require an intricate interplay of many regulators acting at both transcriptional and post-transcriptional levels. In this study, we used a combination of protein-DNA interaction and RNA transcript abundance measurements to study the response of *E. coli* to a broad range of nutrient conditions. Consistent with prior findings, we observed the presence of heterochromatin-like regions of extended protein occupancy that silence hundreds of loci throughout the chromosome; furthermore, we were able to identify several additional cases of dynamic EPODs where nutrient stresses increase the accessibility of some normally-silenced loci and permit transcription. Our RNA-seq data also reveal that small RNAs play a particularly prominent role in the adaptation of *E.* coli to nutrient stress conditions. The majority of DNA regions encoding small regulatory RNAs (sRNAs) show dynamic protein occupancy of small peaks predominantly outside of EPODs, indicative of differential expression by local regulators. We were able to use a novel bioinformatic pipeline to predict many additional sRNA-transcription factor regulatory interactions on the basis of our data, with the vast majority of tested cases confirmed in follow-up experiments.

### Glucose and amino acid starvation activate transcription of multiple stress networks

Using two different carbon sources, we anticipated that Cas medium would induce a glucose starvation, or general stress, response (2, 3) while Glu medium would induce an amino acid starvation, or stringent, response (4–6). The RNA-seq expression results, analyzed for enrichment of Gene Ontology terms, agreed with these hypotheses (**Figure 1C**). Specifically, in the case of the Glu medium, we observed induction of amino acid biosynthesis early in growth due to its lack of exogenous amino acids. Later in growth, both media show reduction in amino acid synthesis and translation, agreeing with prior literature that have found ribosome synthesis to be downregulated during nutrient starvation (58, 59). As for glucose starvation, the Cas medium induced the glyoxylate cycle (an alteration to the TCA cycle that conserves carbon). By stationary phase, Cas medium spurs aerobic respiration, suggesting the cells have shifted to maximize energy extraction per unit carbon. Our analysis also found activation of other stress networks, including iron, oxidative, and pH stresses (**Figure 1C-D**), supporting interconnected stress network activation alongside shifts in carbon metabolism across growth phases.

### Dynamic protein occupancies suggest coordination between nucleoid-associated proteins and transcription factors to regulate gene expression throughout growth

From a global perspective, we were able to observe the presence of both static and dynamic EPODs, with the former representing cases that never opened to allow expression under the tested nutrient stresses, and the latter indicating regions that are normally heterochromatinized but open to allow transcription under specific conditions. Interestingly, the Glu medium had a much higher number of static EPODs than Cas (233 to 160, **Figure S3**), which agreed with observations of higher nucleoid-associated protein activity overall in the Glu medium throughout growth. Gene ontology enrichment of genes within these static EPODs suggest predominant regulatory roles in lipopolysaccharide and amino acid homeostasis in both media, with the Cas conditions also regulating iron and phosphonate homeostasis and Glu medium cells regulating cell adhesion and curli formation through chromatin structure (**Figure 1E**). From the lipopolysaccharide biosynthesis operons example (**Figure 2** and provided examples of *fecIRA* and *yehEDCBA* oprons in **Figure S4** and **Figure S5**) it appears the static EPODs predominantly repress transcription via nucleoid-associated proteins blocking or stalling RNAP as previously postulated (11, 14, 15, 61).

While static EPODs in the examples appear to function as silencers of internal promoters (likely preventing productive transcriptional elongation), agreeing with prior literature in particular for the H-NS nucleoid-associated protein (11, 13–15, 95), they can also co-exist with more classical *cis*-regulatory logic. For example, the disappearance of EPODs does not necessarily result in induced expression (as exemplified by the *yehDCBA* operon in **Figure S5**). Indeed, decoupling of nucleoid-associated protein binding and RNAP activity has been previously observed for Dps (96). While prior IPOD-HR datasets do not show Dps binding at these DNA regions (**Figure S10**, data from (11)), these EPOD examples suggest a more complex model. Building on prior hypotheses in literature, we postulate that release of DNA by nucleoid-associated proteins alone does not necessarily result in activated transcription. Dynamic protein occupancy regions, such as the dynamic EPOD example surrounding the acid response operons in **Figure 3**, likely rely on a concerted balance between nucleoid-associated proteins and transcription factors to regulate transcription depending on the environmental condition. EPODs will silence genes under the conditions when they are present, but more conventional transcriptional regulatory logic is still active even when a dynamic EPOD dissociates at a particular site.

### Dynamic sRNA expression is controlled by numerous transcription factors in response to nutrient stress

We also identified a strong enrichment of sRNAs among the genes induced by our nutrient stress conditions (particularly the onset of stationary phase). As shown in **Figure 4B**, only a few sRNAs are contained within EPODs throughout growth, such RyeG and other toxic sRNAs, consistent with the role of static EPODs (or regions of dense nucleoid-associated protein binding) in silencing potentially harmful horizontally acquired genes (11). While they are mostly not subject to regulation by large scale chromatin structure, many sRNAs do show dynamic protein occupancies locally near their promoters, suggesting transcription factor binding events. Transcription factor regulation of sRNAs is further supported by their measured dynamic expression across growth phases and between media (illustrated by the heatmaps in **Figure 4C** and **Figure S7**). Particularly during the transition from logarithmic to stationary phase, cells reduce replication as nutrients become scarce, relying on post-transcriptional regulation by sRNAs to toggle both the translation and stability of existing mRNAs (19, 24, 97). Media-dependent sRNA expression was also dramatic (**Figure 4E**) in which expression in Cas medium was significantly higher than expression in Glu medium for many sRNAs (mid logarithmic: 23, stationary: 43, late stationary: 27). Given the predominant growth and media-dependent expression alongside the dynamic non-EPOD protein occupancy around sRNA promoters, sRNAs likely rely on numerous cooperative and competitive transcription factors to result in such dynamic expression patterns during glucose and amino acid starvation.

We therefore combined our IPOD-HR and RNA-seq results to bioinformatically predict, and then experimentally test novel transcription factors regulating sRNAs. The plasmid-based screening via Miller assays resulted in 36 novel sRNA-transcription factor relationships for future focused investigations. New regulatory interactions for five important sRNAs for carbon balance in *E. coli*, IsrB, CsrB and CsrC, GcvB, and GadY were confirmed using binding site mutations to test direct functional regulation by the identified transcription factors (FNR, NarP, PhoP for IsrB, ArcA for CsrB, PhoP for CsrC, OxyR for GcvB, and ArcA for GadY). Interestingly four key findings were observed that confirm the complex dynamics of sRNA transcriptional regulation. In all cases, the results emphasize the value of integrated IPOD-HR, RNA-seq, and bioinformatics to piece together overlapping binding sites of transcription factors that control dynamic promoters.

In the case of IsrB, we confirmed three novel transcription factors that complete its aerobic metabolism regulatory network (**Figure 5F**). During periods of anaerobic stress, FNR and NarP repress the transcription of IsrB. A prior ChIP-seq study measured FNR binding near the promoter of IsrB (98). Since FNR also transcriptionally actives the sRNA, FnrS (99), that represses IsrB translation of AzuC, FNR effectively represses IsrB translation in addition to repressing its transcription. As an aside, the transcription factors FNR and CRP have been previously shown to coregulate targets throughout the genome (98), with their binding sites for regulating IsrB completely overlapping. NarP balances nitrate and nitrite levels prevalent during oxygen deprivation (100), interestingly with itself a target of multiple other sRNAs (101). NarP thus provides redundant repression of IsrB with FNR in anaerobic conditions. Finally, PhoP is part of the two-component system PhoQP, in which limited extracellular divalent cations activate DNA binding (102). PhoP activity is also induced during low pH through the acid response two-component system, EvgAS. (103–105). Curiously, there are conflicting reports of IsrB induction (66) and repression (51, 75) in low pH settings. It is hypothesized that this is due to the overlapping transcription factor binding sites, where PhoP binding may be dependent on other protein binding events, such as FNR and NarP, for adequate DNA accessibility.

In the case of the sRNAs, CsrB and CsrC, we find that they are independently regulated for proper control of their shared target: the global RNA regulatory protein CsrA. Beyond metabolism, the Csr system and its homologs in other organisms have been widely implicated in pathogenesis, regulating biofilm formation, motility, and secretion systems (106), yet no pathogenically-relevant transcription factors of CsrA, CsrB, or CsrC have been identified in *E. coli*. Here, we confirmed two separate transcriptional activators, ArcA for CsrB and PhoP for CsrC, suggesting divergent expression conditions to sequester CsrA as illustrated in **Figure 9F**. ArcA is part of the two-component ArcAB system, in which ArcA is activated in microaerobic and anaerobic conditions depending on the redox state of the cell (107). While CsrB expression has not been shown to be induced in strict anaerobic conditions, ArcA may rely on neighboring transcription factor binding events like UvrY and IHF for CsrB induction. Indeed, a ChIP-chip experiment found ArcA binds the promoter of CsrB in nitrosative stress but not in anaerobic conditions alone (108). Secondly, PhoP is part of a two-component PhoQP system that is activated during divalent cation starvation and hyperosmotic stress (102). PhoP controls magnesium and calcium homeostasis, acid resistance, cell membrane composition, among other gene networks. Indeed, an independent RNA-seq dataset found CsrC induction under magnesium starvation (109). CsrA is known to repress translation of EvgA (23, 110), thus PhoP regulation of CsrC provides a feedback loop for the EvgA regulon through the carbon regulatory system. Importantly, PhoP and ArcA both have been implicated in pathogenic regulation in *E. coli* (111, 112), partially explaining the Csr system’s pathogenic utility (106). By having different sets of transcription factors driving their expression, independent titration of CsrA by CsrB and CsrC could be beneficial during dynamic environments, such as when colonizing the human gut where oxygen is scarce and nutrient availability fluctuates.

The last two sRNAs with confirmed novel direct transcription factor regulation are two global sRNAs with large sets of mRNA targets: GcvB (85) and GadY (90, 113). Our study successfully confirmed the direct transcriptional activation of the GcvB sRNA by OxyR and of the GadY sRNA by ArcA, bridging distinct stress networks through these global sRNAs. For the novel OxyR regulator of GcvB expression, recent studies have pointed to the connection of oxidative stress with amino acid balance (89, 114). More directly, a study using a GcvB knockout strain found that GcvB increases *oxyR* translation, albeit direct interaction between sRNA and mRNA was not investigated (89). These relationships support the interplay between the amino acid and oxidative stress networks. GcvB could serve an anticipatory role (94), where oxidative stress degrades many proteins, leading to amino acid imbalances. Prior to glycine abundance and thus GcvA activation, OxyR could stimulate GcvB expression to more quickly respond to the inevitable amino acid stress (**Figure 10E**).

In the case of the sRNA GadY, we found two ArcA motifs within the [-200, 10] bp promoter region of GadY. However, the ArcA motif found nearest the transcription start site was the only one confirmed in this study as we focused only on binding sites within the [-100, 10] bp window. Global studies have found that ArcA directly activates *gadWX* and *gadE* transcription. Given the antisense regulation of *gadWX* by GadY, it is logical that GadY expression is also induced by ArcA to enable translation of GadW and GadX (**Figure 10F**). Furthermore, prior Hfq co-immunoprecipitation studies found GadY binds 21 different mRNAs that are known to be transcriptionally repressed by ArcA (91–93). Therefore, upon ArcA activation, such as during microaerobic conditions, ArcA will repress transcription of these mRNAs while inducing expression of GadY. GadY then likely represses the translation of the many already transcribed mRNAs whose transcription have been halted by ArcA. In this way, GadY could provide a feedback loop with the ArcA regulon by turning off translation of mRNAs until they can be properly degraded (although the repression of the 21 mRNAs by GadY needs experimental confirmation). Thus, our findings bolster the connection between the anaerobic and acid stress networks, through regulation of the sRNA GadY by transcription factor ArcA with GadW and GadX. GadY could serve to mediate between these stress networks, serving an anticipatory role such as spurring translation of the acid regulators GadW and GadX during microaerobic conditions, forecasting acid accumulation by anaerobic respiration.

## CONCLUSION

This study expands the complex transcriptional and post-transcriptional regulation model of *E. coli* during nutrient stress: we observe an intricate interplay between dynamic heterochromatin-based regulation in some cases, alongside local transcription factor regulation (both by well-studied and newly-identified interactions), with the latter also driving dynamic sRNA expression and post-transcriptional regulation. Static and dynamic regions of extended protein occupancy domains (EPODs), typically consisting of nucleoid-associated proteins and bearing striking similarities to eukaryotic heterochromatin, were identified by *in vivo* protein occupancy display – high resolution (IPOD-HR). Amino acid starvation (Glu) media had generally higher protein occupancy in these EPODs across growth phases, suggesting an increased dependence on DNA condensation and thus transcriptional silencing compared to glucose starvation. We found a major dependence on sRNA-mediated regulation during the entry into stationary phase and were able to leverage our IPOD-HR protein occupancy data alongside bioinformatic predictions to reveal a multitude of novel transcription factor-sRNA regulatory interactions, with experimental follow-up confirming new instances of direct transcriptional regulation for five sRNAs of importance in carbon metabolism: IsrB, CsrB and CsrC, GcvB, and GadY. The five sRNAs highlighted above emphasize the complex transcription factor regulation of sRNAs for tailored expression in response to changing glucose and amino acid availability. Additionally, the GadY example highlights the potential coordination between nucleoid-associated proteins and transcription factors to dynamically control sRNA expression across growth phases. While providing a limited set of examples, this work supports the likelihood of complex transcriptional regulation of sRNAs in response to stress throughout bacteria, both locally by transcription factors and globally by EPODs composed of nucleoid-associated proteins. We expect that a similar strategy, combining global protein occupancy profiling to identify the locations of protein binding, RNA-seq to identify differentially expressed genes, and bioinformatic analysis to determine the sequence motifs linking them, will be a powerful approach for characterizing complex regulatory responses in other contexts and organisms.

## SUPPLEMENTARY DATA

Supplementary Data are available at NAR online.

File S1. Combined supplemental figures.

Table S1. Details of strains, plasmids, primers used and sRNAs studied.

Table S2. RNA-seq results, including TPM, differential expression analysis, and Gene Ontology analysis.

Table S3. Protein occupancy results, including EPOD regions, Gene Ontology analysis of EPODs, sRNAs contained in EPODs, and RNA polymerase occupancy at promoters.

Table S4. Bioinformatic analysis results, including protein occupancy peaks identified, motifs found in the transcription-altering peaks, and external RNA-seq support details.

Table S5. Kinetic Miller assay results, including plasmid-based results in stress (Glu and Cas) and rich (RDM) media, control plasmid results, and genomic-based mutational reporter results.

Table S6. Example growth curves for all media with sample time points, slow growing strains, and pathlength optical density correction indicated.

## AUTHOR CONTRIBUTIONS

Conceptualization, A.M.E., L.M.C and P.L.F.; Methodology, A.M.E. and P.L.F.; Investigation, A.M.E, T.J., S.J., J.K.; Writing – Original Draft, A.M.E.; Writing – Review & Editing, A.M.E., L.M.C. and P.L.F; Funding Acquisition, S.T., L.M.C. and P.L.F.; Resources, S.T, L.M.C., and P.L.F.; Supervision, L.M.C. and P.L.F.

## ACKNOWLEDGMENTS

We would like to thank the Wade Lab (University of Albany, Albany, NY) for their sharing of select *E. coli* transcription factor knockout strains (K-12 MG1655 Δ*crp,* Δ*oxyR,* Δ*ulaR*). We would also like to thank the Pfleger Lab (University of Wisconsin, Madison, WI) for sharing their CRISPR recombineering system to improve the efficiency of our lambda-red genomic insertions. Finally, we thank Dr. Mia Mihailovic for her intellectual support in this work.

## REFERENCES

1. Conway, T., Krogfelt, K.A. and Cohen, P.S. (2004) The Life of Commensal *Escherichia coli* in the Mammalian Intestine. EcoSal Plus, 1.

2. Bouillet, S., Bauer, T.S. and Gottesman, S. (2024) RpoS and the bacterial general stress response. Microbiology and Molecular Biology Reviews, 88, e00151–22.

3. Handler, S. and Kirkpatrick, C.L. (2024) New layers of regulation of the general stress response sigma factor RpoS. Front Microbiol, 15, 1363955.

4. Zhu, M. and Dai, X. (2023) Stringent response ensures the timely adaptation of bacterial growth to nutrient downshift. Nat Commun, 14, 467.

5. Irving, S.E., Choudhury, N.R. and Corrigan, R.M. (2021) The stringent response and physiological roles of (pp)pGpp in bacteria. Nat Rev Microbiol, 19, 256–271.

6. Miyakoshi, M. (2024) Multilayered regulation of amino acid metabolism in *Escherichia coli*. Curr Opin Microbiol, 77, 102406.

7. Hussein, R. and Lim, H.N. (2012) Direct comparison of small RNA and transcription factor signaling. Nucleic Acids Res, 40, 7269–7279.

8. Hołówka, J. and Zakrzewska-Czerwińska, J. (2020) Nucleoid Associated Proteins: The Small Organizers That Help to Cope With Stress. Front Microbiol, 11.

9. Vora, T., Hottes, A.K. and Tavazoie, S. (2009) Protein Occupancy Landscape of a Bacterial Genome. Mol Cell, 35, 247–253.

10. Freddolino, P.L., Amemiya, H.M., Goss, T.J. and Tavazoie, S. (2021) Dynamic landscape of protein occupancy across the *Escherichia coli* chromosome. PLoS Biol, 19, e3001306.

11. Amemiya, H.M., Goss, T.J., Nye, T.M., Hurto, R.L., Simmons, L.A. and Freddolino, P.L. (2022) Distinct heterochromatin-like domains promote transcriptional memory and silence parasitic genetic elements in bacteria. EMBO J, 41.

12. Rakibova, Y., Dunham, D.T., Seed, K.D. and Freddolino, P.L. (2024) Nucleoid-associated proteins shape the global protein occupancy and transcriptional landscape of a clinical isolate of Vibrio cholerae. bioRxiv doi: 10.1101/2023.12.30.573743, 25 March 2024, pre- print: not peer-reviewed.

13. Singh, S.S., Singh, N., Bonocora, R.P., Fitzgerald, D.M., Wade, J.T. and Grainger, D.C. (2014) Widespread suppression of intragenic transcription initiation by H-NS. Genes Dev, 28, 214– 219.

14. Boudreau, B.A., Hron, D.R., Qin, L., van der Valk, R.A., Kotlajich, M. V, Dame, R.T. and Landick, R. (2018) StpA and Hha stimulate pausing by RNA polymerase by promoting DNA–DNA bridging of H-NS filaments. Nucleic Acids Res, 46, 5525–5546.

15. Kotlajich, M. V, Hron, D.R., Boudreau, B.A., Sun, Z., Lyubchenko, Y.L. and Landick, R. (2015) Bridged filaments of histone-like nucleoid structuring protein pause RNA polymerase and aid termination in bacteria. eLife, 4, e04970.

16. Gupta, A., Joshi, A., Arora, K., Mukhopadhyay, S. and Guptasarma, P. (2023) The bacterial nucleoid-associated proteins, HU and Dps, condense DNA into context-dependent biphasic or multiphasic complex coacervates. Journal of Biological Chemistry, 299, 104637.

17. Van Assche, E., Van Puyvelde, S., Vanderleyden, J. and Steenackers, H.P. (2015) RNA-binding proteins involved in post-transcriptional regulation in bacteria. Front Microbiol, 6.

18. Christopoulou, N. and Granneman, S. (2022) The role of RNA-binding proteins in mediating adaptive responses in Gram-positive bacteria. FEBS J, 289, 1746–1764.

19. Jørgensen, M.G., Pettersen, J.S. and Kallipolitis, B.H. (2020) sRNA-mediated control in bacteria: An increasing diversity of regulatory mechanisms. Biochimica et Biophysica Acta (BBA) - Gene Regulatory Mechanisms, 1863, 194504.

20. Papenfort, K. and Melamed, S. (2023) Small RNAs, Large Networks: Posttranscriptional Regulons in Gram-Negative Bacteria. Annu Rev Microbiol, 77, 23–43.

21. Leistra, A.N., Curtis, N.C. and Contreras, L.M. (2019) Regulatory non-coding sRNAs in bacterial metabolic pathway engineering. Metab Eng, 52, 190–214.

22. Villa, J.K., Su, Y., Contreras, L.M. and Hammond, M.C. (2018) Synthetic Biology of Small RNAs and Riboswitches. Microbiol Spectr, 6.

23. Leistra, A.N., Gelderman, G., Sowa, S.W., Moon-Walker, A., Salis, H.M. and Contreras, L.M. (2018) A Canonical Biophysical Model of the CsrA Global Regulator Suggests Flexible Regulator-Target Interactions. Sci Rep, 8, 9892.

24. Hör, J., Matera, G., Vogel, J., Gottesman, S. and Storz, G. (2020) Trans-Acting Small RNAs and Their Effects on Gene Expression in *Escherichia coli* and *Salmonella enterica*. EcoSal Plus, 9.

25. Wagner, E.G.H. and Romby, P. (2015) Small RNAs in Bacteria and Archaea: Who They Are, What They Do, and How They Do It. In Friedmann, T., Dunlap, J., Goodwin, S. (eds), Advances in Genetics. Academic Press, Vol. 90, pp. 133–208.

26. Modi, S.R., Camacho, D.M., Kohanski, M.A., Walker, G.C. and Collins, J.J. (2011) Functional characterization of bacterial sRNAs using a network biology approach. Proceedings of the National Academy of Sciences, 108, 15522–15527.

27. Brosse, A. and Guillier, M. (2018) Bacterial Small RNAs in Mixed Regulatory Networks. Microbiol Spectr, 6.

28. Arrieta-Ortiz, M.L., Hafemeister, C., Shuster, B., Baliga, N.S., Bonneau, R. and Eichenberger, P. (2020) Inference of Bacterial Small RNA Regulatory Networks and Integration with Transcription Factor-Driven Regulatory Networks. mSystems, 5.

29. Mandin, P. and Gottesman, S. (2010) Integrating anaerobic/aerobic sensing and the general stress response through the ArcZ small RNA. EMBO J, 29, 3094–3107.

30. Majdalani, N., Cunning, C., Sledjeski, D., Elliott, T. and Gottesman, S. (1998) DsrA RNA regulates translation of RpoS message by an anti-antisense mechanism, independent of its action as an antisilencer of transcription. Proceedings of the National Academy of Sciences, 95, 12462– 12467.

31. Zhang, A., Altuvia, S., Tiwari, A., Argaman, L., Hengge-Aronis, R. and Storz, G. (1998) The OxyS regulatory RNA represses *rpoS* translation and binds the Hfq (HF-I) protein. EMBO J, 17, 6061–6068.

32. Kim, W. and Lee, Y. (2020) Mechanism for coordinate regulation of *rpoS* by sRNA-sRNA interaction in *Escherichia coli*. RNA Biol, 17, 176–187.

33. Majdalani, N., Hernandez, D. and Gottesman, S. (2002) Regulation and mode of action of the second small RNA activator of RpoS translation, RprA. Mol Microbiol, 46, 813–826.

34. Seo, S.W., Gao, Y., Kim, D., Szubin, R., Yang, J., Cho, B.-K. and Palsson, B.O. (2017) Revealing genome-scale transcriptional regulatory landscape of OmpR highlights its expanded regulatory roles under osmotic stress in *Escherichia coli* K-12 MG1655. Sci Rep, 7, 2181.

35. Chubiz, L.M. and Rao, C. V. (2011) Role of the *mar-sox-rob* Regulon in Regulating Outer Membrane Porin Expression. J Bacteriol, 193, 2252–2260.

36. Ferrario, M., Ernsting, B.R., Borst, D.W., Wiese, D.E., Blumenthal, R.M. and Matthews, R.G. (1995) The leucine-responsive regulatory protein of *Escherichia coli* negatively regulates transcription of *ompC* and *micF* and positively regulates translation of *ompF*. J Bacteriol, 177, 103–113.

37. Huang, L., Tsui, P. and Freundlich, M. (1990) Integration host factor is a negative effector of *in vivo* and *in vitro* expression of *ompC* in *Escherichia coli*. J Bacteriol, 172, 5293–5298.

38. Delihas, N. and Forst, S. (2001) MicF: an antisense RNA gene involved in response of *Escherichia coli* to global stress factors. J Mol Biol, 313, 1–12.

39. Baba, T., Ara, T., Hasegawa, M., Takai, Y., Okumura, Y., Baba, M., Datsenko, K.A., Tomita, M., Wanner, B.L. and Mori, H. (2006) Construction of *Escherichia coli* K-12 in-frame, single-gene knockout mutants: the Keio collection. Mol Syst Biol, 2.

40. Cherepanov, P.P. and Wackernagel, W. (1995) Gene disruption in *Escherichia coli*: TcR and KmR cassettes with the option of Flp-catalyzed excision of the antibiotic-resistance determinant. Gene, 158, 9–14.

41. Freddolino, P.L., Amini, S. and Tavazoie, S. (2012) Newly Identified Genetic Variations in Common *Escherichia coli* MG1655 Stock Cultures. J Bacteriol, 194, 303–306.

42. Neidhardt, F.C., Bloch, P.L. and Smith, D.F. (1974) Culture Medium for Enterobacteria. J Bacteriol, 119, 736–747.

43. Quinlan, A.R. and Hall, I.M. (2010) BEDTools: a flexible suite of utilities for comparing genomic features. Bioinformatics, 26, 841–842.

44. Li, H., Handsaker, B., Wysoker, A., Fennell, T., Ruan, J., Homer, N., Marth, G., Abecasis, G. and Durbin, R. (2009) The Sequence Alignment/Map format and SAMtools. Bioinformatics, 25, 2078–2079.

45. Putri, G.H., Anders, S., Pyl, P.T., Pimanda, J.E. and Zanini, F. (2022) Analysing high-throughput sequencing data in Python with HTSeq 2.0. Bioinformatics, 38, 2943–2945.

46. Love, M.I., Huber, W. and Anders, S. (2014) Moderated estimation of fold change and dispersion for RNA-seq data with DESeq2. Genome Biol, 15, 550.

47. Stephens, M. (2016) False discovery rates: a new deal. Biostatistics, 18, 275–294.

48. Sherman, B.T., Hao, M., Qiu, J., Jiao, X., Baseler, M.W., Lane, H.C., Imamichi, T. and Chang, W. (2022) DAVID: a web server for functional enrichment analysis and functional annotation of gene lists (2021 update). Nucleic Acids Res, 50, W216–W221.

49. Huang, D.W., Sherman, B.T. and Lempicki, R.A. (2009) Systematic and integrative analysis of large gene lists using DAVID bioinformatics resources. Nat Protoc, 4, 44–57.

50. Hosack, D.A., Dennis, G., Sherman, B.T., Lane, H.C. and Lempicki, R.A. (2003) Identifying biological themes within lists of genes with EASE. Genome Biol, 4, R70.

51. Mihailovic, M.K., Ekdahl, A.M., Chen, A., Leistra, A.N., Li, B., González Martínez, J., Law, M., Ejindu, C., Massé, É., Freddolino, P.L., et al. (2021) Uncovering Transcriptional Regulators and Targets of sRNAs Using an Integrative Data-Mining Approach: H-NS-Regulated RseX as a Case Study. Front Cell Infect Microbiol, 11.

52. Grant, C.E., Bailey, T.L. and Noble, W.S. (2011) FIMO: scanning for occurrences of a given motif. Bioinformatics, 27, 1017–1018.

53. Robison, K., McGuire, A.M. and Church, G.M. (1998) A comprehensive library of DNA-binding site matrices for 55 proteins applied to the complete *Escherichia coli* K-12 genome. J Mol Biol, 284, 241–254.

54. Pachkov, M., Balwierz, P.J., Arnold, P., Ozonov, E. and van Nimwegen, E. (2012) SwissRegulon, a database of genome-wide annotations of regulatory sites: recent updates. Nucleic Acids Res, 41, D214–D220.

55. Eckweiler, D., Dudek, C.-A., Hartlich, J., Brötje, D. and Jahn, D. (2018) PRODORIC2: the bacterial gene regulation database in 2018. Nucleic Acids Res, 46, D320–D326.

56. Pothoulakis, G., Ceroni, F., Reeve, B. and Ellis, T. (2014) The Spinach RNA Aptamer as a Characterization Tool for Synthetic Biology. ACS Synth Biol, 3, 182–187.

57. Mehrer, C.R., Incha, M.R., Politz, M.C. and Pfleger, B.F. (2018) Anaerobic production of medium-chain fatty alcohols via a β-reduction pathway. Metab Eng, 48, 63–71.

58. Li, S.H.-J., Li, Z., Park, J.O., King, C.G., Rabinowitz, J.D., Wingreen, N.S. and Gitai, Z. (2018) *Escherichia coli* translation strategies differ across carbon, nitrogen and phosphorus limitation conditions. Nat Microbiol, 3, 939–947.

59. Njenga, R., Boele, J., Öztürk, Y. and Koch, H.-G. (2023) Coping with stress: How bacteria fine-tune protein synthesis and protein transport. Journal of Biological Chemistry, 299, 105163.

60. Salgado, H., Gama-Castro, S., Lara, P., Mejia-Almonte, C., Alarcón-Carranza, G., López-Almazo, A.G., Betancourt-Figueroa, F., Peña-Loredo, P., Alquicira-Hernández, S., Ledezma-Tejeida, D., et al. (2024) RegulonDB v12.0: a comprehensive resource of transcriptional regulation in *E. coli* K-12. Nucleic Acids Res, 52, D255–D264.

61. Amemiya, H.M., Schroeder, J. and Freddolino, P.L. (2021) Nucleoid-associated proteins shape chromatin structure and transcriptional regulation across the bacterial kingdom. Transcription, 12, 182–218.

62. Lalaouna, D., Prévost, K., Laliberté, G., Houé, V. and Massé, E. (2018) Contrasting silencing mechanisms of the same target mRNA by two regulatory RNAs in *Escherichia coli*. Nucleic Acids Res, 46, 2600–2612.

63. Aiso, T., Kamiya, S., Yonezawa, H. and Gamou, S. (2014) Overexpression of an antisense RNA, ArrS, increases the acid resistance of Escherichia coli. Microbiology (N Y), 160, 954–961.

64. Raina, M., Aoyama, J.J., Bhatt, S., Paul, B.J., Zhang, A., Updegrove, T.B., Miranda-Ríos, J. and Storz, G. (2022) Dual-function AzuCR RNA modulates carbon metabolism. Proceedings of the National Academy of Sciences, 119.

65. Chen, S., Lesnik, E.A., Hall, T.A., Sampath, R., Griffey, R.H., Ecker, D.J. and Blyn, L.B. (2002) A bioinformatics based approach to discover small RNA genes in the *Escherichia coli* genome. Biosystems, 65, 157–177.

66. Hemm, M.R., Paul, B.J., Miranda-Ríos, J., Zhang, A., Soltanzad, N. and Storz, G. (2010) Small Stress Response Proteins in *Escherichia coli*: Proteins Missed by Classical Proteomic Studies. J Bacteriol, 192, 46–58.

67. Kanjee, U. and Houry, W.A. (2013) Mechanisms of Acid Resistance in *Escherichia coli*. Annu Rev Microbiol, 67, 65–81.

68. Moreau, P.L. (2007) The Lysine Decarboxylase CadA Protects *Escherichia coli* Starved of Phosphate against Fermentation Acids. J Bacteriol, 189, 2249–2261.

69. Zhao, B. and Houry, W.A. (2010) Acid stress response in enteropathogenic gammaproteobacteria: an aptitude for survival. Biochemistry and Cell Biology, 88, 301–314.

70. Wilson, D.B. and Hogness, D.S. (1964) The Enzymes of the Galactose Operon in *Escherichia coli*. Journal of Biological Chemistry, 239, 2469–2481.

71. Seo, S.W., Kim, D., O’Brien, E.J., Szubin, R. and Palsson, B.O. (2015) Decoding genome-wide GadEWX-transcriptional regulatory networks reveals multifaceted cellular responses to acid stress in *Escherichia coli*. Nat Commun, 6, 7970.

72. L.B. Almeida, B., M. Bahrudeen, M.N., Chauhan, V., Dash, S., Kandavalli, V., Häkkinen, A., Lloyd-Price, J., S.D. Cristina, P., Baptista, I.S.C., Gupta, A., et al. (2022) The transcription factor network of *E. coli* steers global responses to shifts in RNAP concentration. Nucleic Acids Res, 50, 6801–6819.

73. Deter, H.S., Hossain, T. and Butzin, N.C. (2021) Antibiotic tolerance is associated with a broad and complex transcriptional response in *E. coli*. Sci Rep, 11, 6112.

74. Wassarman, K.M., Repoila, F., Rosenow, C., Storz, G. and Gottesman, S. (2001) Identification of novel small RNAs using comparative genomics and microarrays. Genes Dev, 15, 1637–1651.

75. Gao, Y., Yurkovich, J.T., Seo, S.W., Kabimoldayev, I., Dräger, A., Chen, K., Sastry, A. V, Fang, X., Mih, N., Yang, L., et al. (2018) Systematic discovery of uncharacterized transcription factors in *Escherichia coli* K-12 MG1655. Nucleic Acids Res, 10.1093/nar/gky752.

76. Bordbar, A., Nagarajan, H., Lewis, N.E., Latif, H., Ebrahim, A., Federowicz, S., Schellenberger, J. and Palsson, B.O. (2014) Minimal metabolic pathway structure is consistent with associated biomolecular interactions. Mol Syst Biol, 10.

77. Kim, D., Seo, S.W., Gao, Y., Nam, H., Guzman, G.I., Cho, B.-K. and Palsson, B.O. (2018) Systems assessment of transcriptional regulation on central carbon metabolism by Cra and CRP. Nucleic Acids Res, 46, 2901–2917.

78. Seo, S.W., Kim, D., Latif, H., O’Brien, E.J., Szubin, R. and Palsson, B.O. (2014) Deciphering Fur transcriptional regulatory network highlights its complex role beyond iron metabolism in *Escherichia coli*. Nat Commun, 5, 4910.

79. Esquerré, T., Bouvier, M., Turlan, C., Carpousis, A.J., Girbal, L. and Cocaign-Bousquet, M. (2016) The Csr system regulates genome-wide mRNA stability and transcription and thus gene expression in *Escherichia coli*. Sci Rep, 6, 25057.

80. Contreras, F.U., Camacho, M.I., Pannuri, A., Romeo, T., Alvarez, A.F. and Georgellis, D. (2023) Spatiotemporal regulation of the BarA/UvrY two-component signaling system. Journal of Biological Chemistry, 299, 104835.

81. Zere, T.R., Vakulskas, C.A., Leng, Y., Pannuri, A., Potts, A.H., Dias, R., Tang, D., Kolaczkowski, B., Georgellis, D., Ahmer, B.M.M., et al. (2015) Genomic Targets and Features of BarA-UvrY (-SirA) Signal Transduction Systems. PLoS One, 10, e0145035.

82. Rojano-Nisimura, A.M., Grismore, K.B., Ruzek, J.S., Avila, J.L. and Contreras, L.M. (2024) The Post-Transcriptional Regulatory Protein CsrA Amplifies Its Targetome through Direct Interactions with Stress-Response Regulatory Hubs: The EvgA and AcnA Cases. Microorganisms, 12, 636.

83. Rojano-Nisimura, A.M., Simmons, T.R., Leistra, A.N., Mihailovic, M.K., Buchser, R., Ekdahl, A.M., Joseph, I., Curtis, N.C. and Contreras, L.M. (2023) CsrA selectively modulates sRNA-mRNA regulator outcomes. Front Mol Biosci, 10.

84. Pannuri, A., Vakulskas, C.A., Zere, T., McGibbon, L.C., Edwards, A.N., Georgellis, D., Babitzke, P. and Romeo, T. (2016) Circuitry Linking the Catabolite Repression and Csr Global Regulatory Systems of Escherichia coli. J Bacteriol, 198, 3000–3015.

85. Urbanowski, M.L., Stauffer, L.T. and Stauffer, G. V. (2000) The *gcvB* gene encodes a small untranslated RNA involved in expression of the dipeptide and oligopeptide transport systems in *Escherichia coli*. Mol Microbiol, 37, 856–868.

86. Miyakoshi, M., Okayama, H., Lejars, M., Kanda, T., Tanaka, Y., Itaya, K., Okuno, M., Itoh, T., Iwai, N. and Wachi, M. (2022) Mining RNA-seq data reveals the massive regulon of GcvB small RNA and its physiological significance in maintaining amino acid homeostasis in *Escherichia coli*. Mol Microbiol, 117, 160–178.

87. Jørgensen, M.G., Nielsen, J.S., Boysen, A., Franch, T., Møller-Jensen, J. and Valentin-Hansen, P. (2012) Small regulatory RNAs control the multi-cellular adhesive lifestyle of *Escherichia coli*. Mol Microbiol, 84, 36–50.

88. Coornaert, A., Chiaruttini, C., Springer, M. and Guillier, M. (2013) Post-Transcriptional Control of the *Escherichia coli* PhoQ-PhoP Two-Component System by Multiple sRNAs Involves a Novel Pairing Region of GcvB. PLoS Genet, 9, e1003156.

89. Ju, X., Fang, X., Xiao, Y., Li, B., Shi, R., Wei, C. and You, C. (2021) Small RNA GcvB Regulates Oxidative Stress Response of *Escherichia coli*. Antioxidants, 10, 1774.

90. Opdyke, J.A., Kang, J.-G. and Storz, G. (2004) GadY, a Small-RNA Regulator of Acid Response Genes in *Escherichia coli*. J Bacteriol, 186, 6698–6705.

91. Melamed, S., Peer, A., Faigenbaum-Romm, R., Gatt, Y.E., Reiss, N., Bar, A., Altuvia, Y., Argaman, L. and Margalit, H. (2016) Global Mapping of Small RNA-Target Interactions in Bacteria. Mol Cell, 63, 884–897.

92. Melamed, S., Adams, P.P., Zhang, A., Zhang, H. and Storz, G. (2020) RNA-RNA Interactomes of ProQ and Hfq Reveal Overlapping and Competing Roles. Mol Cell, 77, 411–425.e7.

93. Iosub, I.A., van Nues, R.W., McKellar, S.W., Nieken, K.J., Marchioretto, M., Sy, B., Tree, J.J., Viero, G. and Granneman, S. (2020) Hfq CLASH uncovers sRNA-target interaction networks linked to nutrient availability adaptation. eLife, 9.

94. Freddolino, P.L. and Tavazoie, S. (2012) Beyond Homeostasis: A Predictive-Dynamic Framework for Understanding Cellular Behavior. Annu Rev Cell Dev Biol, 28, 363–384.

95. Hommais, F., Krin, E., Laurent-Winter, C., Soutourina, O., Malpertuy, A., Le Caer, J., Danchin, A. and Bertin, P. (2001) Large-scale monitoring of pleiotropic regulation of gene expression by the prokaryotic nucleoid-associated protein, H-NS. Mol Microbiol, 40, 20–36.

96. Janissen, R., Arens, M.M.A., Vtyurina, N.N., Rivai, Z., Sunday, N.D., Eslami-Mossallam, B., Gritsenko, A.A., Laan, L., de Ridder, D., Artsimovitch, I., et al. (2018) Global DNA Compaction in Stationary-Phase Bacteria Does Not Affect Transcription. Cell, 174, 1188–1199.e14.

97. Holmqvist, E. and Wagner, E.G.H. (2017) Impact of bacterial sRNAs in stress responses. Biochem Soc Trans, 45, 1203–1212.

98. Myers, K.S., Yan, H., Ong, I.M., Chung, D., Liang, K., Tran, F., Keleş, S., Landick, R. and Kiley, P.J. (2013) Genome-scale Analysis of Escherichia coli FNR Reveals Complex Features of Transcription Factor Binding. PLoS Genet, 9, e1003565.

99. Boysen, A., Møller-Jensen, J., Kallipolitis, B., Valentin-Hansen, P. and Overgaard, M. (2010) Translational Regulation of Gene Expression by an Anaerobically Induced Small Non-coding RNA in Escherichia coli. Journal of Biological Chemistry, 285, 10690–10702.

100. Stewart, V. (1993) Nitrate regulation of anaerobic respiratory gene expression in *Escherichia coli*. Mol Microbiol, 9, 425–434.

101. Brosse, A., Boudry, P., Walburger, A., Magalon, A. and Guillier, M. (2022) Synthesis of the NarP response regulator of nitrate respiration in *Escherichia coli* is regulated at multiple levels by Hfq and small RNAs. Nucleic Acids Res, 50, 6753–6768.

102. Groisman, E.A. (2001) The Pleiotropic Two-Component Regulatory System PhoP-PhoQ. J Bacteriol, 183, 1835–1842.

103. Eguchi, Y., Ishii, E., Hata, K. and Utsumi, R. (2011) Regulation of Acid Resistance by Connectors of Two-Component Signal Transduction Systems in *Escherichia coli*. J Bacteriol, 193, 1222– 1228.

104. Eguchi, Y., Itou, J., Yamane, M., Demizu, R., Yamato, F., Okada, A., Mori, H., Kato, A. and Utsumi, R. (2007) B1500, a small membrane protein, connects the two-component systems EvgS/EvgA and PhoQ/PhoP in *Escherichia coli*. Proceedings of the National Academy of Sciences, 104, 18712–18717.

105. Ishii, E., Eguchi, Y. and Utsumi, R. (2013) Mechanism of Activation of PhoQ/PhoP Two-Component Signal Transduction by SafA, an Auxiliary Protein of PhoQ Histidine Kinase in *Escherichia coli*. Biosci Biotechnol Biochem, 77, 814–819.

106. Vakulskas, C.A., Potts, A.H., Babitzke, P., Ahmer, B.M.M. and Romeo, T. (2015) Regulation of Bacterial Virulence by Csr (Rsm) Systems. Microbiology and Molecular Biology Reviews, 79, 193–224.

107. Brown, A.N., Anderson, M.T., Bachman, M.A. and Mobley, H.L.T. (2022) The ArcAB Two-Component System: Function in Metabolism, Redox Control, and Infection. Microbiology and Molecular Biology Reviews, 86.

108. Federowicz, S., Kim, D., Ebrahim, A., Lerman, J., Nagarajan, H., Cho, B., Zengler, K. and Palsson, B. (2014) Determining the Control Circuitry of Redox Metabolism at the Genome-Scale. PLoS Genet, 10, e1004264.

109. McClune, C.J., Alvarez-Buylla, A., Voigt, C.A. and Laub, M.T. (2019) Engineering orthogonal signalling pathways reveals the sparse occupancy of sequence space. Nature, 574, 702–706.

110. Sowa, S.W., Gelderman, G., Leistra, A.N., Buvanendiran, A., Lipp, S., Pitaktong, A., Vakulskas, C.A., Romeo, T., Baldea, M. and Contreras, L.M. (2017) Integrative FourD omics approach profiles the target network of the carbon storage regulatory system. Nucleic Acids Res, 10.1093/nar/gkx048.

111. Jiang, F., An, C., Bao, Y., Zhao, X., Jernigan, R.L., Lithio, A., Nettleton, D., Li, L., Wurtele, E.S., Nolan, L.K., et al. (2015) ArcA Controls Metabolism, Chemotaxis, and Motility Contributing to the Pathogenicity of Avian Pathogenic *Escherichia coli*. Infect Immun, 83, 3545–3554.

112. Alteri, C.J., Lindner, J.R., Reiss, D.J., Smith, S.N. and Mobley, H.L.T. (2011) The broadly conserved regulator PhoP links pathogen virulence and membrane potential in *Escherichia coli*. Mol Microbiol, 82, 145–163.

113. Tramonti, A., De Canio, M. and De Biase, D. (2008) GadX/GadW-dependent regulation of the *Escherichia coli* acid fitness island: transcriptional control at the *gadY–gadW* divergent promoters and identification of four novel 42bp GadX/GadW-specific binding sites. Mol Microbiol, 70, 965–982.

114. Seo, S.W., Kim, D., Szubin, R. and Palsson, B.O. (2015) Genome-wide Reconstruction of OxyR and SoxRS Transcriptional Regulatory Networks under Oxidative Stress in *Escherichia coli* K-12 MG1655. Cell Rep, 12, 1289–99.

