## Supplementary material for "Multiscale regulation of nutrient stress responses in *Escherichia coli* from chromatin structure to small regulatory RNAs": FileS1_SupplementaryFigures.pdf

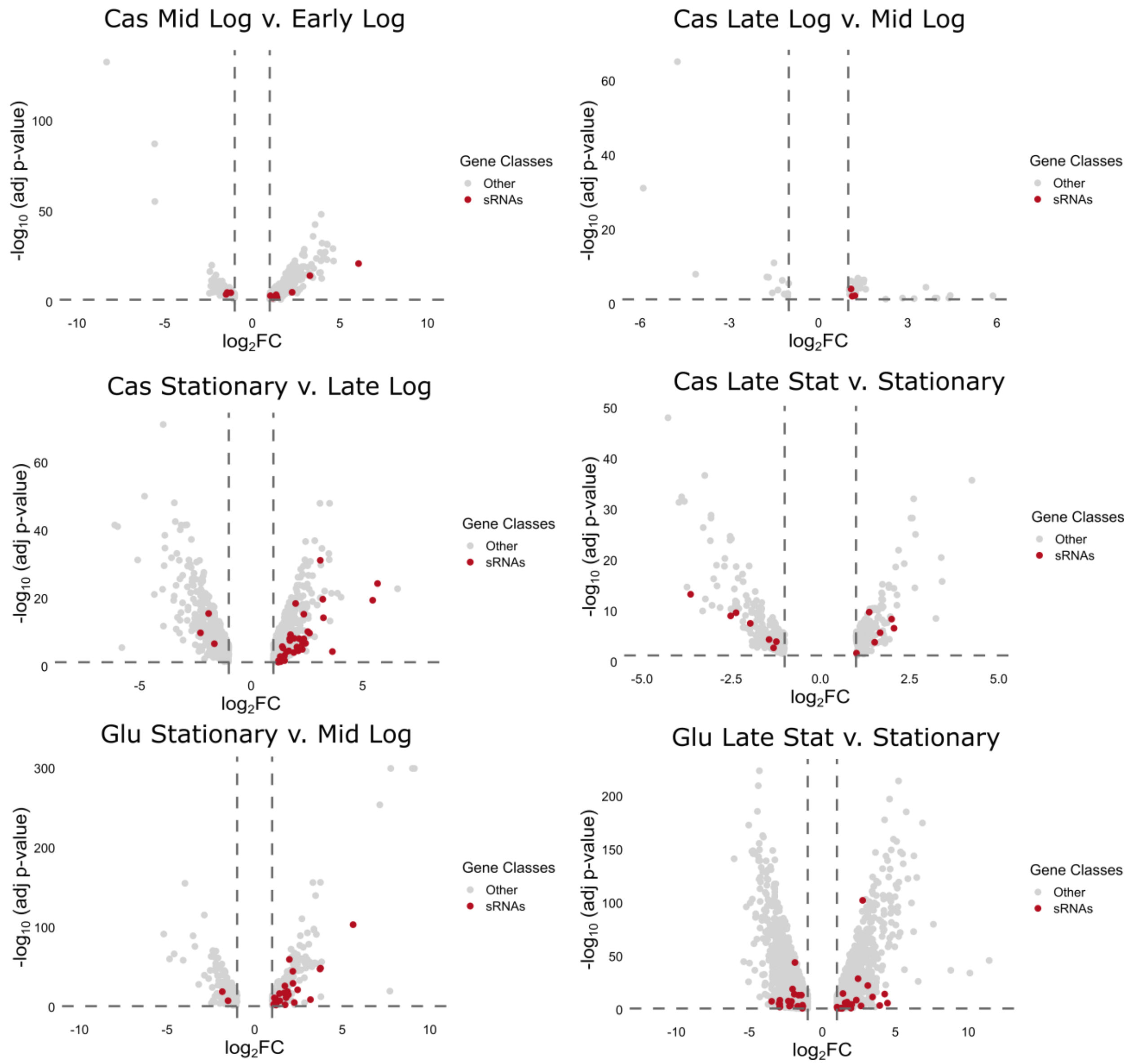


**Figure S1. Volcano plots of RNA-seq differential expression results across growth.** Differentially expressed genes (p-adj < 0.05, $log_2FC > |1|$) plotted for each neighboring growth phase in each media. sRNAs are indicated in red, all other genes in grey.

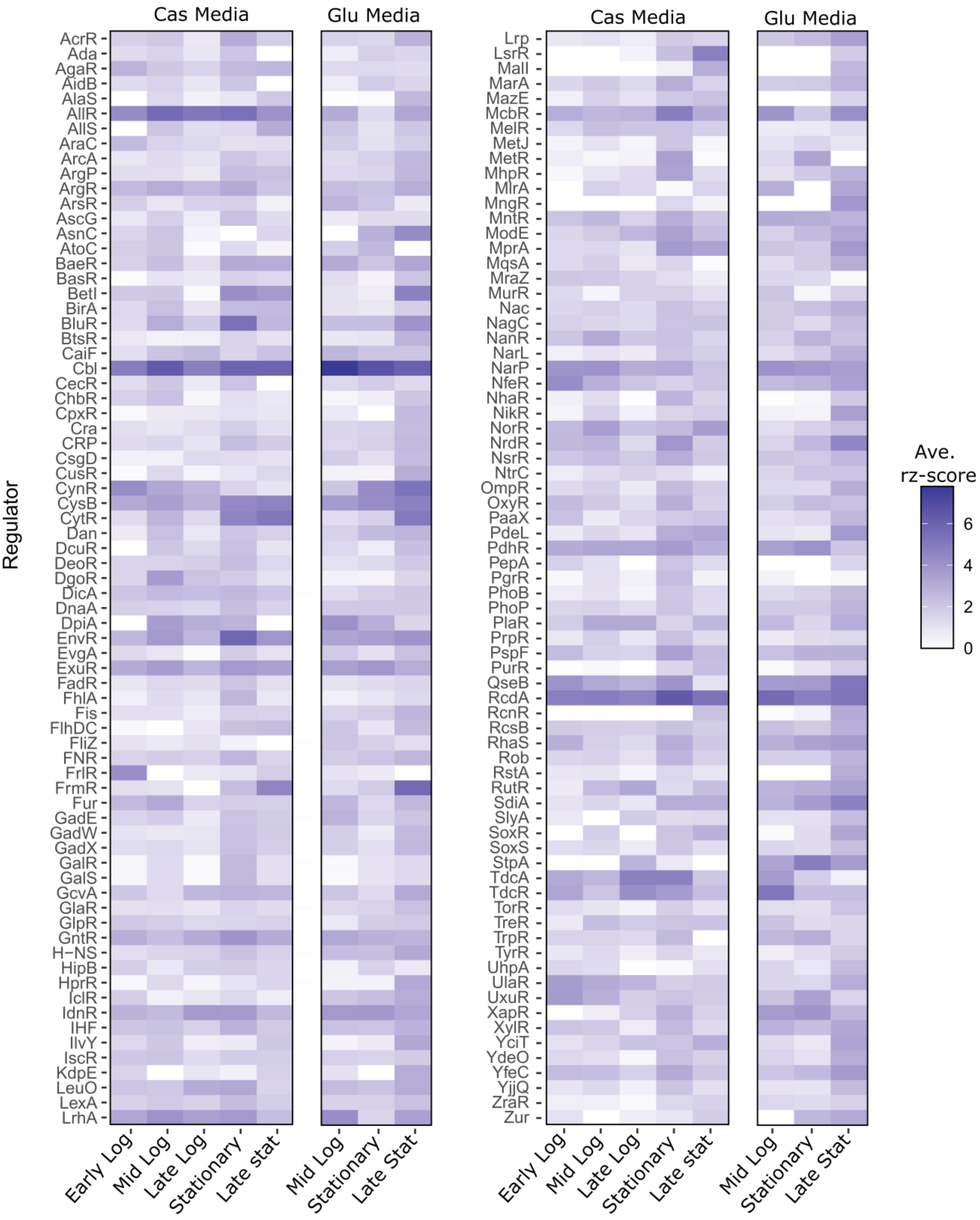


**Figure S2. Transcription factor regulon activities using protein occupancy**. The average raw z-score (rz-score) for all known binding sites (as reported by RegulonDB accessed March 1, 2024) for each regulator in the 8 conditions tested.

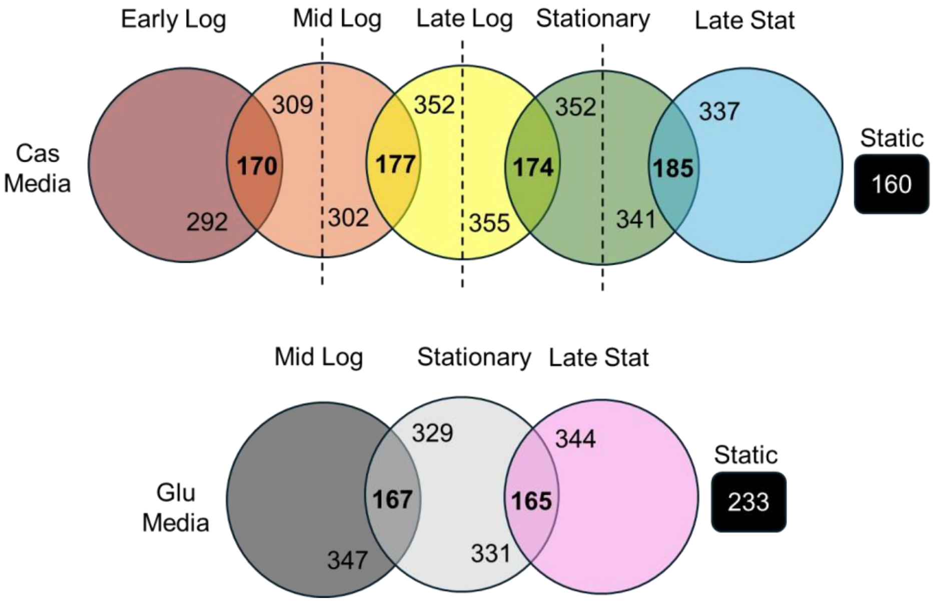


**Figure S3. Extended Protein Occupancy Domains (EPODs) conserved through growth**. The number of EPODs that are conserved (by location overlap comparison using bedtools intersect function) between each neighboring growth condition, in each media tested. Loose EPODs (75% cutoff) were compared between each growth phase separately, indicated by the dashed vertical lines, then filtered to only those that overlap with a strict EPOD (90% cutoff) in either of the two compared growth conditions. For example, early to mid logarithmic (log) phase in Cas medium had 170 shared loose EPOD locations that also make the strict EPOD threshold in early and/or mid logarithmic phases, with 292 and 309 unique (*i.e.*, do not overlap) or only loose (*i.e.*, do not achieve strict cutoff) EPODs in early log and mid log phase, respectively. Static EPODs refer to those loose EPOD locations that are present in all tested condition in that specified media and achieve the strict (90%) threshold in at least one compared condition.

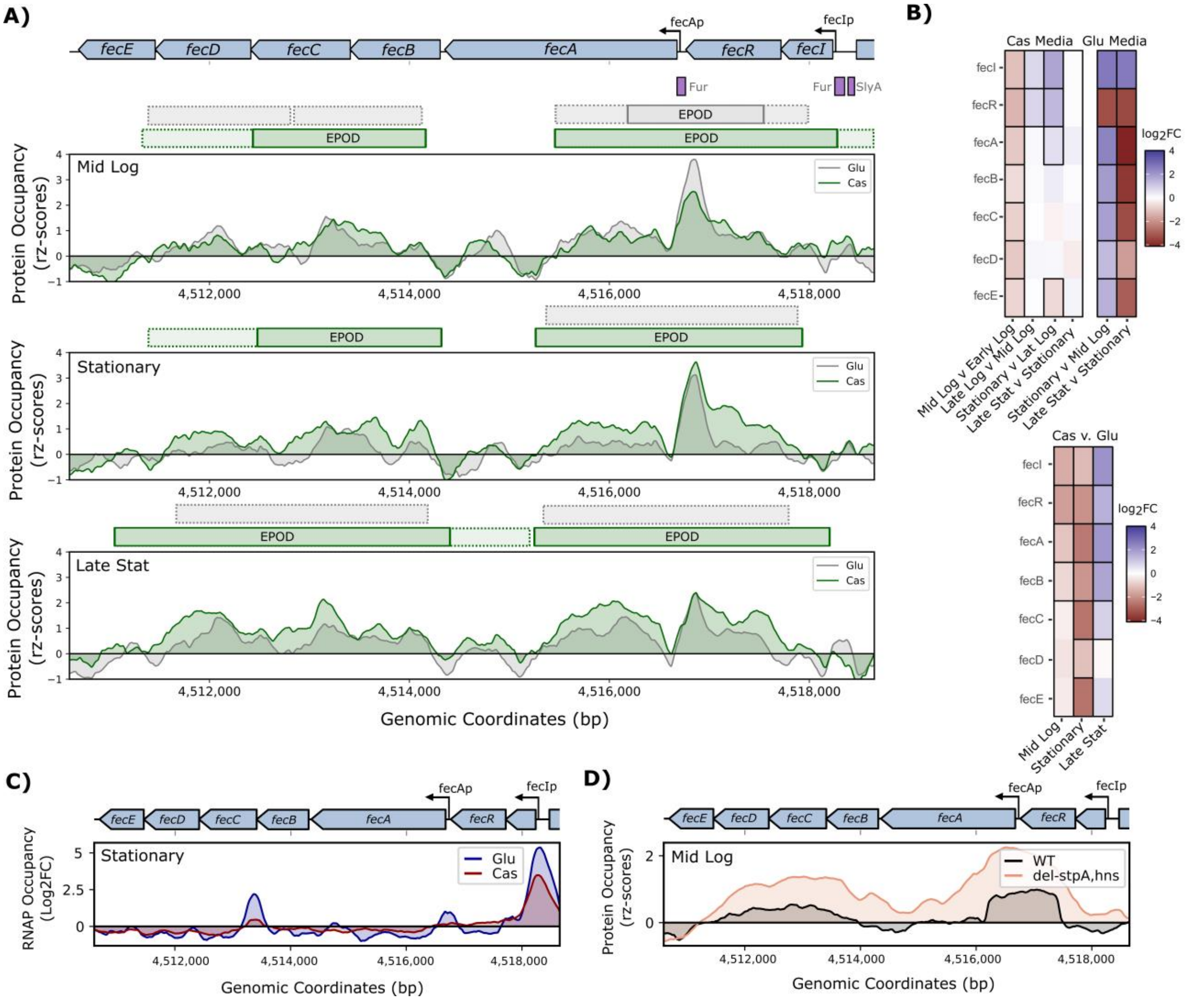


**Figure S4. Static EPOD in Cas medium example: iron regulatory *fecIRABCDE* operons. A)** Protein occupancy (rz-scores, moving average window of 250 bp) for mid logarithmic (top), stationary (middle) and late stationary phase (bottom) in Glu (grey) and Cas media (green) of the *fecIRABCDE* operons. Location of strict (solid) and loose (dashed) EPODs designated by the horizontal bars above each plot, color-coded by media. Note the Cas medium contains overlapping EPOD regions in all tested conditions, making two static EPODs, while Glu medium contains only one static EPOD designation for the downstream EPOD. The prominent peak upstream of *fecAp* is likely from the known Fur repressor binding sites (purple bars). **B)** Differential expression of the genes in the *fecIRABCDE* operons in Cas and Glu media (top) as well as between Cas vs. Glu media (bottom) in the three growth conditions shown in panel A. Significant differential expression is designated with black borders. Note the lack of dynamic changes in expression for genes in this operon across growth in Cas medium, when the static EPODs are present, and large shifts in expression in Glu medium when the strict EPODs are absent. However, comparing across media, while mid logarithmic and stationary match expectations (higher expression in Glu media without the strict EPODs designated by red), late stationary has higher expression in Cas media even with strict EPODs for *fecIR* and *fecABC* genes. **C)** RNAP occupancy ($log_2$FC compared to input, moving average window of 50 bp) at stationary phase for Glu (blue) and Cas (red) media. Note the high promoter activity outside of the strict EPODs compared to the internal cryptic promoters in the Cas medium. However, Glu media shows similar behavior with minimal strict EPODs. **D)** Protein occupancy (rz-scores) from a prior study (11) (NCBI GEO GSE164796) comparing a wildtype strain (grey) to nucleoid-associated protein deletion strain, Δ*stpA*Δ*hns*, is plotted (orange). None of the published nucleoid associated protein deletion strains tested in this dataset (Dps, IhfAB, H-NS, StpA, Fis, Hfq, HupAB) removed protein occupancy at these operons, suggesting a different protein composition.

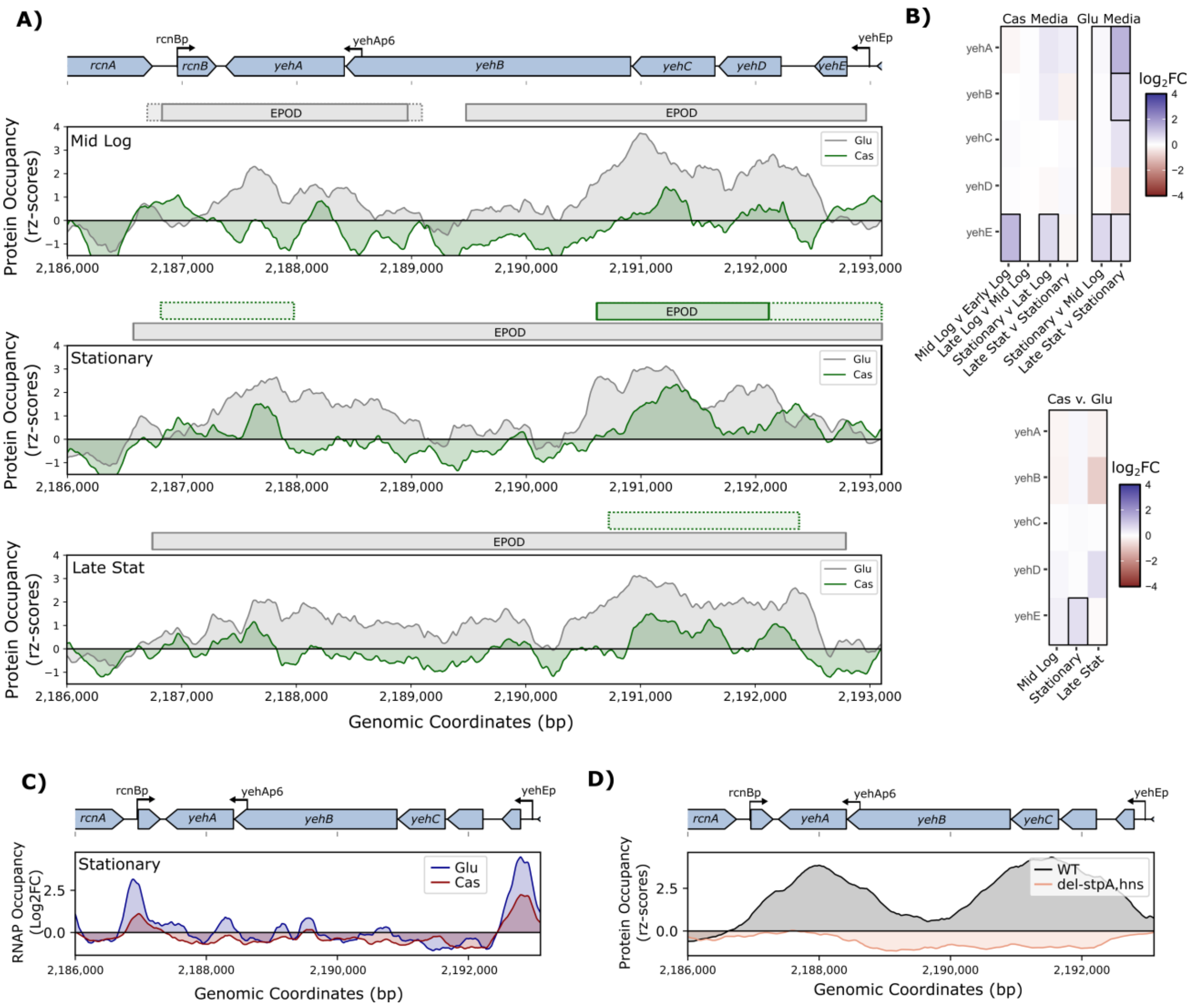


**Figure S5. Static EPOD in Glu medium example: cryptic flagellum *yehEDCBA* operon. A)** Protein occupancy (rz-scores, moving average window of 250 bp) for mid logarithmic (top), stationary (middle) and late stationary phase (bottom) in Glu (grey) and Cas media (green) of the *yehEDCBA* operon. Location of strict (solid) and loose (dashed) EPODs designated by the horizontal bars above each plot, color-coded by media. Note the Glu medium contains overlapping EPOD regions in all tested conditions, making it a static EPOD. **B)** Differential expression of the genes in the *yehEDCBA* operon in Cas and Glu media (top) as well as between Cas vs. Glu media (bottom) in the three growth conditions shown in panel A. Significant differential expression is designated with black borders. Note the lack of dynamic changes in expression for genes in this operon, regardless of the static EPOD being present (in Glu medium) or absent (in Cas medium). Likely transcription factors are also required to activate transcription of this operon when the EPOD is not present. **C)** RNAP occupancy ($log_2$FC compared to input, moving average window of 50 bp) at stationary phase for Glu (blue) and Cas (red) media of the *yehEDCBA* operon. Note the high promoter activity outside of the strict EPOD compared to the internal cryptic promoters in the Glu medium, and the lack of differences in the Cas medium even though the strict EPOD is absent. **D)** Protein occupancy (rz-scores) from a prior study (11) (NCBI GEO GSE164796) comparing a wildtype strain (grey) to nucleoid-associated protein deletion strains, in this case Δ*stpA*Δ*hns* is plotted (orange). Note the lack of protein occupancy when these two nucleoid-associated proteins are absent, supporting their contribution to this EPOD region.

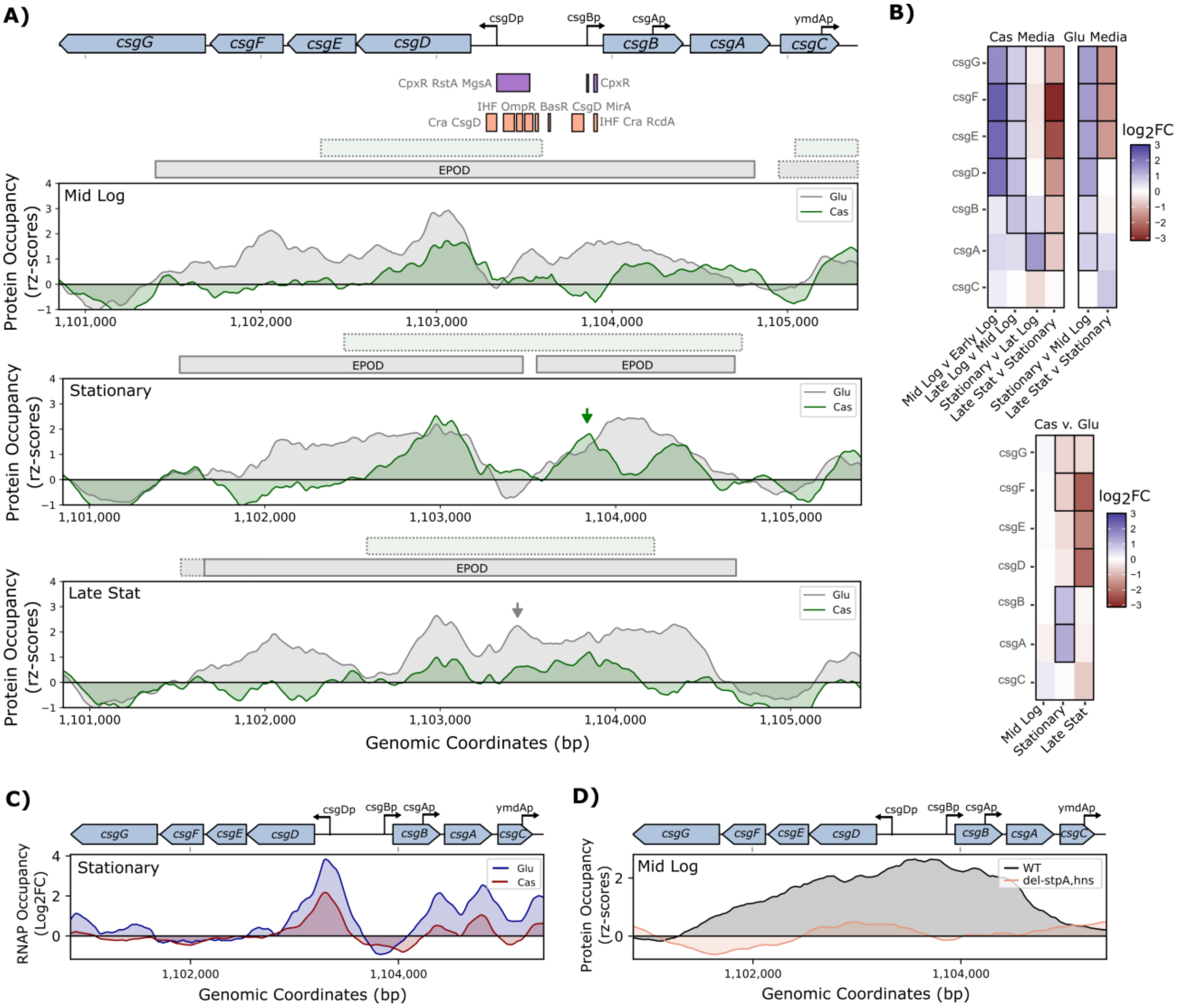


**Figure S6. Static EPOD with transcription factor coordination example: curli synthesis *csgBAC* and *csgDEFG* operons. A)** Protein occupancy (rz-scores, moving average window of 250 bp) for mid logarithmic (top), stationary (middle) and late stationary phase (bottom) in Glu (grey) and Cas media (green) of the *csgBAC* and *csgDEFG* operons. Location of strict (solid) and loose (dashed) EPODs designated by the horizontal bars above each plot, color-coded by media. Note the Glu media contains overlapping strict EPOD regions in all tested conditions, making it a static EPOD. It is worth noting that a small portion of this operon [1,102,605 – 1,103,450] is also a static EPOD in Cas media due to a strict EPOD call in the late logarithmic growth phase (not shown, Table S3). Well known binding sites of many transcription factors (including activators CsgD and Cra) are annotated by the arrows, color coded for the medium the peak occurs in. **B)** Differential expression of the genes in the *csgBAC* and *csgDEFG* operons in Cas and Glu media (top) as well as between Cas vs. Glu media (bottom) in the three growth conditions shown in panel A. Significant differential expression is designated with black borders. Note the differential expression of genes in the Cas vs. Glu in line with the appearance of the well characterized transcription factor binding sites noted by the purple (repressor) and orange (activator) horizontal boxes and protein occupancy peaks noted by arrows in panel A with binding sites. **C)** RNAP occupancy ($log_2$FC compared to input, moving average window of 50 bp) at stationary phase for Glu (blue) and Cas (red) media of the *csgBAC* and *csgDEFG* operons. **D)** Protein Occupancy (rz-scores) from a prior study (11) (NCBI GEO GSE164796) comparing a wildtype strain (grey) to nucleoid-associated protein deletion strains, in this case ΔstpAΔ*hns*, is plotted (orange). Note the lack of protein occupancy when these two nucleoid-associated proteins are absent, supporting their contribution to this EPOD region, with some minimal protein binding still present likely from other nucleoid-associated proteins and transcription factors.

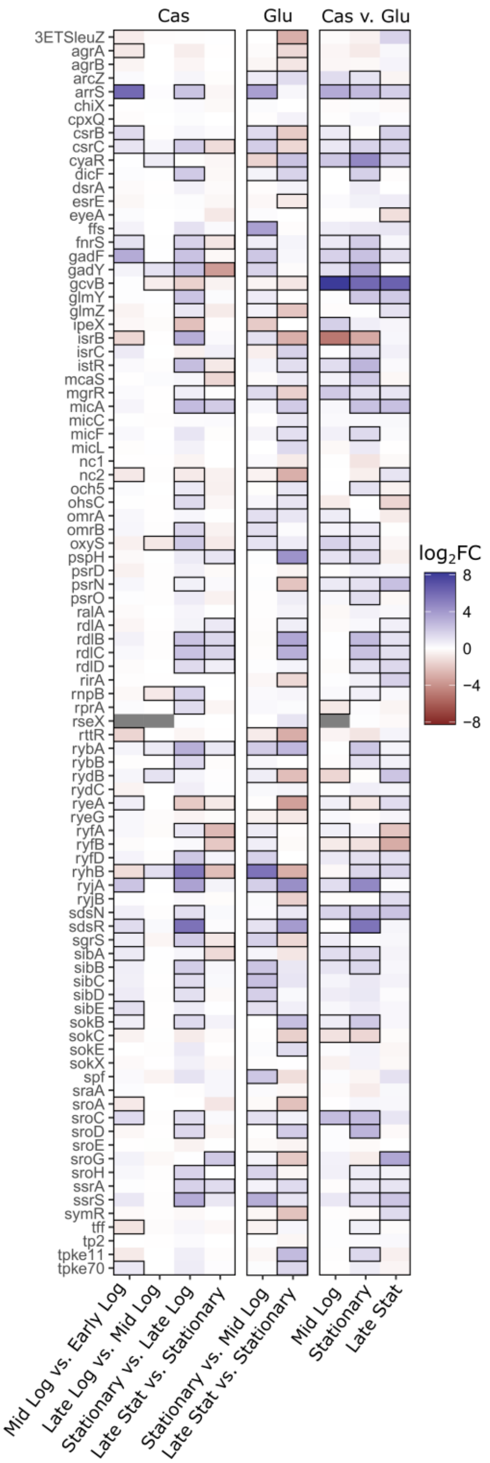


**Figure S7. Heatmap of sRNA differential expression between growth phases and media conditions.** Differential expression of the 91 sRNAs tested ($log_2FC$) over growth and between media conditions. Significant differential expression $log_2FC$ values (p-adj < 0.05) additionally annotated with black borders. The Glu and Cas media columns (left and middle) compare sRNA expression between neighboring growth stages, while the Cas vs. Glu media columns (right) compare between media at the three equivalent growth stages.

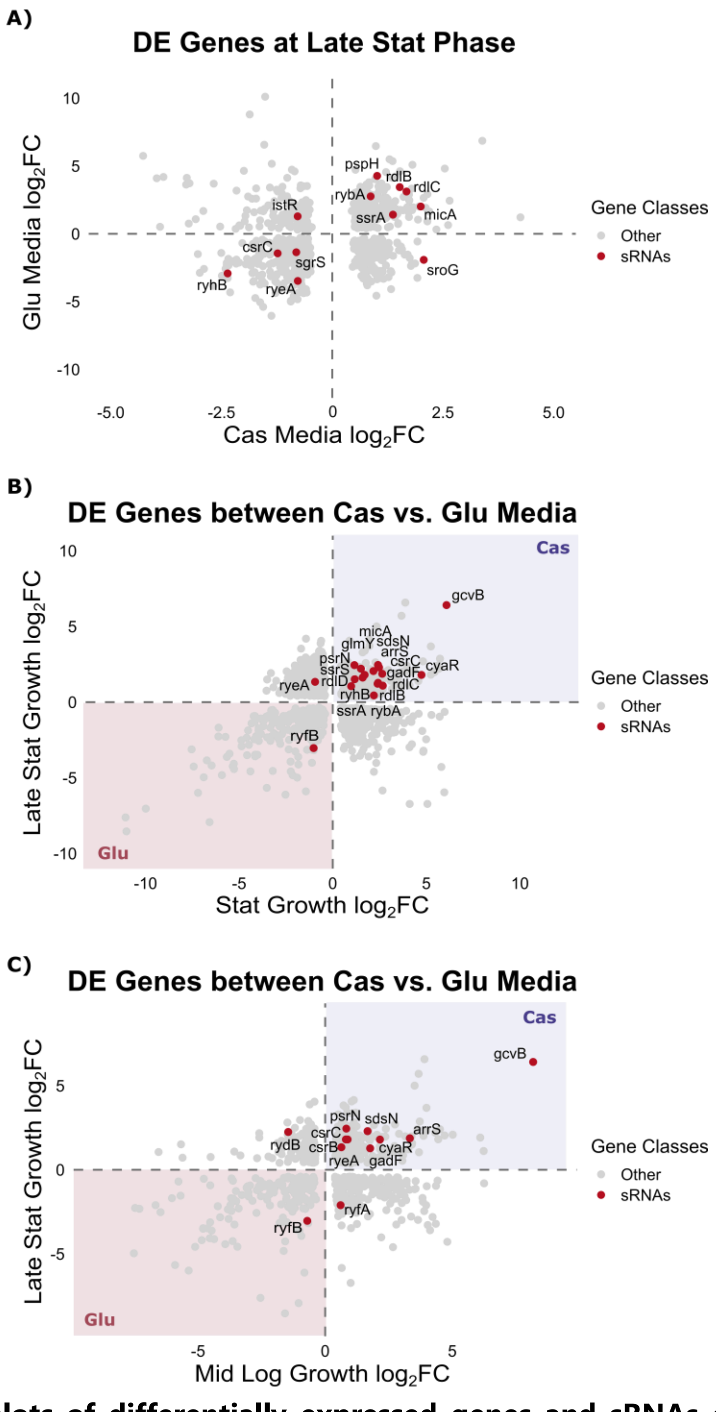


**Figure S8. Scatterplots of differentially expressed genes and sRNAs compared between media and growth conditions**. Differentially expressed genes in both plotted conditions (must be significant in both conditions to be plotted, p-adj < 0.05, $log_2FC > |1|$). sRNAs are shown in red. Conditions compared are **A)** Cas and Glu Media comparing late vs. stationary phase, **B)** stationary and late stationary phase comparing Cas vs. Glu media, and **C)** mid log and late stationary phase comparing Cas vs. Glu media.

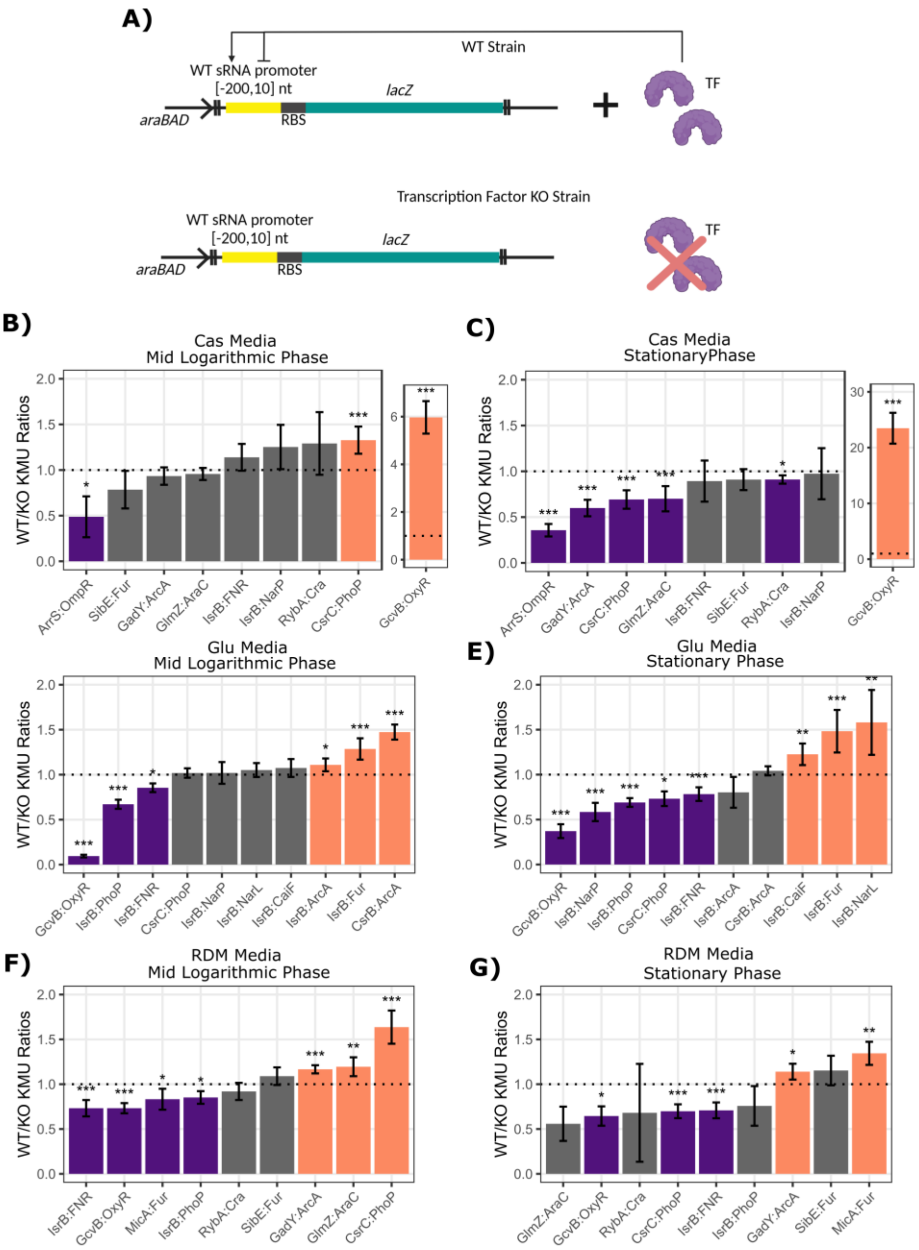


**Figure S9. Genomic Miller assays validate plasmid screening results and support direct transcription factors of sRNAs. A)** The wildtype (WT) promoter transcriptional reporter was genomically inserted at the ara operon in the wildtype (WT) and transcription factor (TF) knockout (KO) strain (similar to the plasmid-based screening experiments). **B-G)** The ratios of the wildtype promoter kinetic Miller Units (KMUs) in the wildtype to transcription factor knockout strain (WT/KO) at mid logarithmic and stationary phases, for Cas **(B-C)**, Glu **(D-E)** and RDM **(F-G)** media, respectively. Statistically significant ratios are colored in orange (activation, WT/Mut >1) and purple (repression, WT/Mut <1) and annotated with significance (from equal designated with dashed line, unpaired two-tailed t-test, each in biological quadruplicates, p-value: * < 0.05, ** <0.01, *** <0.005). Summarized results provided in **Table 2**.

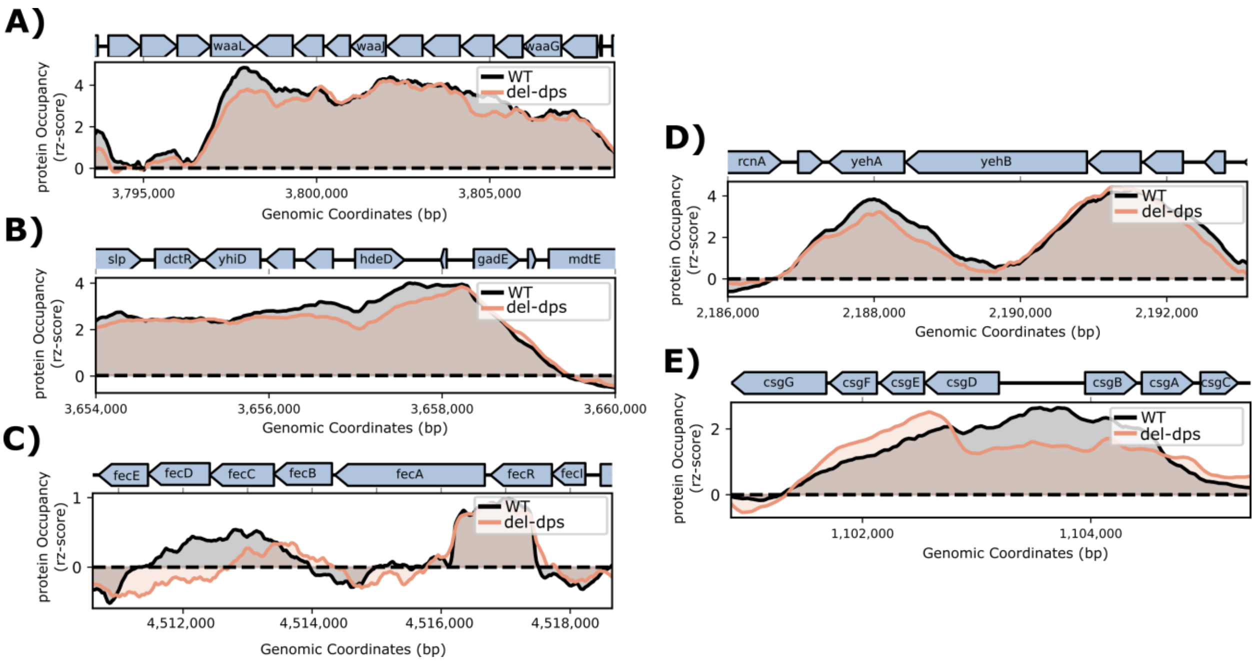


**Figure S10. EPOD examples in wildtype versus Δ*dps* strains from prior IPOD-HR study (11)**. The regions used as example EPODs in Figures 2D, 3D, and S4, S5, and S6, respectively, in which the IPOD-HR raw z-scores (rz-scores) from prior study (11) (NCBI GEO GSE164796) is shown for both the wildtype (WT, grey) strain and the Δ*dps* (KO, orange) strain. In all examples, deletion of the nucleoid associated protein, Dps, does not result in diminished protein occupancy. These results suggest other nucleoid-associated proteins independent of Dps are responsible for the EPODs. The rz-scores plotted are of a sliding window average over 250 bp.
